# Deciphering the role of adenosine-related genes in regulating smooth muscle cell phenotypic switch in coronary heart disease via single-cell RNA sequencing

**DOI:** 10.64898/2026.11.12.766320

**Authors:** Kaiyu Cao, Runze Wang, Yanling Qu, Xiaoyan Zhang, Zhonglin Xie, Xuehan Ma, Heping Ma, Ning Wang, Zhilu Li, Lin Cui, Xiyao Wang, Ke Su, Lijun Zhang

## Abstract

**Background:** Coronary heart disease (CHD) is closely associated with dysregulated lipid metabolism, inflammation, and impaired adenosine signaling. However, the contribution of adenosine-related genes (ARGs) and their relationship with the immune microenvironment remain unclear.

**Methods:** Public transcriptomic datasets were integrated with curated ARGs. Differential expression analysis and weighted gene co-expression network analysis identified candidate genes, and XGBoost, support vector machine-recursive feature elimination, and Boruta were used to select diagnostic features. Gene set enrichment, immune infiltration, regulatory-network, and single-cell RNA-sequencing analyses were performed, followed by reverse transcription-quantitative polymerase chain reaction validation.

**Results:** We identified 4, 884 differentially expressed genes and 243 ARG-associated module genes, yielding 231 candidate genes. DYRK2 and SIGIRR were prioritized as key genes and were consistently downregulated in bulk CHD datasets, including the validation cohort. A two-gene nomogram showed strong diagnostic performance. Functional analyses linked both genes to mitochondrial dysfunction and innate immune regulation, while immune profiling indicated reduced adaptive immune populations and enhanced myeloid-associated inflammation. Single-cell analysis identified smooth muscle cells as a key compartment with altered DYRK2 and SIGIRR expression along pseudotime trajectories and remodeling of vascular endothelial growth factor and pleiotrophin signaling. Reverse transcription-quantitative polymerase chain reaction in clinical samples (n = 5 per group) supported the bioinformatic expression trends (P < 0.05).

**Conclusions:** DYRK2 and SIGIRR emerged as key adenosine-related genes associated with immune-metabolic imbalance and smooth muscle cell dysfunction in CHD. These findings provide a basis for further mechanistic and translational validation.

## 1 Introduction

Coronary heart disease (CHD) stands as the most prevalent cardiovascular disease worldwide, with a multifaceted pathogenesis encompassing lipid metabolic disturbances, chronic inflammatory responses, and a range of other pathological processes^1^. A growing body of evidence has established that adenosine plays an indispensable role in the onset and progression of CHD ^2, 3^. As an endogenous purine nucleotide, adenosine exerts its vascular effects primarily through specific cell surface receptors on coronary vascular smooth muscle and endothelial cells, influencing CHD progression by regulating vascular tone, platelet activity, and microcirculation function^2^. In CHD, the imbalance in adenosine metabolism leads to aberrant signaling pathways of receptors such as A2AR, consequently affecting downstream targets (e.g., H₂O₂ and K⁺ channels) and ultimately impairing coronary arterial diastolic function ^2^. Furthermore, adenosine modulates immune responses through antagonistic actions on A2A/A2B receptors ^4^.

In recent years, the development of single-cell RNA sequencing (scRNA-seq) has offered novel perspectives on the cellular diversity and molecular pathways associated with CHD. Unlike bulk RNA sequencing, scRNA-seq enables the identification of cell-type-specific gene expression patterns and cell-cell interactions, offering a powerful tool to dissect the complex pathophysiology of CHD at single-cell resolution. Despite these advances, several critical questions remain unanswered. First, the specific ARGs that orchestrate CHD pathogenesis at the cellular level have not been systematically identified. Second, the crosstalk between ARG-mediated signaling and immune dysregulation in CHD remains poorly characterized. Third, how ARGs influence smooth muscle cell (SMC) phenotypic switching—a hallmark of atherosclerotic progression—has not been elucidated at single-cell resolution. These knowledge gaps hinder the development of adenosine-targeted therapeutic strategies for CHD. To bridge these knowledge gaps, this study conducted a systematic investigation into the function of ARGs in CHD pathogenesis, especially regarding their influence on immune dysregulation and SMC phenotypic transition. Transcriptomic data derived from peripheral blood mononuclear cells of CHD patients were integrated with a compiled ARG dataset obtained from public databases. Potential diagnostic genes were screened via differential expression analysis, WGCNA, and three machine learning methods: XGBoost, SVM-RFE, and Boruta. The expression patterns of key genes across different cell types were subsequently verified using scRNA-seq, and their roles in vascular SMC differentiation and intercellular signaling were systematically characterized. Computational drug prediction was further employed to identify potential therapeutic agents targeting these genes. This comprehensive, multi-dimensional analytical framework is anticipated to uncover novel adenosine-related mechanisms in CHD and offer clinically translatable insights for early diagnosis and precision medicine.

## 2 Methods

### 2.1 Data acquisition

The training set (GSE113079) was downloaded from the Gene Expression Omnibus (GEO) database, encompassing 141 peripheral blood mononuclear cell samples (93 CHD and 48 normal controls) profiled on the GPL20115 platform. The validation set (GSE42148), also retrieved from GEO, comprised 24 peripheral blood samples (13 CHD and 11 normal controls) assayed on the GPL1360 platform. scRNA-seq data (GSE121893, platform GPL18573) included tissue samples from 2 CHD patients and 2 normal controls; 4 dilated cardiomyopathy samples were excluded. A total of 22 ARGs were curated from published literature ^5^ **(Tables S1)**.

### 2.2 Differential expression analysis

DEGs between CHD and control samples in the training set (GSE113079) were identified using the R package “limma” (v 3.58.1) ^6^, with thresholds of P < 0.05 and |log₂FC| > 0.5. Distribution of DEGs was visualized as a volcano plot with “ggplot2” (v 3.5.1) ^7^, and expression patterns of the top 10 up- and downregulated genes (ranked by |log₂FC|) were displayed as a heatmap using “ComplexHeatmap” (v 2.21.1) ^8^.

### 2.3 WGCNA

ARG enrichment scores across training set samples were quantified via ssGSEA using the R package “GSVA” (v 1.50.0) ^9^. WGCNA was then performed using the R package “WGCNA” (v 1.72.5) ^10^. After outlier removal via the goodSamplesGenes function, a soft-thresholding power was selected where the scale-free topology fit index (R²) reached 0.9 and mean connectivity approached zero (powers 1–20). Co-expression modules were defined by dynamic tree cutting (minimum module size: 200 genes; merging cut height: 0.25). Module eigengenes (MEs) were calculated, and Spearman correlations between MEs and ARG scores were visualized as a heatmap using “pheatmap” (v 1.0.12)^11^. Modules with the strongest positive and negative correlations were selected, and genes within these modules meeting |GS| > 0.5 and |MM| > 0.7 ^12^ were retained as ARG-related module genes.

### 2.4 Identification of candidate gene and functional characteristics

Candidate genes at the intersection of ARG-related module genes and DEGs were identified using “ggvenn” (v 0.1.10) ^13^ GO and KEGG enrichment analyses were conducted via “clusterProfiler” (v 4.8.3) ^14^ (P < 0.05), covering BP, CC, and MF domains. A PPI network was constructed using the STRING database (https://string-db.org/; Homo sapiens; confidence score ≥ 0.4) and visualized with “circlize” (v 0.4.16) ^15^.

### 2.5 Machine learning

Three machine learning algorithms were applied to prioritize feature genes. XGBoost was implemented via the R package “xgboost” (v 2.0.3.1) ^16^. with 5-fold cross-validation and grid search optimization. SVM-RFE was performed using “caret” (v 6.0.94) ^17^ with 5-fold cross-validation, retaining the minimal feature subset yielding the highest accuracy. The Boruta algorithm, executed via “Boruta” (v 8.0.0) ^18^, permuted original features to construct shadow distributions and labeled genes with significantly higher Z-scores as “Confirmed.” Final feature genes were defined as the intersection of results from all three algorithms.

### 2.6 ROC curve and expression analysis

Diagnostic performance of each feature gene was assessed by ROC analysis using “pROC” (v 1.18.5) ^19^ in both training and validation sets; genes with AUC > 0.7 and AUC ≠ 1 in both sets were retained. Wilcoxon tests were used to compare expression between CHD and controls in each dataset, and genes with consistent, statistically significant differences (P < 0.05) across both sets were designated as key genes. Results were visualized with “ggplot2” (v 3.5.1).

### 2.7 Construction of a nomogram

A nomogram for CHD prediction based on key genes was developed using “rms” (v 6.8.1) ^20^. Calibration was evaluated by calibration curve and the Hosmer-Lemeshow test (P > 0.05 indicating adequate fit). Overall diagnostic accuracy was assessed by ROC analysis (“pROC” v 1.18.5; AUC > 0.7), and clinical utility was evaluated by DCA using “ggDCA” (v 1.2) ^21^.

### 2.8 Gene set enrichment analysis (GSEA) and functional network

GSEA was conducted to identify pathways associated with key genes using the gene set “c2.cp.kegg_legacy.v2025.1.Hs.symbols.gmt” (MSigDB) as reference. Genes were ranked by Spearman correlation with each key gene, computed via “psych” (v 2.4.3) ^22^, and analyzed through “clusterProfiler” (v 4.8.3). Significantly enriched pathways were defined by |NES| > 1, adjusted P < 0.05, and FDR < 0.25. The top five pathways per key gene were visualized with “GseaVis” (v 0.1.0) ^23^, and commonly enriched pathways were displayed as a heatmap using “pheatmap” (v 1.0.12). Functional interactions were further explored via GeneMANIA (http://www.genemania.org/).

### 2.9 Immune cell infiltration

Immune cell infiltration was estimated by ssGSEA ^24^. across all training set samples. Differences in infiltration between CHD and controls were assessed by Wilcoxon tests (P < 0.05). Spearman correlation analysis using “psych” (v 2.4.3) was applied to examine associations among differentially infiltrated immune cells and between key genes and these cells (|cor| > 0.3, P < 0.05).

### 2.10 Location and regulatory network

Chromosomal locations of key genes were mapped with “RCircos” (v 1.2.2) ^25^. Targeting miRNAs were predicted via miRDB (https://mirdb.org/; accessed September 4, 2025; target score > 80), and associated lncRNAs were identified through starBase (https://rnasysu.com/encori/; accessed September 4, 2025; human species; clipExpNum > 10). The top 20 mRNA–miRNA–lncRNA pairs per key gene, ranked by clipExpNum, were selected for visualization.

### 2.11 Identification of potential therapeutic drugs

Potential therapeutic drugs targeting the key genes were predicted using the R package “enrichR” (v 3.2) ^26^, leveraging the DsigDB (https://dsigdb.tanlab.org/) with a significance threshold of P < 0.05. Drug-gene relationships were visualized as a network plot using the R package “ggraph” (v 2.2.1) ^27^.

### 2.12 Data of scRNA-seq preprocessing and cell type annotation

scRNA-seq data were preprocessed using “Seurat” (v 5.1.0) ^28^. Genes expressed in fewer than 3 cells were removed. Retained cells met the following criteria: total counts 200–30, 000, detected genes 200–4, 000, and mitochondrial content ≤ 5%. The top 2, 000 HVGs were identified using the vst method and labeled with LabelPoints. Expression data were normalized via ScaleData, and PCA was performed. Significant principal components were determined by JackStrawPlot. Cell clusters were defined using FindNeighbors and FindClusters (resolution = 0.9), and UMAP was applied for visualization. Clusters were annotated by integrating marker gene profiles with published literature ^29^ and the CellMarker database.

### 2.13 Identification of key cell types

Cell type proportions across samples were calculated and visualized with “ggplot2” (v 3.5.1). Differences between CHD and controls were assessed by Wilcoxon tests (P < 0.05). Key gene expression within each differentially abundant cell type was evaluated by Wilcoxon tests (P < 0.05). Cell types showing both significant abundance changes and significant key gene expression alterations were designated as key cell types.

### 2.14 Pseudotime trajectory analysis of key cell cluster types

Expression data of key cell types were standardized using ScaleData, followed by PCA with the optimal number of PCs determined by ElbowPlot (Figure S2A). Cells were sub-clustered, and subtype marker genes were identified by FindAllMarkers (min.pct = 0.25, logfc.threshold = 1, test.use = “wilcox”). Top 5 markers per subtype were visualized as a heatmap using “pheatmap” (v 1.0.12). Pseudotime analysis via “monocle” (v 2.28.0)^30^ was used to reconstruct differentiation trajectories and characterize key gene expression dynamics during state transitions.

### 2.15 Pathway enrichment and cell-cell communication

Pathway enrichment of key cell types in CHD was analyzed using “ReactomeGSA” (v 1.20.0) ^31^, with the top 10 significant pathways visualized. Cell–cell communication was assessed with “CellChat” (v 1.6.1) ^32^ by comparing signaling networks between CHD and controls, focusing on key ligand–receptor interactions within the VEGF and PTN pathways.

### 2.16 Reverse transcription-quantitative polymerase chain reaction (RT-qPCR)

Samples from CHD patients (n = 5) and controls (n = 5) were collected at the Department of Cardiology, The 941th Hospital of Joint Logistic Support Force of Chinese People’s Liberation Army. All participants provided written informed consent prior to enrollment. This study was approved by the Ethics Committee of The 941th Hospital of Joint Logistic Support Force of Chinese People’s Liberation Army (Approval No.: [2026022]). Total RNA was isolated from samples using TRIzol reagent (Vazyme Biotech, cat. no. R401-01, Nanjing, China) following the protocol of the manufacturer. RT-qPCR was performed using Servicebio® 2× Universal Blue SYBR Green qPCR Master Mix (Kunming Yungen Biotechnology Co., Ltd., cat. no. 24Y0028, Kunming, China) on a CFX Connect Real-Time PCR System (Bio-Rad Laboratories, cat. no. XLFZ006, Hercules, CA, USA). Primer sequences were shown in **Table 1**. All reactions were implemented in technical triplicates, and Ct values were standardized against GAPDH using the 2^−ΔΔCt^ method. Statistical analyses were conducted using GraphPad Prism 10 software^33^. A t-test was employed to compare the expression differences between the two groups (P < 0.05).

**Table 1.**
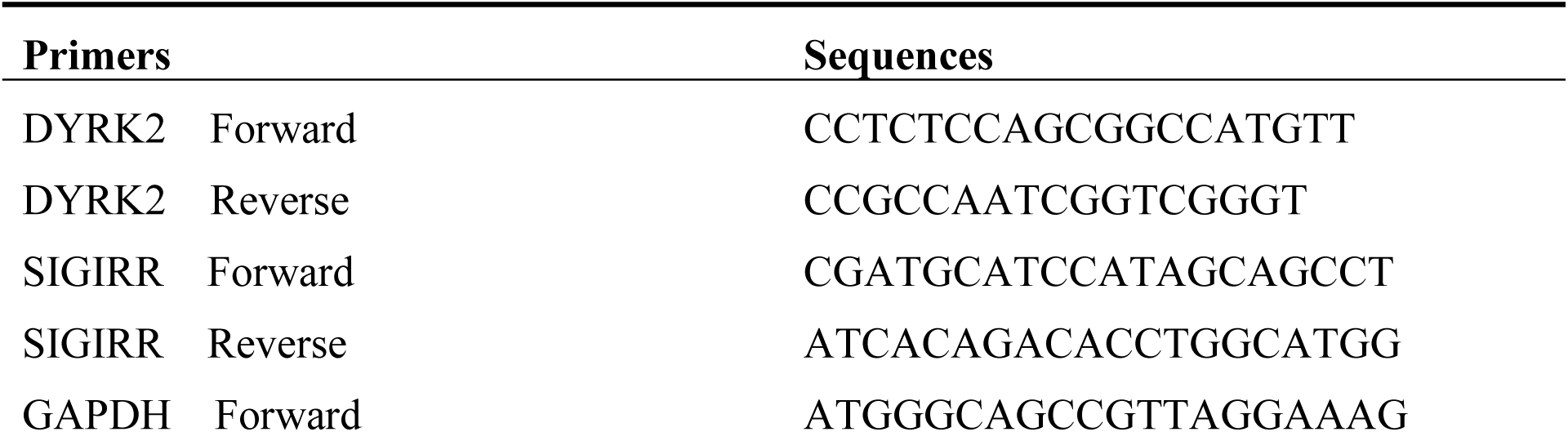
Primer Sequences Used for RT-qPCR.

### 2.17 Statistical analysis

All analyses were conducted in R (v 4.3.1). Between-group comparisons were performed using the Wilcoxon test (P < 0.05).

## 3 Results

### 3.1 Identification of candidate genes and functional characteristics

Differential expression analysis revealed 4, 884 DEGs (P < 0.05, |log₂FC| > 0.5) between CHD samples and controls, including 2, 773 upregulated and 2, 111 downregulated genes (**Figure 1A**). ARG scores were significantly elevated in CHD samples relative to controls (P < 0.001) (**Figure 1B**). All samples passed quality control and were retained for downstream analyses. WGCNA with a soft-thresholding power of 7 yielded nine co-expression modules (including the grey module for unassigned genes) (**Figure 1C**). The blue module showed the most pronounced positive correlation with ARG scores, whereas the brown module exhibited the strongest negative correlation (**Figure 1D**). A total of 243 ARG-related module genes were obtained (155 from the blue module; 88 from the brown module), of which 231 overlapped with the DEG list and were defined as candidate genes (**Figure 1E**). GO enrichment analysis identified significant associations across 110 BP terms (e.g., rRNA processing, rRNA metabolic process, regulation and positive regulation of dendrite extension, and maturation of 5.8S rRNA), 18 CC terms (e.g., exosome (RNase complex), keratin filament, postsynaptic specialization, nuclear exosome (RNase complex), and GABAergic synapse), and 18 MF terms (e.g., protein−cysteine S−acyltransferase activity, protein−cysteine S−palmitoyltransferase activity, TBP-class protein binding, calcium-dependent phospholipid binding, sphingolipid transporter activity, and poly(A) RNA polymerase activity). KEGG analysis identified enrichment in 19 pathways, including cholinergic synapse, serotonergic synapse, long-term depression, and the hedgehog signaling pathway (P < 0.05) (**Figure 1F, Supplementary Tables 2–3)**. PPI network analysis of the 231 candidate genes retained 74 interacting nodes after excluding 157 isolated genes, with RPL7L1, CSNK1A1, and CDH2 identified as prominent hub genes (**Figure 1G**). Taken together, these results suggest that candidate genes may participate in CHD pathogenesis via rRNA processing, synaptic signaling, and lipid metabolism.

**Figure 1.**
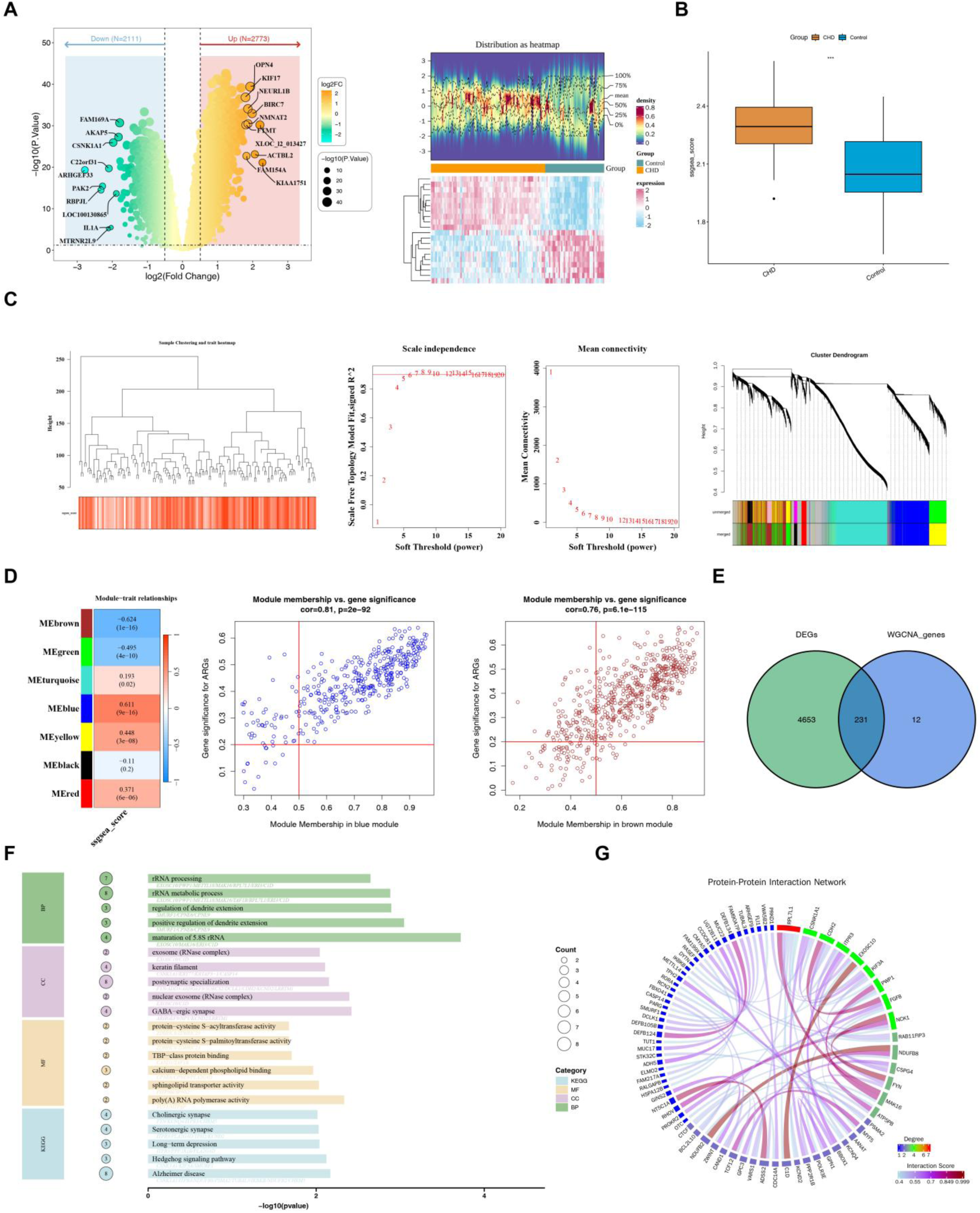
Identification of candidate genes and functional characteristics. (A) Volcano plot and heatmap depicting differentially expressed genes (DEGs) between coronary heart disease (CHD) samples and controls in the training set (GSE113079). A total of 4, 884 DEGs were identified (P < 0.05, |log₂FC| > 0.5). (B) Comparison of adenosine-related gene (ARG) scores between CHD samples and controls, showing significantly elevated scores in CHD (P < 0.001). (C) Dendrogram from weighted gene co-expression network analysis (WGCNA) illustrating nine co-expression modules using a soft-threshold of 7. (D) Correlations between module eigengenes and ARG scores, with the blue module showing the strongest positive correlation and the brown module the strongest negative correlation. (E) Venn diagram representing the intersection of ARG-related module genes and DEGs, yielding 231 candidate genes. (F) Bar plots of Gene Ontology (GO) and Kyoto Encyclopedia of Genes and Genomes (KEGG) enrichment analyses for candidate genes, highlighting top-enriched terms. (G) Protein-protein interaction (PPI) network of candidate genes, with nodes like RPL7L1 and CDH2 exhibiting extensive interactions. Statistical significance symbols: *P < 0.05; **P < 0.01; ***P < 0.001; ****P < 0.0001; ns, not significant.

### 3.2 Machine learning-based identification and validation of key genes

To identify key genes for CHD, three machine learning algorithms were employed on 231 candidate genes. The XGBoost method selected 68 feature genes, including C10orf118, RCN2, and ZBTB11 (**Figure 2A**). SVM-RFE achieved its highest accuracy (0.9073) and lowest error (0.00927), identifying 36 feature genes such as RCN2, C10orf118, and AGPAT4 (**Figure 2B**). Boruta analysis produced 32 feature genes, including GPN1, AGPAT4, and C10orf118 (**Figure 2C**). The intersection of these three gene sets yielded 17 common feature genes (**Figure 2D**). ROC curve analysis indicated that six genes—AGPAT4, DYRK2, GPN1, RCN2, SIGIRR, and ZBTB11—displayed AUC values greater than 0.7 in both the training and validation sets (**Figure 2E**). Expression analysis in the training set revealed that all six genes were significantly downregulated in CHD samples (P < 0.0001). In the validation set, only DYRK2 and SIGIRR remained significantly downregulated in CHD samples (P < 0.05), while the others showed no statistically significant difference (**Figure 2F**). These findings highlight DYRK2 and SIGIRR as consistently dysregulated genes across independent cohorts, implying their potential involvement in CHD pathogenesis.

**Figure 2.**
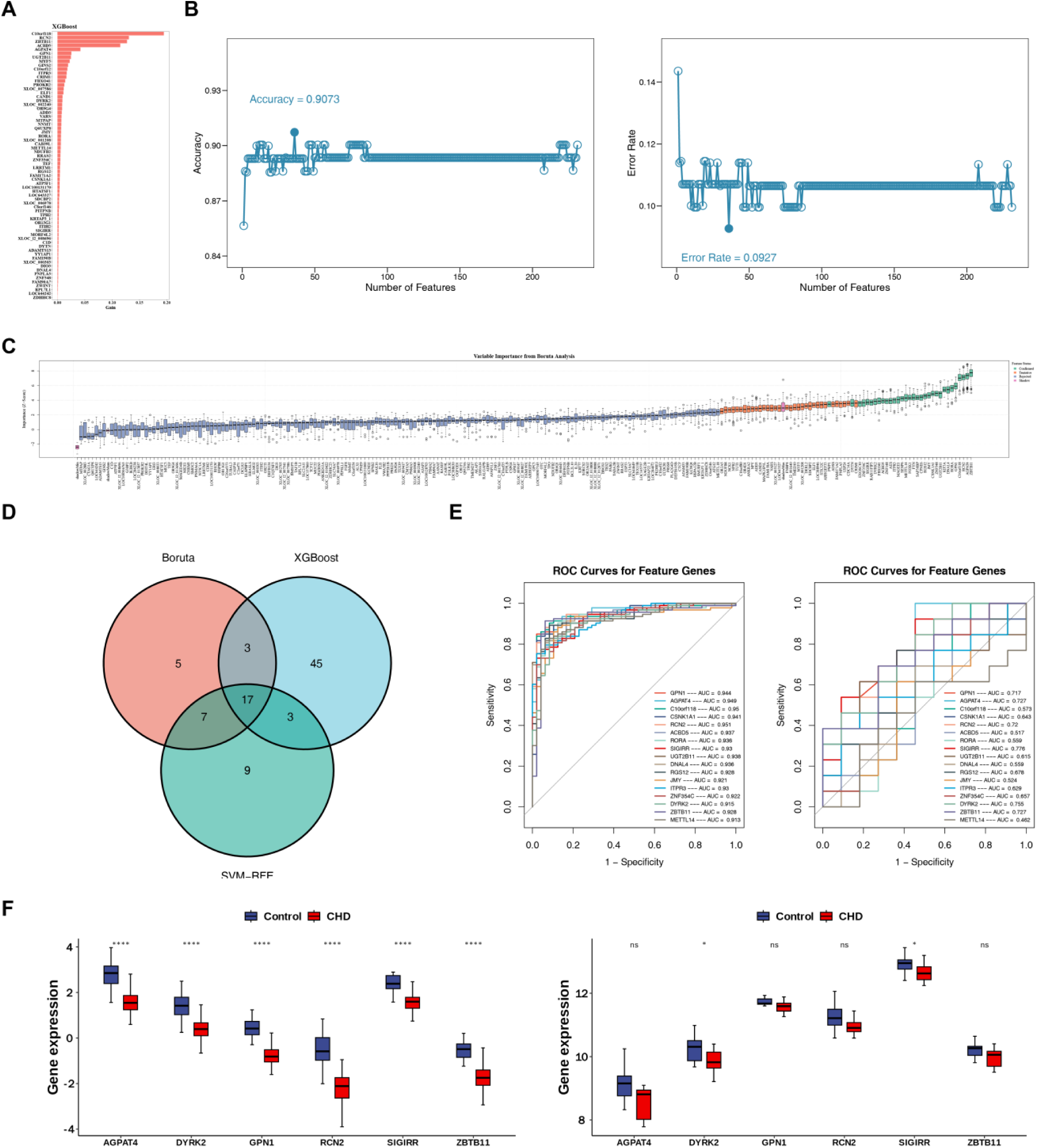
Machine learning-based identification and validation of key genes. (A) Feature importance plot from XGBoost analysis, identifying 68 feature genes. (B) Accuracy plot from support vector machine-recursive feature elimination (SVM-RFE). (C) Feature confirmation plot from Boruta analysis, designating 32 “Confirmed” feature genes. (D) Venn diagram showing the intersection of feature genes from all three algorithms, resulting in 17 genes. (E) Receiver operating characteristic (ROC) curves (e.g., AGPAT4, DYRK2) in training and validation sets, with area under the curve (AUC) values. (F) Box plots comparing expression levels of key genes between CHD samples and controls in training and validation sets, highlighting significant downregulation of DYRK2 and SIGIRR in both cohorts (P < 0.05). Statistical significance symbols: *P < 0.05; **P < 0.01; ***P < 0.001; ****P < 0.0001; ns, not significant.

### 3.3 Construction and evaluation of a diagnostic nomogram

A nomogram integrating DYRK2 and SIGIRR expression was developed to quantify CHD risk. Higher total points correlated with an increased probability of CHD. For example, a total point of 129 corresponded to a 97.8% risk of CHD occurrence (**Figure 3A**). The model exhibited excellent calibration, with the actual calibration curve closely aligning with the ideal line (Hosmer-Lemeshow test, P = 0.66) (**Figure 3B**). ROC analysis confirmed high diagnostic accuracy (AUC = 0.94) (**Figure 3C**). DCA further demonstrated superior net benefit compared to single-gene models across threshold probabilities (**Figure 3D**). These results established a nomogram as a robust tool for CHD risk assessment.

**Figure 3.**
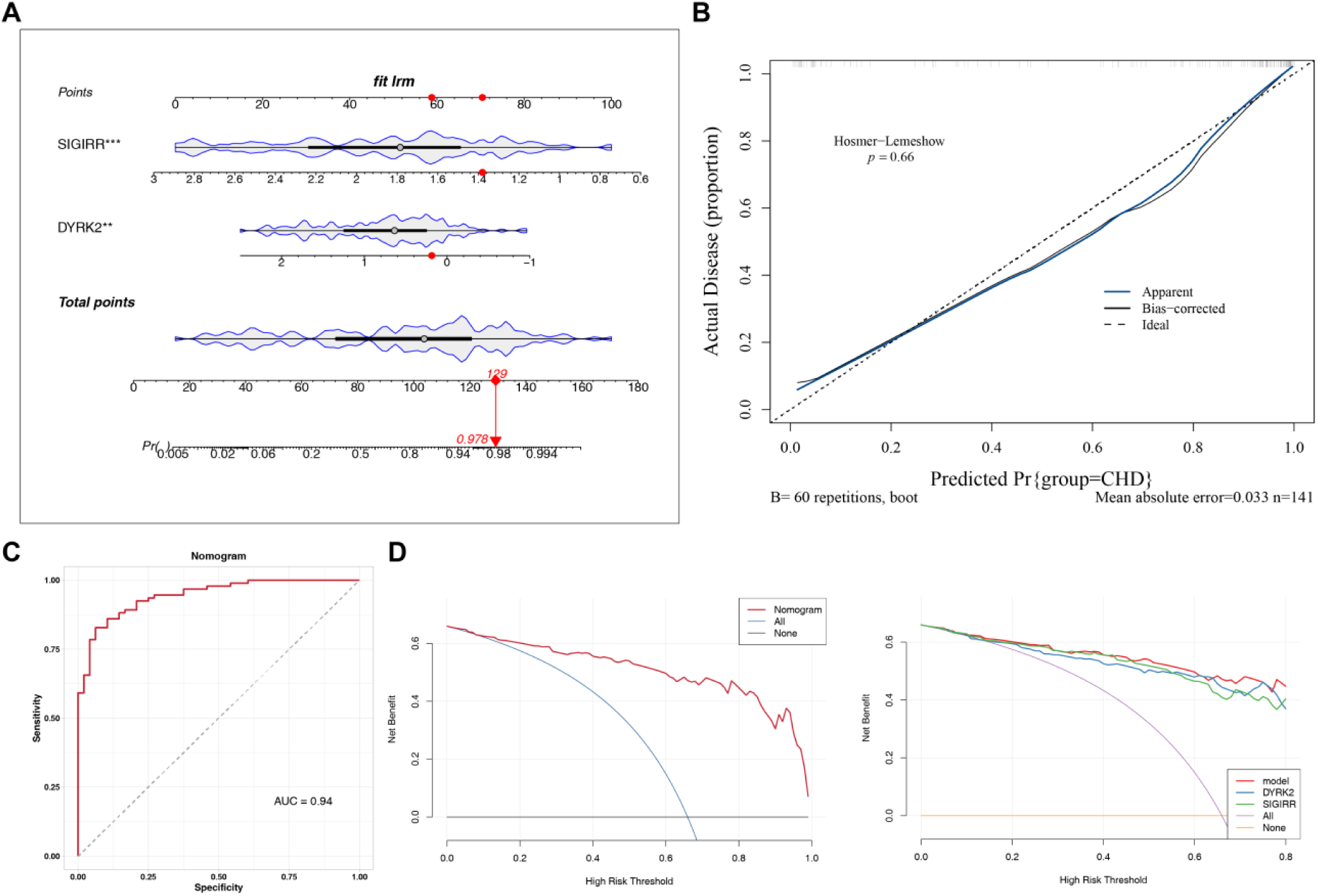
Construction and evaluation of a diagnostic nomogram. (A) Nomogram integrating DYRK2 and SIGIRR expression to predict CHD risk, where higher total points indicate increased probability of CHD. (B) Calibration curve demonstrating agreement between predicted and observed outcomes (Hosmer-Lemeshow test, P = 0.66). (C) ROC curve for the nomogram model, showing high diagnostic accuracy (AUC = 0.94). (D) Decision curve analysis (DCA) comparing net benefits of the nomogram versus single-gene models across threshold probabilities.

### 3.4 Functional annotation of key genes

GSEA revealed distinct yet partially overlapping pathway enrichment profiles for DYRK2 and SIGIRR. DYRK2 was significantly associated with pathways including the Cd80/Cd86/Ctla4/Pp2a signaling pathway, activation of PRC2.2 by H2AK119 ubiquitination, and the Pp2a/Akt signaling pathway (|NES| > 1, adjusted P < 0.05, FDR < 0.25) (**Figure 4A, Tables S4)**. SIGIRR enrichment was prominent in pathways related to impaired PINK1-mediated electron transfer in complex I and aberrant SNCA-associated complex I dysfunction (|NES| > 1, adjusted P < 0.05, FDR < 0.25) (**Figure 4B, Tables S5)**. Notably, 14 pathways were co-enriched by both genes, with prominent overlaps in mitochondrial complex I-related pathways involving PINK1 inactivation, aberrant TDP-43, and reference electron transfer (**Figure 4C-D**). GeneMANIA analysis revealed that DYRK2 and SIGIRR interact with 20 functionally related genes through physical interactions, co-expression, predicted interactions, and co-localization, implicating processes such as toll-like receptor signaling, pattern recognition receptor signaling, and interleukin-8 production and regulation (**Figure 4E**). These findings collectively suggest that DYRK2 and SIGIRR converge on mitochondrial dysfunction and innate immune regulation, potentially driving CHD pathogenesis through coordinated modulation of inflammatory signaling and cellular energy homeostasis.

**Figure 4.**
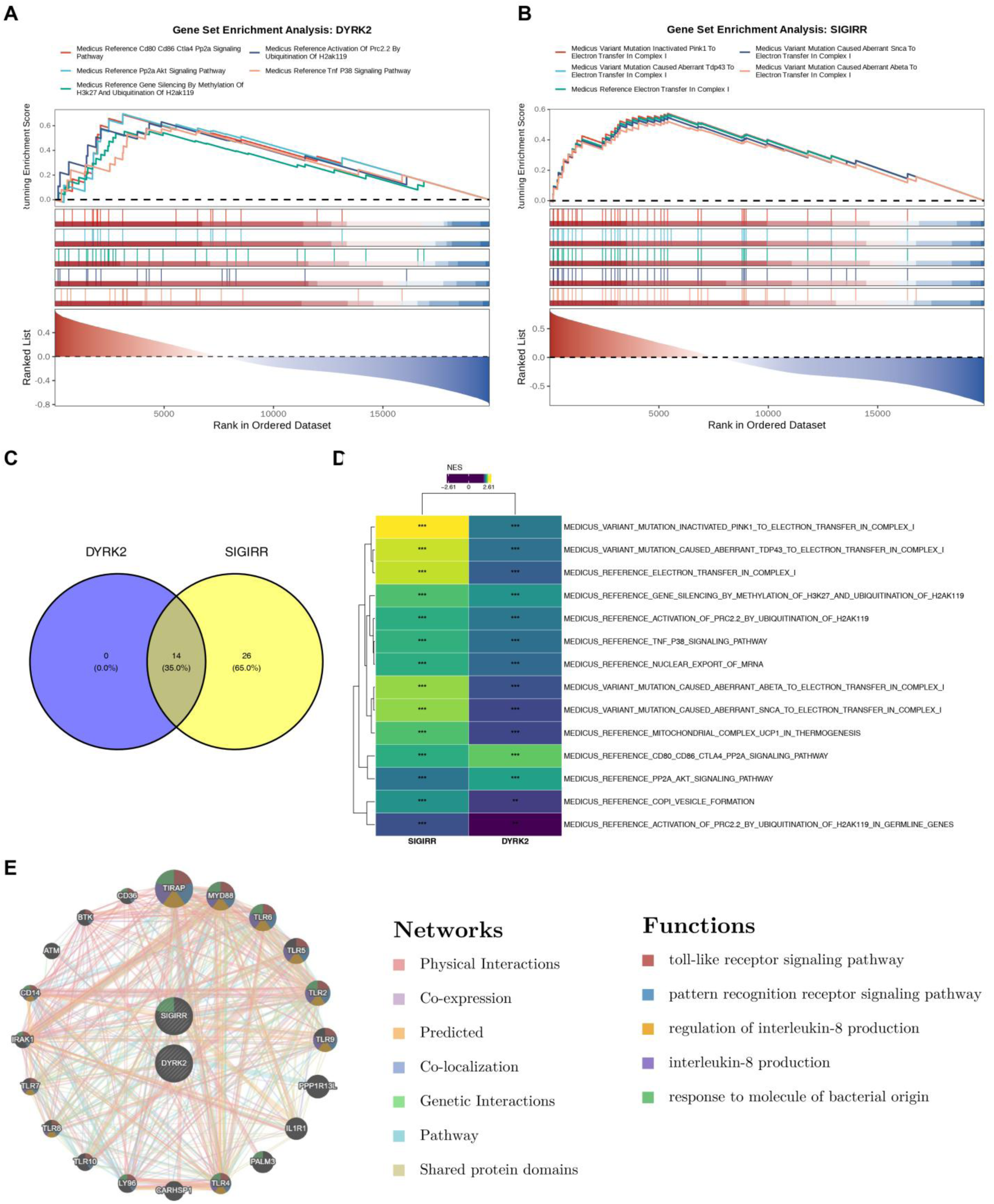
Functional annotation of key genes. (A) Gene set enrichment analysis (GSEA) results for DYRK2, showing top-enriched pathway. (B) GSEA results for SIGIRR. (C) Venn diagram illustrating 14 shared enriched pathways. (D) Heatmap of pathways co-enriched by both DYRK2 and SIGIRR. (E) GeneMANIA network depicting interactions between SIGIRR, DYRK2, and 20 related genes, involving biological processes such as toll-like receptor signaling and interleukin-8 production.

### 3.5 Immune microenvironment characterization and regulatory mechanisms

Comprehensive profiling of immune cell composition revealed extensive remodeling of the immunological landscape in CHD. Among adaptive immune populations, activated CD4⁺ T cells, activated CD8⁺ T cells, CD56dim NK cells, central memory CD4⁺ T cells, effector memory CD8⁺ T cells, immature dendritic cells, MDSCs, memory B cells, NKT cells, plasmacytoid dendritic cells, and Th2 cells were all substantially depleted in CHD tissue (P < 0.001). Conversely, innate immune effectors — including monocytes, neutrophils, macrophages, Treg cells, Th17 cells, and immature B cells — were significantly expanded (P < 0.05) (**Figure 5A–B**). Inter-cellular correlation analysis further delineated coherent co-infiltration patterns. Effector memory CD8⁺ T cells exhibited robust positive associations with activated CD4⁺ T cells (r = 0.55, P < 0.001), activated CD8⁺ T cells (r = 0.78, P < 0.001), and central memory CD4⁺ T cells (r = 0.56, P < 0.001), collectively suggesting coordinated adaptive immune contraction. Within the innate compartment, macrophages co-varied positively with monocytes (r = 0.65, P < 0.001) and neutrophils (r = 0.69, P < 0.001). Reciprocal antagonism between innate and adaptive populations was evidenced by negative correlations of monocytes with activated CD8⁺ T cells (r = −0.40, P < 0.001) and effector memory CD8⁺ T cells (r = −0.45, P < 0.001), and of Treg cells with activated CD4⁺ T cells (r = −0.46, P < 0.001) (**Figure 5C**). Gene–immune cell association analyses demonstrated that both DYRK2 and SIGIRR expression correlated positively with CD56dim NK cells and central memory CD4⁺ T cells (r > 0.50, P < 0.001). SIGIRR further exhibited significant positive associations with activated CD8⁺ T cells and effector memory CD8⁺ T cells (r > 0.60, P < 0.001), alongside a significant negative correlation with Treg cells (r < −0.40, P < 0.001) (**Figure 5D**). Chromosomal localization mapped DYRK2 to chromosome 12 and SIGIRR to chromosome 11 (**Figure 5E**). Upstream regulatory network reconstruction predicted 71 miRNAs with potential targeting capacity for DYRK2, among which 24 were linked to 91 lncRNAs, yielding regulatory axes including NORAD–hsa-miR-92b-3p–DYRK2 and OIP5-AS1–hsa-miR-92a-3p– DYRK2; no corresponding miRNA–lncRNA interactions were predicted for SIGIRR (**Figure 5F**). Pharmacological network analysis identified 59 candidate compounds with predicted affinity for DYRK2 — notably GW5074, staurosporine, and a PKR inhibitor — while no druggable interactions were identified for SIGIRR (**Figure 5G**).Taken together, these findings delineate a profound immunological imbalance in CHD, characterized by selective attrition of adaptive immune effectors concurrent with innate immune amplification. The identified lncRNA–miRNA–DYRK2 regulatory axes offer mechanistic insight into transcriptional control of key ARGs, while the candidate therapeutic compounds provide a rational basis for future DYRK2-targeted intervention strategies.

**Figure 5.**
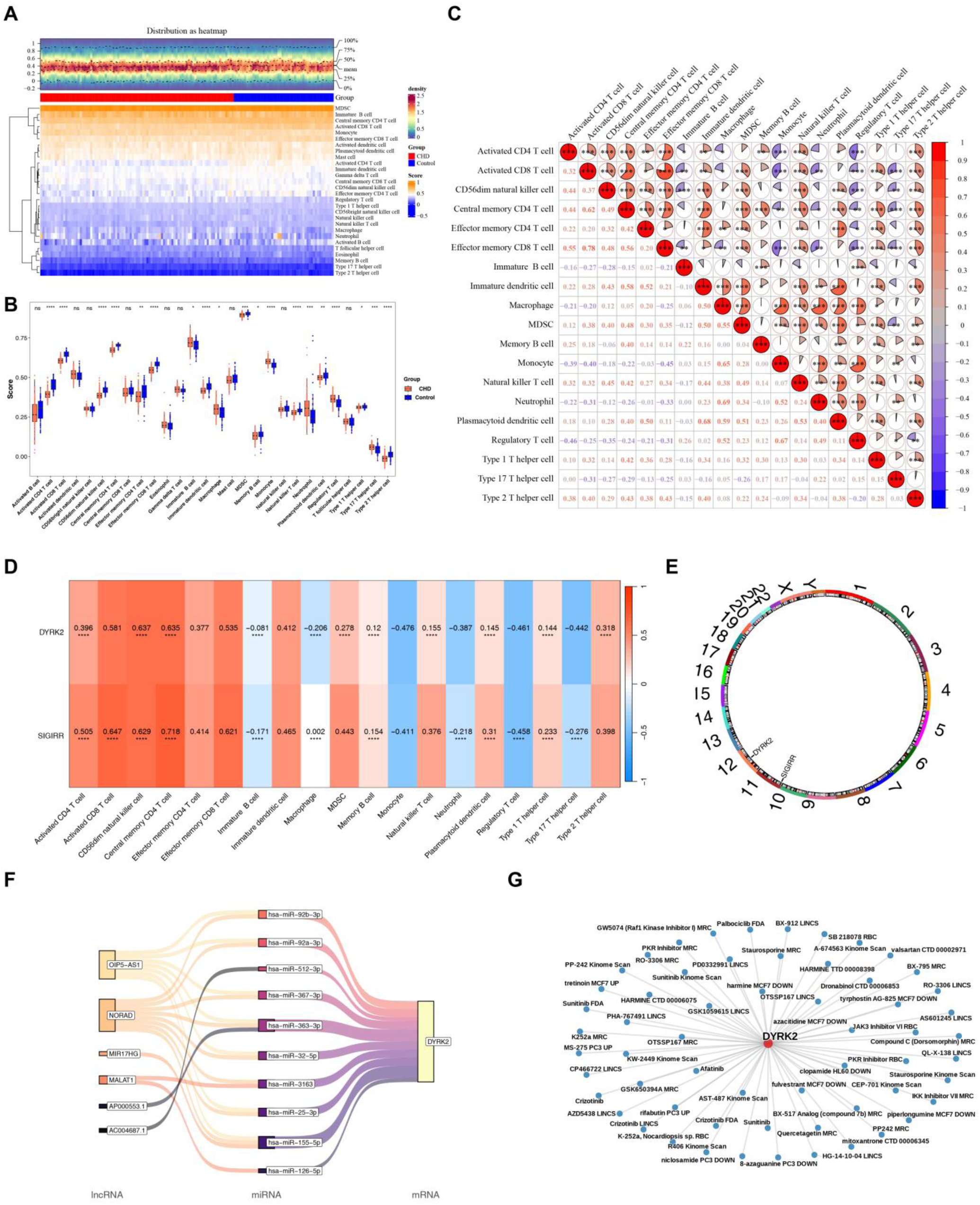
Immune microenvironment characterization and regulatory mechanisms. (A) Heatmap comparing immune cell infiltration abundances between CHD samples and controls. (B) Box plots summarizing differentially infiltrated immune cells (P < 0.05). (C) Correlation heatmap of differentially infiltrated immune cells, showing strong positive associations and negative correlations. (D) Correlation heatmap demonstrating correlations between key genes (DYRK2, SIGIRR) and differentially infiltrated immune cells. (E) Chromosomal location plot indicating DYRK2 on chromosome 12 and SIGIRR on chromosome 11. (F) Regulatory network. (G) Network plot of predicted therapeutic drugs targeting DYRK2, such as GW5074 and staurosporine. Statistical significance symbols: *P < 0.05; **P < 0.01; ***P < 0.001; ****P < 0.0001; ns, not significant.

### 3.6 Single-cell transcriptomic profiling identified specific dysregulation of key genes

After quality control, 16, 330 cells and 25, 742 genes were retained for scRNA-seq analysis **(Figure S1A)**. The top 2, 000 HVGs were selected for downstream processing **(Figure S1B)**, and the first 20 PCs were determined to capture the primary sources of biological variation **(Figure S1C)**. Unsupervised clustering resolved nine cell clusters **(Figure S1D)**, subsequently annotated into five cell types—cardiomyocytes (CMs), endothelial cells (ECs), fibroblasts (FBs), macrophages (MPs), and SMCs—based on canonical marker genes (**Figure 6A**). CMs predominated in both groups, while ECs were enriched in CHD samples and FBs in controls (**Figure 6B**). Both DYRK2 and SIGIRR were significantly downregulated in SMCs from CHD samples compared to controls (P < 0.05) (**Figure 6C**). Given the central role of SMCs in atherosclerotic plaque formation and CHD pathogenesis, SMCs were designated as the key cell type for further investigation. Subclustering of SMCs identified two subtypes **(Figure S2B)**, with distinct marker genes **(Figure S2C)**. Pseudotime trajectory analysis mapped subtype 1 to early differentiation stages and subtype 0 to late stages (**Figure 6D**). During SMC differentiation, DYRK2 exhibited an initial high expression followed by a decline in early stages, transient reactivation during mid-differentiation, and subsequent downregulation in late stages. In contrast, SIGIRR decreased progressively (**Figure 6E**). These results pinpointed SMCs as a critical cellular compartment in CHD pathogenesis, with DYRK2 and SIGIRR exhibiting stage-specific dysregulation during SMC differentiation. The aberrant expression dynamics suggested potential roles in SMC phenotypic modulation and vascular pathology.

**Figure 6.**
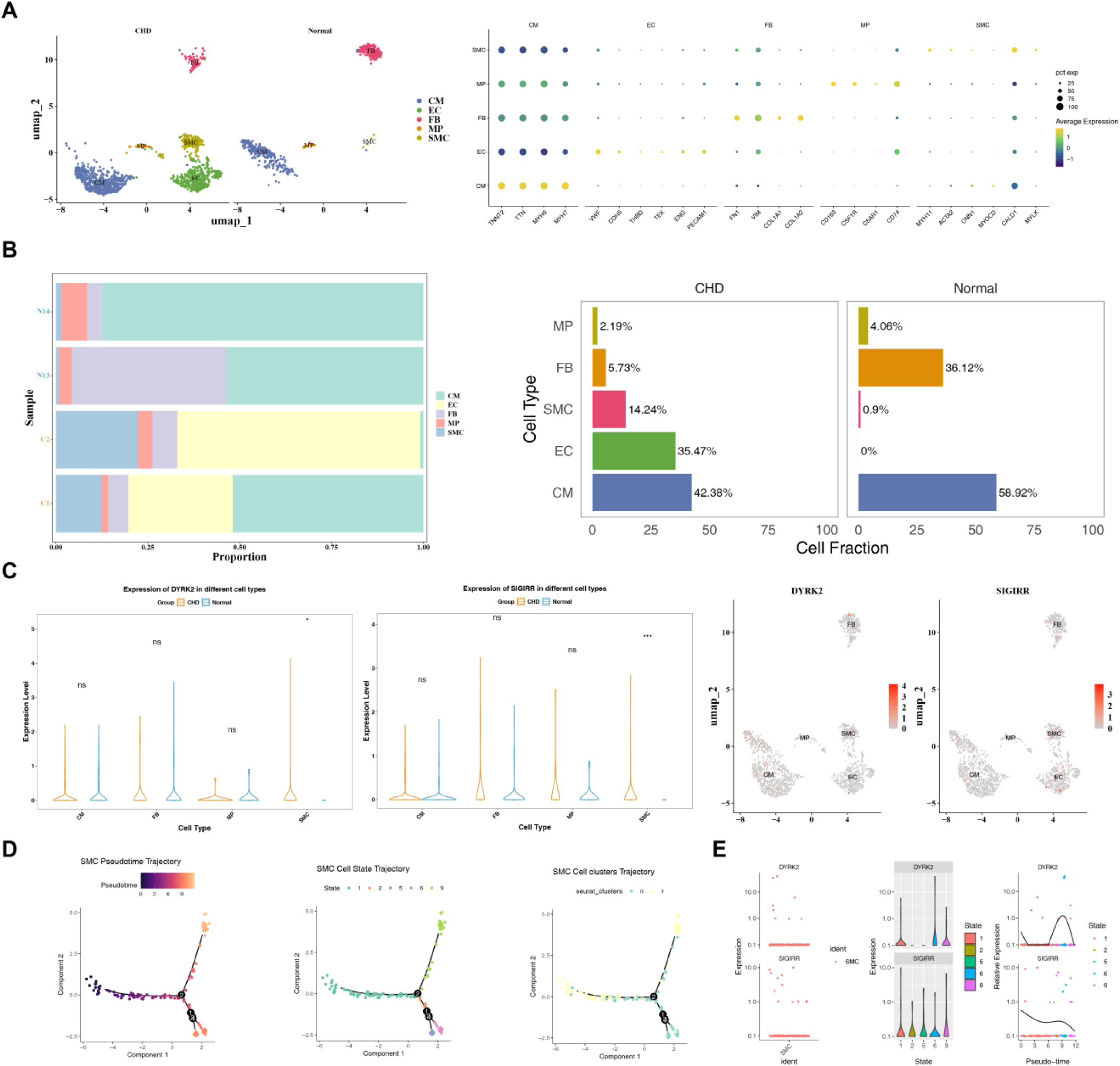
Single-cell transcriptomic profiling identified specific dysregulation of key genes. (A) Uniform manifold approximation and projection (UMAP) plot annotating five cell types: cardiomyocytes (CMs), endothelial cells (ECs), fibroblasts (FBs), macrophages (MPs), and smooth muscle cells (SMCs). (B) Bar plots showing relative abundances of cell types in CHD samples versus controls, with ECs enriched in CHD and FBs in controls. (C) Violin plots comparing DYRK2 and SIGIRR expression in annotated cell types between CHD samples and controls, revealing significant upregulation (P < 0.05). (D) Pseudotime trajectory analysis of SMCs. (E) Pseudotime trajectory analysis of SMCs, illustrating expression dynamics of DYRK2 and SIGIRR during differentiation. Statistical significance symbols: *P < 0.05; **P < 0.01; ***P < 0.001; ****P < 0.0001; ns, not significant.

### 3.7 Metabolic pathway alterations and rewired intercellular communication

Metabolic pathway analysis revealed cell-type-specific alterations in CHD. Ribosome pathway activity was markedly suppressed in SMCs and ECs, while CMs and MPs exhibited high enrichment. Allograft rejection pathways showed minimal activity in CMs, whereas MPs demonstrated significant enrichment in antigen processing and presentation, indicating cell-type-dependent metabolic reprogramming in CHD pathogenesis (**Figure 7A**). Cell-cell communication networks underwent substantial reorganization in CHD samples compared to controls. In controls, SMCs maintained frequent interactions with FBs, and FBs communicated extensively with CMs (**Figure 7B**). Conversely, CHD samples exhibited diminished SMC-FB interactions alongside enhanced SMC-CM crosstalk (**Figure 7C**), suggesting CHD-associated rewiring of stromal-vascular communication. Ligand-receptor pair analysis identified IGFBP3-TMEM219 and GRN-SORT1 as dominant signaling axes in controls. SMCs functioned bidirectionally in this network, notably sending PTN-SDC2 signals to FBs (**Figure 7D**). In CHD samples, SMCs primarily received PTN-NCL signals from FBs, indicating a shift toward unidirectional signaling (**Figure 7E**). Pathway-specific interrogation revealed spatial reorganization of signaling hubs. In controls, SMCs showed a strong VEGF signal input while CMs demonstrated a prominent VEGF output (**Figure 7F, first plot)**. CHD samples exhibited enhanced VEGF input in ECs with maintained CM output (**Figure 7F, second plot)**. For the PTN pathway, controls featured robust PTN input/output in FBs but minimal activity in SMCs (**Figure 7F, third plot)**. CHD samples displayed attenuated FB PTN signaling alongside intensified PTN output from SMCs (**Figure 7F, fourth plot)**. These results delineate a CHD-specific reconfiguration of intercellular communication networks, with aberrant VEGF and PTN signaling potentially driving pathological vascular remodeling through altered SMC-EC-FB cross-talk.

**Figure 7.**
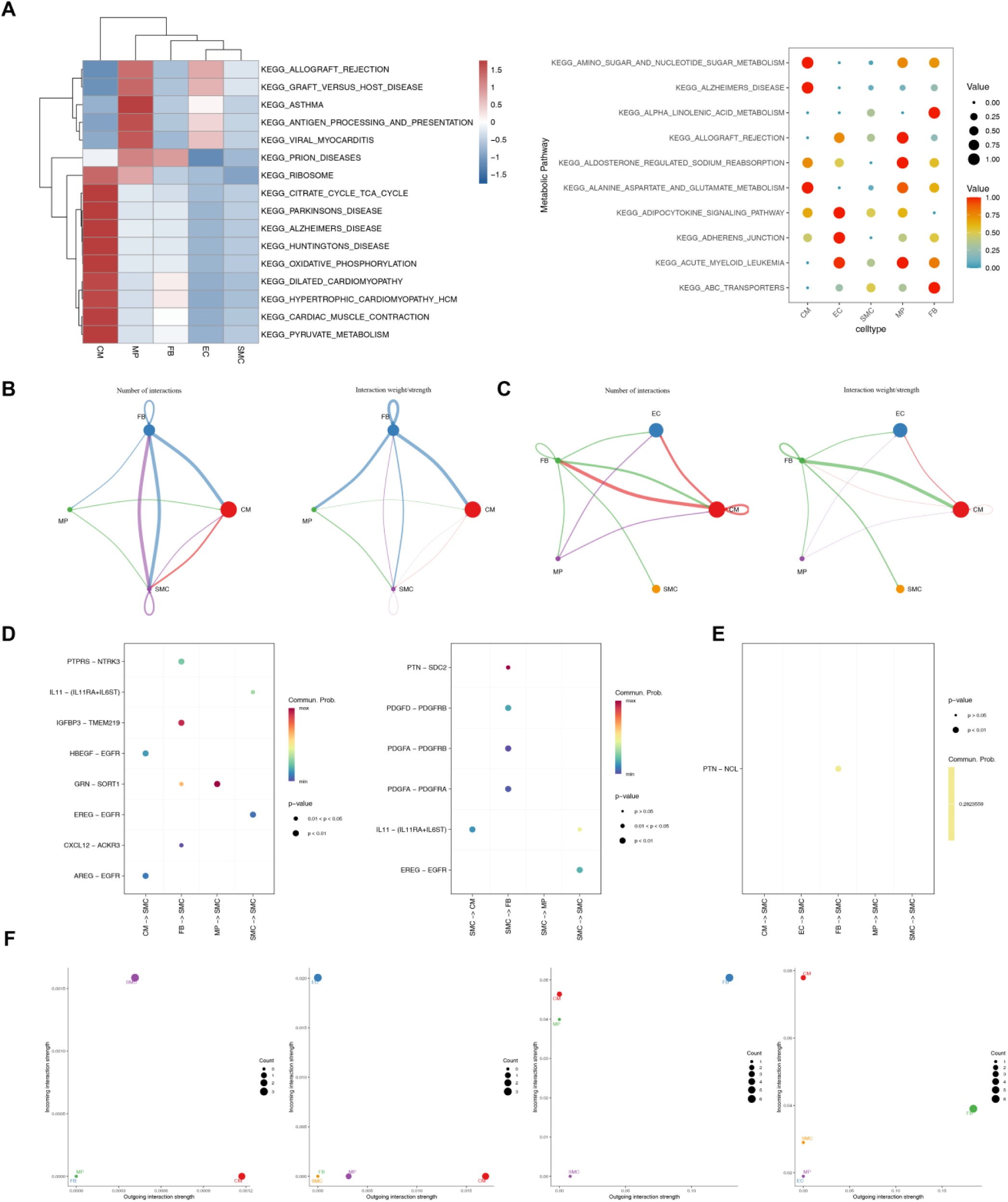
Metabolic pathway alterations and rewired intercellular communication. (A) The pathway enrichment scores across cell types, showing suppression of the ribosome pathway in SMCs and ECs, and enrichment of antigen processing in MPs. (B-C) Cell-cell communication networks in controls (B) and CHD samples (C), highlighting diminished SMC-FB interactions and enhanced SMC-CM crosstalk in CHD. (D-E) Dominant ligand-receptor pairs in controls and CHD samples. (F) Spatial visualization of vascular endothelial growth factor (VEGF) signaling input/output in controls (first plot) and CHD samples (second plot), followed by pleiotrophin (PTN) signaling in controls (third plot) and CHD samples (fourth plot); CHD shows enhanced VEGF input in ECs and intensified PTN output from SMCs.

### 3.8 Expression levels of key genes

To validate the bioinformatics predictions, the mRNA expression levels of DYRK2 and SIGIRR were quantitatively analyzed in CHD samples (n = 5) and control samples (n = 5) using RT-qPCR. Both genes were significantly downregulated in CHD samples compared to controls (P < 0.05) (**Figure 8**), consistent with the bioinformatics predictions, suggesting their potential involvement in CHD pathogenesis.

**Figure 8.**
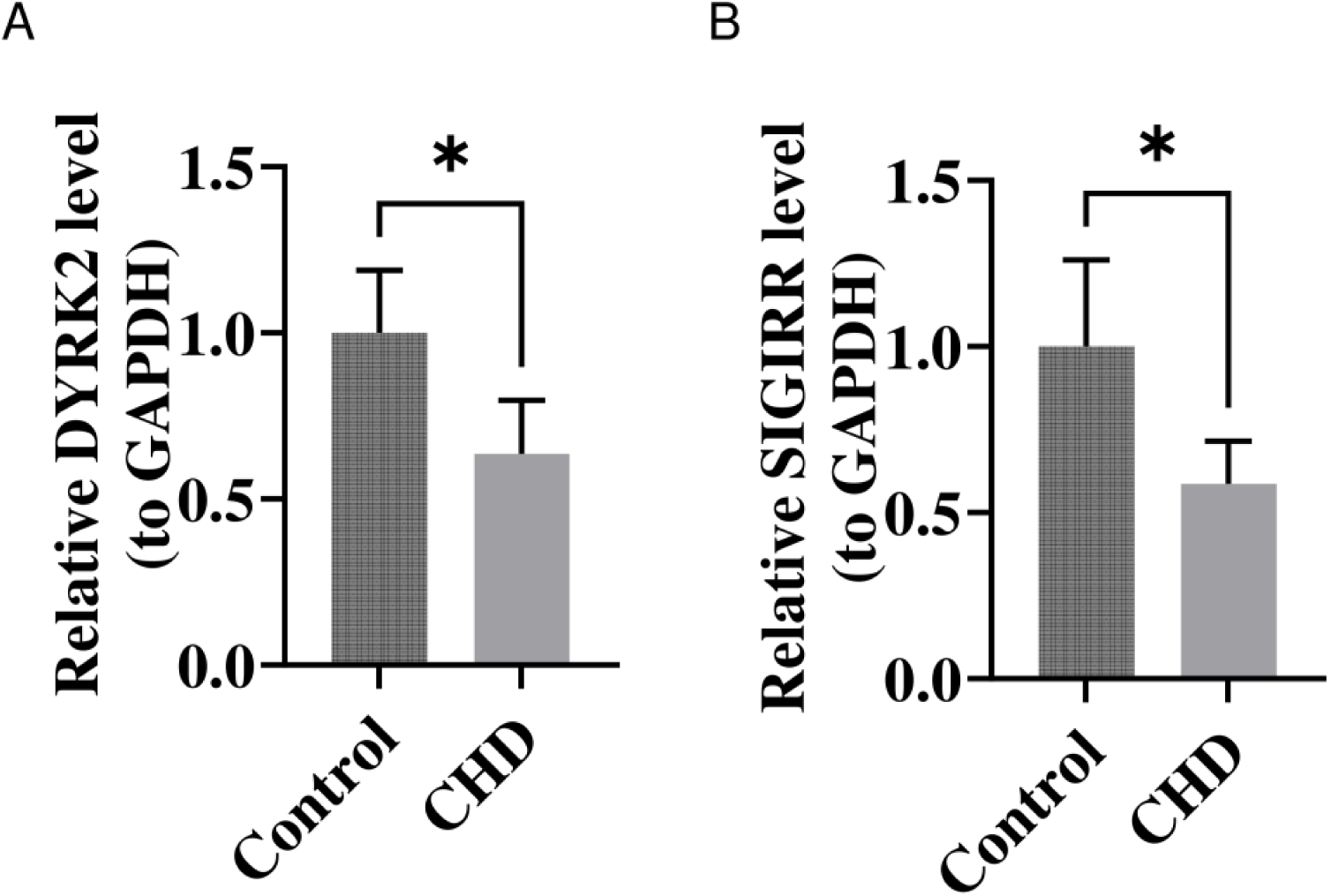
RT-qPCR validation of DYRK2 and SIGIRR expression in CHD samples. RT-qPCR analysis of mRNA expression levels in CHD samples (n = 5) and control samples (n = 5). (A) DYRK2 expression. (B) SIGIRR expression. Relative expression was calculated using the 2^−ΔΔCt^ method with GAPDH as the internal control. Data are presented as mean ± SD. Statistical significance was determined by unpaired Student’s t-test. *P < 0.05; **P < 0.01; ns, not significant.

## 4 Discussion

CHD stands as the primary cause of death globally, and its pathogenesis encompasses intricate interactions among metabolic disorders, immune dysregulation, and vascular remodeling^3, 34, 35^. Nevertheless, the ARGs governing these pathological processes and their crosstalk with immune cells remain insufficiently characterized. The present study fills this gap by providing, for the first time, a systematic characterization of ARG involvement in CHD pathogenesis. Through delineating the mechanistic contributions of two hub genes - DYRK2 and SIGIRR - this work introduces a novel molecular framework for understanding how the adenosine signaling pathway underlies CHD development.

Building upon the identification of these key genes, our analyses revealed that DYRK2 and SIGIRR are significantly downregulated in PBMCs of patients with CHD, and the two genes synergistically promote disease pathology. DYRK2 belongs to the CMGC kinase family and regulates the cell cycle and energy metabolism via phosphorylation^36, 37^. GSEA and Kyoto KEGG analysis indicated that DYRK2 is significantly enriched in mitochondrial ETC complex I and the Hedgehog signaling pathway. Its deficiency may result in mitochondrial damage, ROS accumulation, impairment of cellular stress maintenance^38^ and protein homeostasis^37^, thereby influencing vascular development and repair^39^ and exacerbating CHD progression by mediating metabolic inflammatory crosstalk, thus participating in disease pathogenesis through multiple molecular pathways^39, 40^. Complementing DYRK2’s metabolic role, SIGIRR, as a crucial negative regulator of TLR/IL-1R^41^, leads to excessive activation of the immune response and exacerbation of vascular inflammation by affecting the TLR pathway^41^, CD18 protein stability^42^, IL - 37 function^43^, and the adenosine signaling pathway. The deficiency of DYRK2 results in impaired electron transport chain (ETC) function, particularly affecting complex I, which in turn increases mitochondrial reactive oxygen species (ROS) release and promotes the maturation and secretion of pro-inflammatory cytokines like IL-1β. Concurrently, the downregulation of SIGIRR lifts the suppression of receptor signaling pathways, such as TLR4, thereby amplifying the inflammatory response. This self-perpetuating “energy crisis-inflammatory storm” positive feedback loop plays a pivotal role in exacerbating the instability of atherosclerotic plaques, promoting thrombosis, and ultimately leading to cardiovascular events.

Immune cell infiltration profiling revealed pronounced remodeling of the immunological landscape in CHD, consistent with previous reports of adaptive immune dysregulation in atherosclerosis ^44^. Adaptive immune populations — including activated CD4⁺ T cells, activated CD8⁺ T cells, and CD56dim NK cells — were substantially diminished, while innate immune effectors such as monocytes, macrophages, and neutrophils were markedly enriched. This immunological transition is governed by the coordinated actions of DYRK2 and SIGIRR. DYRK2 expression correlated positively with CD56dim NK cells and central memory CD4⁺ T cells; its downregulation compromises NK cytotoxicity and T cell memory, thereby weakening immune surveillance of vascular injury. Furthermore, elevated ROS resulting from DYRK2 loss promotes monocyte and macrophage recruitment and pro-inflammatory polarization, amplifying innate immune responses^45^. SIGIRR was additionally associated with activated and effector memory CD8⁺ T cells, and inversely correlated with Treg abundance. SIGIRR deficiency suppresses effector T cell activation — particularly of CD8⁺ T cells — intensifying adaptive immune dysfunction ^44^ and indirectly favoring Treg expansion or functional deviation ^46^. This compounded immune imbalance impairs vascular repair mechanisms, sustains local inflammation, and accelerates atherosclerotic plaque progression and rupture.

To further elucidate the cell-type-specific roles of DYRK2 and SIGIRR, single-cell clustering analysis successfully identified five primary cell types: CMs, ECs, FBs, MPs, and vascular SMCs. Notably, in contrast to their downregulation in circulating PBMCs, expression profiling revealed a significant upregulation of DYRK2 and SIGIRR in SMCs derived from CHD samples, suggesting distinct regulatory mechanisms and functional roles in different cellular compartments. The phenotypic plasticity of SMCs represents a critical pathological feature of CHD^47^. To delve deeper into the role of SMCs in CHD, we performed a subpopulation analysis of SMCs and observed that DYRK2 expression peaked transiently before declining during the later stages of differentiation, suggesting its crucial regulatory role at specific stages of SMC differentiation, such as proliferation or early phenotypic transition. Conversely, SIGIRR expression exhibited a gradual decline throughout differentiation. The overall upregulation of these two genes contradicts the normal differentiation timeline, strongly indicating that, under pathological conditions of CHD, abnormalities occur in the differentiation process or gene expression regulation, resulting in the sustained or abnormally high expression of these genes at “inappropriate” stages, thereby driving or perpetuating the pathological phenotype of SMCs. Beyond these cell-intrinsic alterations, this study unveils that disruption of ARGs results in significant reconfiguration of the intercellular communication network in CHD, with aberrant signals from the core pathways, VEGF and PTN, driving vascular remodeling. The activity of the ribosome pathway is markedly suppressed in SMCs and ECs within CHD, whereas it is highly enriched in CMs and MPs. The interaction between SMCs and FBs in CHD samples is attenuated, while communication between SMCs and CMs is intensified, suggesting that the stromal-vascular cell communication network undergoes reprogramming in CHD. In CHD samples, SMCs transition from bidirectional signaling to primarily receiving PTN-NCL signals from FBs, indicating a shift in the directionality of signal transmission. The VEGF signal input to ECs in CHD samples is augmented, while the output from CMs remains stable. These alterations may impact angiogenesis and endothelial function. PTN signals emanating from FBs are diminished in CHD samples, whereas PTN output from SMCs is notably increased. SMCs emerge as the primary source of PTN signals, potentially abnormally promoting their own proliferation, migration, or inflammatory response through autocrine or paracrine mechanisms, thereby driving pathological vascular remodeling.

Given the central role of DYRK2 in CHD pathogenesis revealed by our multi-level analyses, we explored its therapeutic targeting potential. This study identified 59 potential therapeutic drugs that might target DYRK2, including GW5074, staurosporine, and PKR inhibitors^48, 49^. Although not originally designed for cardiovascular applications, these compounds provide an important starting point for developing DYRK2-targeted CHD therapies.

In conclusion, this study has, for the first time, identified DYRK2 and SIGIRR as pivotal genes associated with CHD. Functional analyses indicate that these two genes are jointly implicated in mitochondrial complex I dysfunction and the dysregulation of innate immune inflammatory responses. Given the central role of the adenosine signaling pathway in cardiovascular and immune regulation, it is postulated that abnormalities in this pathway within CHD may, through the downregulation of DYRK2 and SIGIRR expression, synergistically induce mitochondrial energy metabolism disorders and uncontrolled innate immune inflammatory responses, thereby collectively contributing to the onset and progression of CHD. Moreover, at the single-cell level, this study unveils the dynamic regulatory anomalies of DYRK2 and SIGIRR, as well as the reconfiguration of intercellular communication networks during SMC phenotypic transition, offering novel insights into the vascular remodeling mechanisms underlying CHD.

Several limitations merit consideration. This investigation is primarily based on retrospective analysis of publicly available GEO datasets, which may introduce batch effects and inter-platform variability. The mechanistic roles of DYRK2 and SIGIRR are largely inferred from bioinformatics analyses and await confirmation through experimental validation in vitro and in vivo. Moreover, the study focuses on transcriptional profiles of ARGs while providing limited insight into dynamic changes in adenosine metabolite concentrations, receptor occupancy states, and downstream effector signaling. Future experimental studies are therefore necessary to substantiate the functional mechanisms of DYRK2, SIGIRR, and related ARGs in CHD pathobiology.

## 5 Conclusion

This study systematically identified DYRK2 and SIGIRR as key adenosine-related genes involved in the pathogenesis of coronary heart disease. By integrating bulk transcriptomic analysis, machine learning, immune infiltration profiling, and single-cell RNA sequencing, we demonstrated that these two genes were consistently dysregulated in CHD and showed promising diagnostic value when incorporated into a nomogram model. Functional analyses suggested that DYRK2 and SIGIRR may contribute to CHD progression through mitochondrial dysfunction, innate immune activation, and disruption of immune-metabolic homeostasis. At the single-cell level, smooth muscle cells emerged as a critical cellular compartment, with abnormal expression dynamics of these genes during phenotypic transition and marked remodeling of VEGF- and PTN-related intercellular communication. Collectively, these findings provide new insight into adenosine-related molecular mechanisms underlying CHD and highlight DYRK2 and SIGIRR as potential biomarkers and therapeutic targets. Further experimental validation is warranted to confirm their biological functions and clinical relevance.

## Non-standard Abbreviations and Acronyms

ARGs: adenosine-related genes
AUC: area under the curve
BP: biological process
CC: cellular component
CHD: coronary heart disease
CMs: cardiomyocytes
DCA: decision curve analysis
DEGs: differentially expressed genes
ECs: endothelial cells
ETC: electron transport chain
FBs: fibroblasts
GEO: Gene Expression Omnibus
GO: Gene Ontology
GSEA: gene set enrichment analysis
HVGs: highly variable genes
KEGG: Kyoto Encyclopedia of Genes and Genomes
lncRNA: long non-coding RNA
miRNA: microRNA
MPs: macrophages
NK: natural killer
PBMCs: peripheral blood mononuclear cells
PPI: protein-protein interaction
PTN: pleiotrophin
ROC: receiver operating characteristic
ROS: reactive oxygen species
scRNA-seq: single-cell RNA sequencing
SMCs: smooth muscle cells
ssGSEA: single-sample gene set enrichment analysis
SVM-RFE: support vector machine-recursive feature elimination
TLR: toll-like receptor
UMAP: uniform manifold approximation and projection
VEGF: vascular endothelial growth factor
WGCNA: weighted gene co-expression network analysis

## Acknowledgments

We would like to express our sincere gratitude to all individuals and organizations who supported and assisted us throughout this research. In conclusion, we extend our thanks to everyone who has supported and assisted us along the way. Without your support, this research would not have been possible.

## Sources of Funding

This research received no specific grant from funding agencies in the public, commercial, or not-for-profit sectors.

## Disclosures

The authors declare that the research was conducted in the absence of any commercial or financial relationships that could be construed as a potential conflict of interest.

## Author Contributions

KC: Conceptualization, Data curation, Validation, Visualization, Writing–original draft, Writing– review & editing. RW, YQ: Data curation, Validation, Visualization, Writing–review & editing. XZ, ZX, XM, HM: Validation, Writing–review & editing. NW, ZL, LC, XW: Visualization, Writing– review & editing. KS: Conceptualization, Supervision, Writing–review & editing. LZ: Conceptualization, Project administration, Supervision, Writing–review & editing. This work currently described has not been published, is not being considered for publication elsewhere, and its publication was approved by all authors.

## Ethical approval and consent to participate

This study was performed in line with the principles of the Declaration of Helsinki. Approval was granted by the Ethics Committee of The 941th Hospital of Joint Logistic Support Force of Chinese People’s Liberation Army, and the approval number and date of approval are [Approval No.: [2026022]] and [2026.07.14]. Informed consent was obtained from all individual participants included in the study.

## Consent for publication

Not applicable.

## Data Availability Statement

The datasets analyzed for this study can be found in the Gene Expression Omnibus (GEO) database under accession numbers GSE113079, GSE42148, and GSE121893 at [http://www.ncbi.nlm.nih.gov/geo/].

---

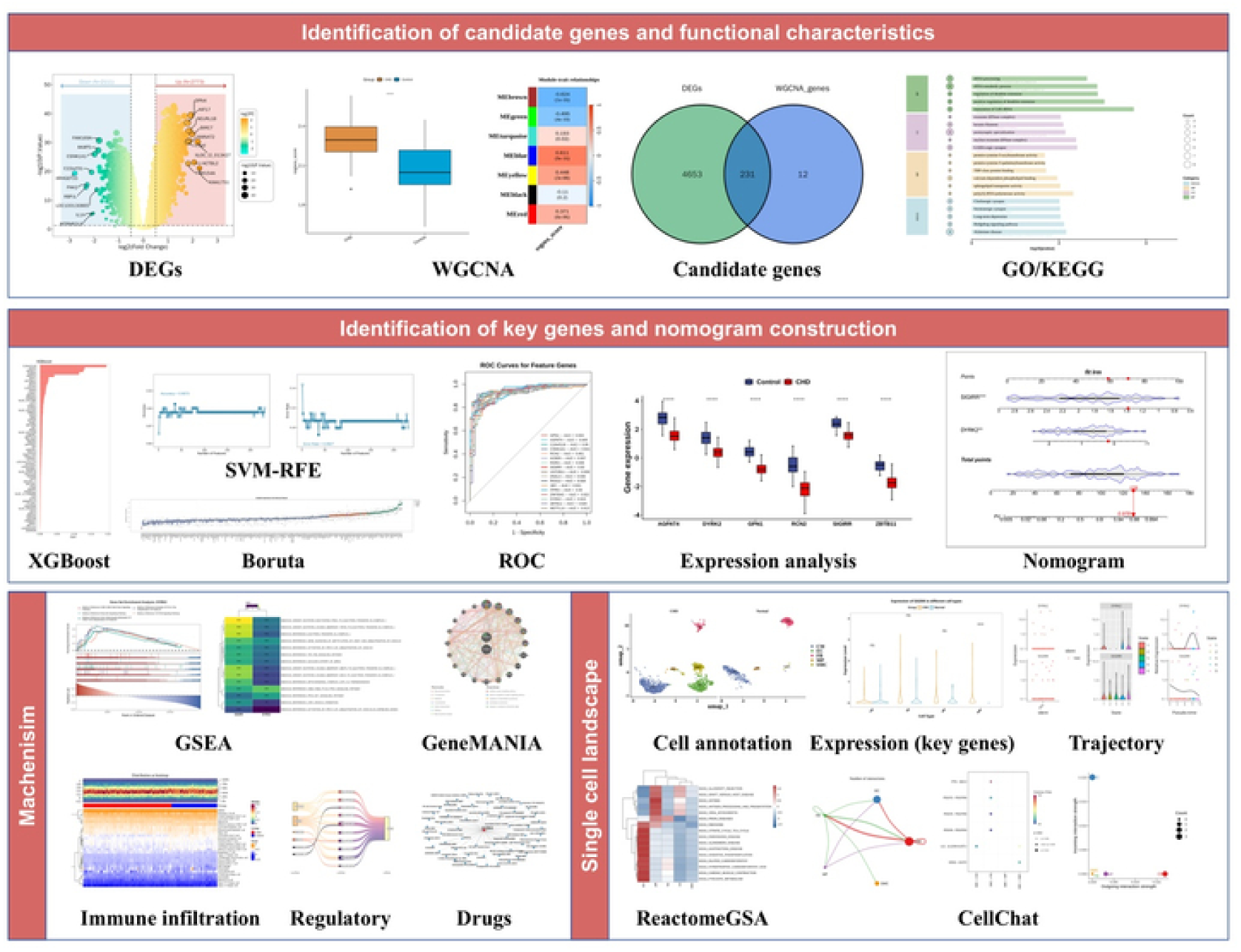

## Supplementary Material

### Supplementary Figures

**Figure S1.**
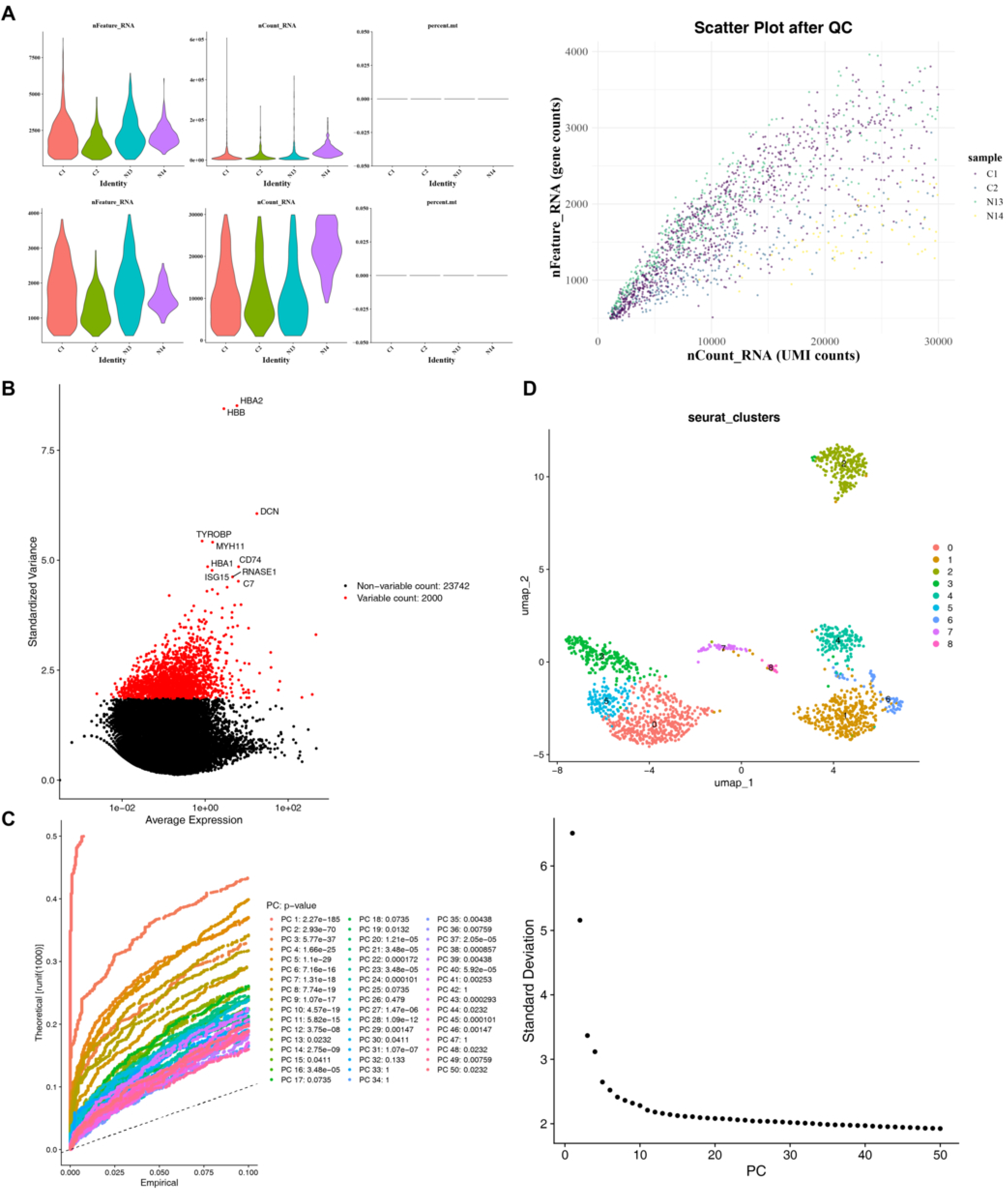
Quality control and preprocessing of scRNA-seq data. Violin plots displaying quality metrics before and after filtering. A total of 16, 330 cells and 25, 742 genes were retained. (B) Plot of highly variable genes (HVGs), with the top 2, 000 HVGs labeled for downstream analysis. (C) The contribution of principal components (PCs), where the first 20 PCs captured the majority of biological variation. (D) UMAP identifies nine distinct cell clusters before annotation.

**Figure S2.**
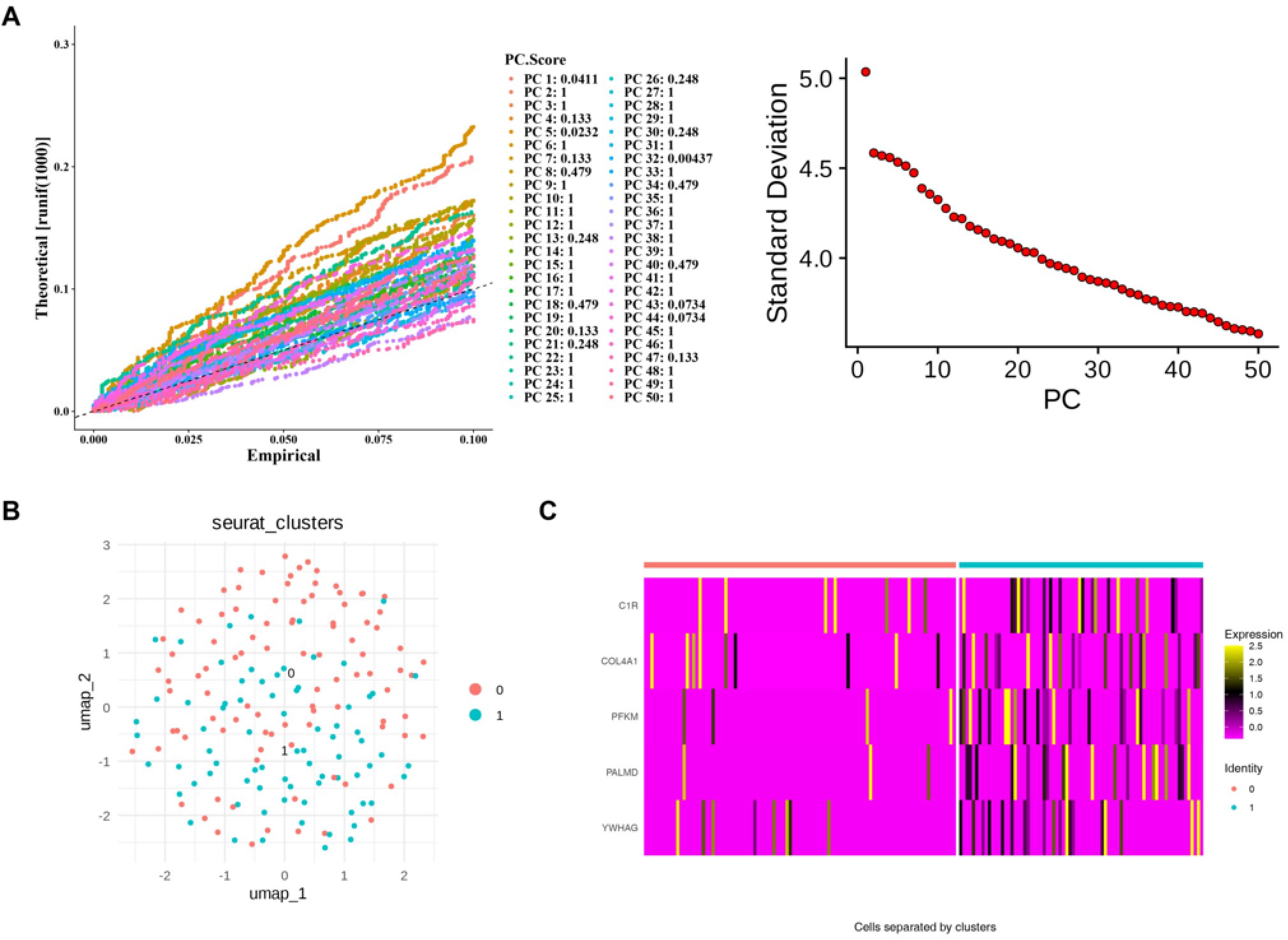
Subclustering and marker gene identification of SMCs. The contribution of PCs, where the first 50 PCs captured the majority of biological variation. (B) UMAP plot showing subclustering of SMCs into two subtypes. (C) The top five differentially expressed marker genes per SMC subtype.

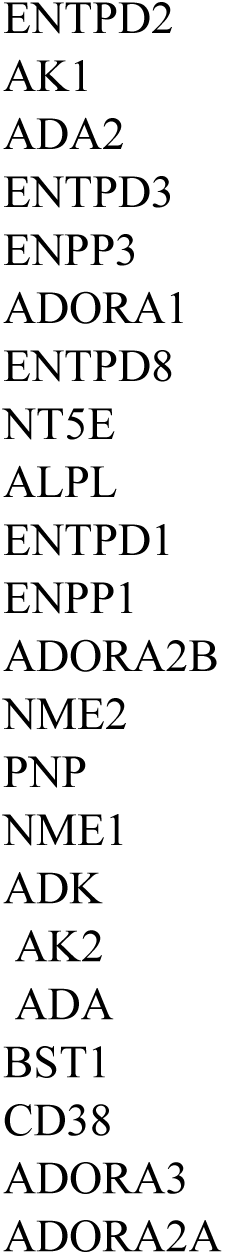

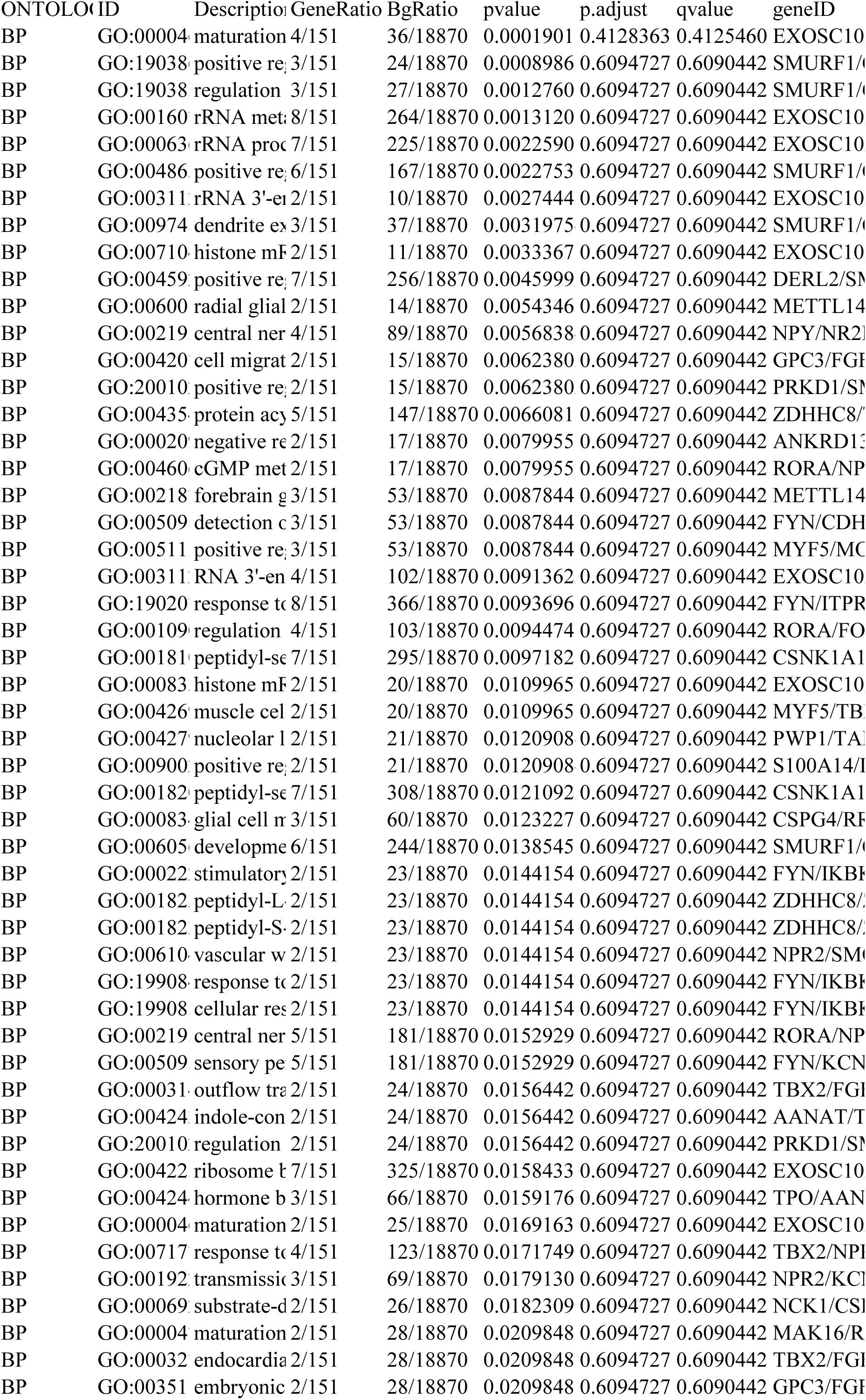

| ONTOLO | ID | Descriptio | GeneRatio | BgRatio | pvalue | p.adjust | qvalue | geneID |
|---|---|---|---|---|---|---|---|---|
| BP | GO:00004 | maturation | 4/151 | 36/18870 | 0.0001901 | 0.4128363 | 0.4125460 | EXOSC10 |
| BP | GO:19038 | positive re | 3/151 | 24/18870 | 0.0008986 | 0.6094727 | 0.6090442 | SMURF1/ |
| BP | GO:19038 | regulation | 3/151 | 27/18870 | 0.0012760 | 0.6094727 | 0.6090442 | SMURF1/ |
| BP | GO:00160 | rRNA met | 8/151 | 264/18870 | 0.0013120 | 0.6094727 | 0.6090442 | EXOSC10 |
| BP | GO:00063 | rRNA proc | 7/151 | 225/18870 | 0.0022590 | 0.6094727 | 0.6090442 | EXOSC10 |
| BP | GO:00486 | positive re | 6/151 | 167/18870 | 0.0022753 | 0.6094727 | 0.6090442 | SMURF1/ |
| BP | GO:00311 | rRNA 3'-e | 2/151 | 10/18870 | 0.0027444 | 0.6094727 | 0.6090442 | EXOSC10 |
| BP | GO:00974 | dendrite e | 3/151 | 37/18870 | 0.0031975 | 0.6094727 | 0.6090442 | SMURF1/ |
| BP | GO:00710 | histone m | 2/151 | 11/18870 | 0.0033367 | 0.6094727 | 0.6090442 | EXOSC10 |
| BP | GO:00459 | positive re | 7/151 | 256/18870 | 0.0045999 | 0.6094727 | 0.6090442 | DERL2/S |
| BP | GO:00600 | radial glial | 2/151 | 14/18870 | 0.0054346 | 0.6094727 | 0.6090442 | METTL14 |
| BP | GO:00219 | central ner | 4/151 | 89/18870 | 0.0056838 | 0.6094727 | 0.6090442 | NPY/NR2 |
| BP | GO:00420 | cell migrat | 2/151 | 15/18870 | 0.0062380 | 0.6094727 | 0.6090442 | GPC3/FG |
| BP | GO:20010 | positive re | 2/151 | 15/18870 | 0.0062380 | 0.6094727 | 0.6090442 | PRKD1/S |
| BP | GO:00435 | protein ac | 5/151 | 147/18870 | 0.0066081 | 0.6094727 | 0.6090442 | ZDHHC8/ |
| BP | GO:00020 | negative r | 2/151 | 17/18870 | 0.0079955 | 0.6094727 | 0.6090442 | ANKRD1 |
| BP | GO:00460 | cGMP met | 2/151 | 17/18870 | 0.0079955 | 0.6094727 | 0.6090442 | RORA/NP |
| BP | GO:00218 | forebrain | 3/151 | 53/18870 | 0.0087844 | 0.6094727 | 0.6090442 | METTL14 |
| BP | GO:00509 | detection | 3/151 | 53/18870 | 0.0087844 | 0.6094727 | 0.6090442 | FYN/CDH |
| BP | GO:00511 | positive re | 3/151 | 53/18870 | 0.0087844 | 0.6094727 | 0.6090442 | MYF5/M |
| BP | GO:00311 | RNA 3'-en | 4/151 | 102/18870 | 0.0091362 | 0.6094727 | 0.6090442 | EXOSC10 |
| BP | GO:19020 | response t | 8/151 | 366/18870 | 0.0093696 | 0.6094727 | 0.6090442 | FYN/ITPR |
| BP | GO:00109 | regulation | 4/151 | 103/18870 | 0.0094474 | 0.6094727 | 0.6090442 | RORA/FO |
| BP | GO:00181 | peptidyl-s | 7/151 | 295/18870 | 0.0097182 | 0.6094727 | 0.6090442 | CSNK1A1 |
| BP | GO:00083 | histone m | 2/151 | 20/18870 | 0.0109965 | 0.6094727 | 0.6090442 | EXOSC10 |
| BP | GO:00426 | muscle cel | 2/151 | 20/18870 | 0.0109965 | 0.6094727 | 0.6090442 | MYF5/TB |
| BP | GO:00427 | nucleolar | 2/151 | 21/18870 | 0.0120908 | 0.6094727 | 0.6090442 | PWP1/TA |
| BP | GO:00900 | positive re | 2/151 | 21/18870 | 0.0120908 | 0.6094727 | 0.6090442 | S100A14/ |
| BP | GO:00182 | peptidyl-s | 7/151 | 308/18870 | 0.0121092 | 0.6094727 | 0.6090442 | CSNK1A1 |
| BP | GO:00083 | glial cell | 3/151 | 60/18870 | 0.0123227 | 0.6094727 | 0.6090442 | CSPG4/R |
| BP | GO:00605 | developme | 6/151 | 244/18870 | 0.0138545 | 0.6094727 | 0.6090442 | SMURF1/ |
| BP | GO:00022 | stimulator | 2/151 | 23/18870 | 0.0144154 | 0.6094727 | 0.6090442 | FYN/IKB |
| BP | GO:00182 | peptidyl-L | 2/151 | 23/18870 | 0.0144154 | 0.6094727 | 0.6090442 | ZDHHC8/ |
| BP | GO:00182 | peptidyl-S | 2/151 | 23/18870 | 0.0144154 | 0.6094727 | 0.6090442 | ZDHHC8/ |
| BP | GO:00610 | vascular w | 2/151 | 23/18870 | 0.0144154 | 0.6094727 | 0.6090442 | NPR2/SM |
| BP | GO:19908 | response t | 2/151 | 23/18870 | 0.0144154 | 0.6094727 | 0.6090442 | FYN/IKB |
| BP | GO:19908 | cellular re | 2/151 | 23/18870 | 0.0144154 | 0.6094727 | 0.6090442 | FYN/IKB |
| BP | GO:00219 | central ner | 5/151 | 181/18870 | 0.0152929 | 0.6094727 | 0.6090442 | RORA/NP |
| BP | GO:00509 | sensory pe | 5/151 | 181/18870 | 0.0152929 | 0.6094727 | 0.6090442 | FYN/KCN |
| BP | GO:00031 | outflow tr | 2/151 | 24/18870 | 0.0156442 | 0.6094727 | 0.6090442 | TBX2/FG |
| BP | GO:00424 | indole-con | 2/151 | 24/18870 | 0.0156442 | 0.6094727 | 0.6090442 | AANAT/T |
| BP | GO:20010 | regulation | 2/151 | 24/18870 | 0.0156442 | 0.6094727 | 0.6090442 | PRKD1/S |
| BP | GO:00422 | ribosome | 7/151 | 325/18870 | 0.0158433 | 0.6094727 | 0.6090442 | EXOSC10 |
| BP | GO:00424 | hormone b | 3/151 | 66/18870 | 0.0159176 | 0.6094727 | 0.6090442 | TPO/AAN |
| BP | GO:00004 | maturation | 2/151 | 25/18870 | 0.0169163 | 0.6094727 | 0.6090442 | EXOSC10 |
| BP | GO:00717 | response t | 4/151 | 123/18870 | 0.0171749 | 0.6094727 | 0.6090442 | TBX2/NP |
| BP | GO:00192 | transmissi | 3/151 | 69/18870 | 0.0179130 | 0.6094727 | 0.6090442 | NPR2/KC |
| BP | GO:00069 | substrate-d | 2/151 | 26/18870 | 0.0182309 | 0.6094727 | 0.6090442 | NCK1/CS |
| BP | GO:00004 | maturation | 2/151 | 28/18870 | 0.0209848 | 0.6094727 | 0.6090442 | MAK16/R |
| BP | GO:00032 | endocardia | 2/151 | 28/18870 | 0.0209848 | 0.6094727 | 0.6090442 | TBX2/FG |
| BP | GO:00351 | embryonic | 2/151 | 28/18870 | 0.0209848 | 0.6094727 | 0.6090442 | GPC3/FG |

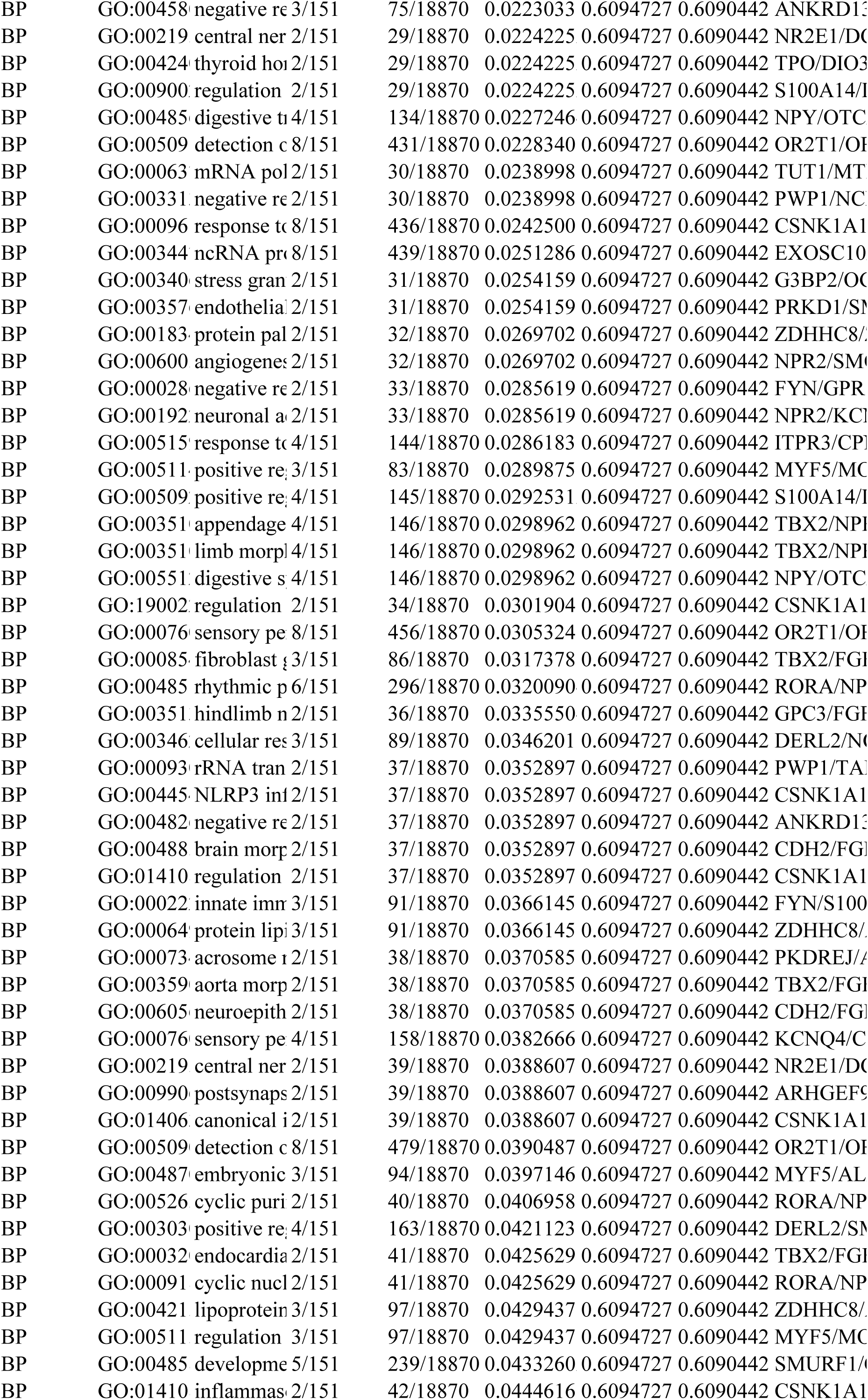

| BP | GO:00458 | negative re | 3/151 | 75/18870 | 0.0223033 | 0.6094727 | 0.6090442 | ANKRD1 |
|---|---|---|---|---|---|---|---|---|
| BP | GO:00219 | central ner | 2/151 | 29/18870 | 0.0224225 | 0.6094727 | 0.6090442 | NR2E1/D |
| BP | GO:00424 | thyroid ho | 2/151 | 29/18870 | 0.0224225 | 0.6094727 | 0.6090442 | TPO/DIO |
| BP | GO:00900 | regulation | 2/151 | 29/18870 | 0.0224225 | 0.6094727 | 0.6090442 | S100A14/ |
| BP | GO:00485 | digestive t | 4/151 | 134/18870 | 0.0227246 | 0.6094727 | 0.6090442 | NPY/OTC |
| BP | GO:00509 | detection c | 8/151 | 431/18870 | 0.0228340 | 0.6094727 | 0.6090442 | OR2T1/O |
| BP | GO:00063 | mRNA pol | 2/151 | 30/18870 | 0.0238998 | 0.6094727 | 0.6090442 | TUT1/MT |
| BP | GO:00331 | negative re | 2/151 | 30/18870 | 0.0238998 | 0.6094727 | 0.6090442 | PWP1/NC |
| BP | GO:00096 | response to | 8/151 | 436/18870 | 0.0242500 | 0.6094727 | 0.6090442 | CSNK1A1 |
| BP | GO:00344 | ncRNA pr | 8/151 | 439/18870 | 0.0251286 | 0.6094727 | 0.6090442 | EXOSC10 |
| BP | GO:00340 | stress gran | 2/151 | 31/18870 | 0.0254159 | 0.6094727 | 0.6090442 | G3BP2/O |
| BP | GO:00357 | endothelia | 2/151 | 31/18870 | 0.0254159 | 0.6094727 | 0.6090442 | PRKD1/S |
| BP | GO:00183 | protein pal | 2/151 | 32/18870 | 0.0269702 | 0.6094727 | 0.6090442 | ZDHHC8/ |
| BP | GO:00600 | angiogenes | 2/151 | 32/18870 | 0.0269702 | 0.6094727 | 0.6090442 | NPR2/SM |
| BP | GO:00028 | negative re | 2/151 | 33/18870 | 0.0285619 | 0.6094727 | 0.6090442 | FYN/GPR |
| BP | GO:00192 | neuronal a | 2/151 | 33/18870 | 0.0285619 | 0.6094727 | 0.6090442 | NPR2/KC |
| BP | GO:00515 | response t | 4/151 | 144/18870 | 0.0286183 | 0.6094727 | 0.6090442 | ITPR3/CP |
| BP | GO:00511 | positive re | 3/151 | 83/18870 | 0.0289875 | 0.6094727 | 0.6090442 | MYF5/MC |
| BP | GO:00509 | positive re | 4/151 | 145/18870 | 0.0292531 | 0.6094727 | 0.6090442 | S100A14/ |
| BP | GO:00351 | appendage | 4/151 | 146/18870 | 0.0298962 | 0.6094727 | 0.6090442 | TBX2/NP |
| BP | GO:00351 | limb morp | 4/151 | 146/18870 | 0.0298962 | 0.6094727 | 0.6090442 | TBX2/NP |
| BP | GO:00551 | digestive s | 4/151 | 146/18870 | 0.0298962 | 0.6094727 | 0.6090442 | NPY/OTC |
| BP | GO:19002 | regulation | 2/151 | 34/18870 | 0.0301904 | 0.6094727 | 0.6090442 | CSNK1A1 |
| BP | GO:00076 | sensory pe | 8/151 | 456/18870 | 0.0305324 | 0.6094727 | 0.6090442 | OR2T1/O |
| BP | GO:00085 | fibroblast | 3/151 | 86/18870 | 0.0317378 | 0.6094727 | 0.6090442 | TBX2/FG |
| BP | GO:00485 | rhythmic p | 6/151 | 296/18870 | 0.0320090 | 0.6094727 | 0.6090442 | RORA/NP |
| BP | GO:00351 | hindlimb n | 2/151 | 36/18870 | 0.0335550 | 0.6094727 | 0.6090442 | GPC3/FG |
| BP | GO:00346 | cellular res | 3/151 | 89/18870 | 0.0346201 | 0.6094727 | 0.6090442 | DERL2/N |
| BP | GO:00093 | rRNA tran | 2/151 | 37/18870 | 0.0352897 | 0.6094727 | 0.6090442 | PWP1/TA |
| BP | GO:00445 | NLRP3 in | 2/151 | 37/18870 | 0.0352897 | 0.6094727 | 0.6090442 | CSNK1A1 |
| BP | GO:00482 | negative re | 2/151 | 37/18870 | 0.0352897 | 0.6094727 | 0.6090442 | ANKRD1 |
| BP | GO:00488 | brain morp | 2/151 | 37/18870 | 0.0352897 | 0.6094727 | 0.6090442 | CDH2/FG |
| BP | GO:01410 | regulation | 2/151 | 37/18870 | 0.0352897 | 0.6094727 | 0.6090442 | CSNK1A1 |
| BP | GO:00022 | innate imm | 3/151 | 91/18870 | 0.0366145 | 0.6094727 | 0.6090442 | FYN/S100 |
| BP | GO:00064 | protein lip | 3/151 | 91/18870 | 0.0366145 | 0.6094727 | 0.6090442 | ZDHHC8/ |
| BP | GO:00073 | acrosome | 2/151 | 38/18870 | 0.0370585 | 0.6094727 | 0.6090442 | PKDREJ/ |
| BP | GO:00359 | aorta morp | 2/151 | 38/18870 | 0.0370585 | 0.6094727 | 0.6090442 | TBX2/FG |
| BP | GO:00605 | neuroepith | 2/151 | 38/18870 | 0.0370585 | 0.6094727 | 0.6090442 | CDH2/FG |
| BP | GO:00076 | sensory pe | 4/151 | 158/18870 | 0.0382666 | 0.6094727 | 0.6090442 | KCNQ4/C |
| BP | GO:00219 | central ner | 2/151 | 39/18870 | 0.0388607 | 0.6094727 | 0.6090442 | NR2E1/D |
| BP | GO:00990 | postsynaps | 2/151 | 39/18870 | 0.0388607 | 0.6094727 | 0.6090442 | ARHGEF |
| BP | GO:01406 | canonical i | 2/151 | 39/18870 | 0.0388607 | 0.6094727 | 0.6090442 | CSNK1A1 |
| BP | GO:00509 | detection c | 8/151 | 479/18870 | 0.0390487 | 0.6094727 | 0.6090442 | OR2T1/O |
| BP | GO:00487 | embryonic | 3/151 | 94/18870 | 0.0397146 | 0.6094727 | 0.6090442 | MYF5/AL |
| BP | GO:00526 | cyclic puri | 2/151 | 40/18870 | 0.0406958 | 0.6094727 | 0.6090442 | RORA/NP |
| BP | GO:00303 | positive re | 4/151 | 163/18870 | 0.0421123 | 0.6094727 | 0.6090442 | DERL2/S |
| BP | GO:00032 | endocardia | 2/151 | 41/18870 | 0.0425629 | 0.6094727 | 0.6090442 | TBX2/FG |
| BP | GO:00091 | cyclic nucl | 2/151 | 41/18870 | 0.0425629 | 0.6094727 | 0.6090442 | RORA/NP |
| BP | GO:00421 | lipoprotein | 3/151 | 97/18870 | 0.0429437 | 0.6094727 | 0.6090442 | ZDHHC8/ |
| BP | GO:00511 | regulation | 3/151 | 97/18870 | 0.0429437 | 0.6094727 | 0.6090442 | MYF5/MC |
| BP | GO:00485 | developme | 5/151 | 239/18870 | 0.0433260 | 0.6094727 | 0.6090442 | SMURF1/ |
| BP | GO:01410 | inflammas | 2/151 | 42/18870 | 0.0444616 | 0.6094727 | 0.6090442 | CSNK1A1 |

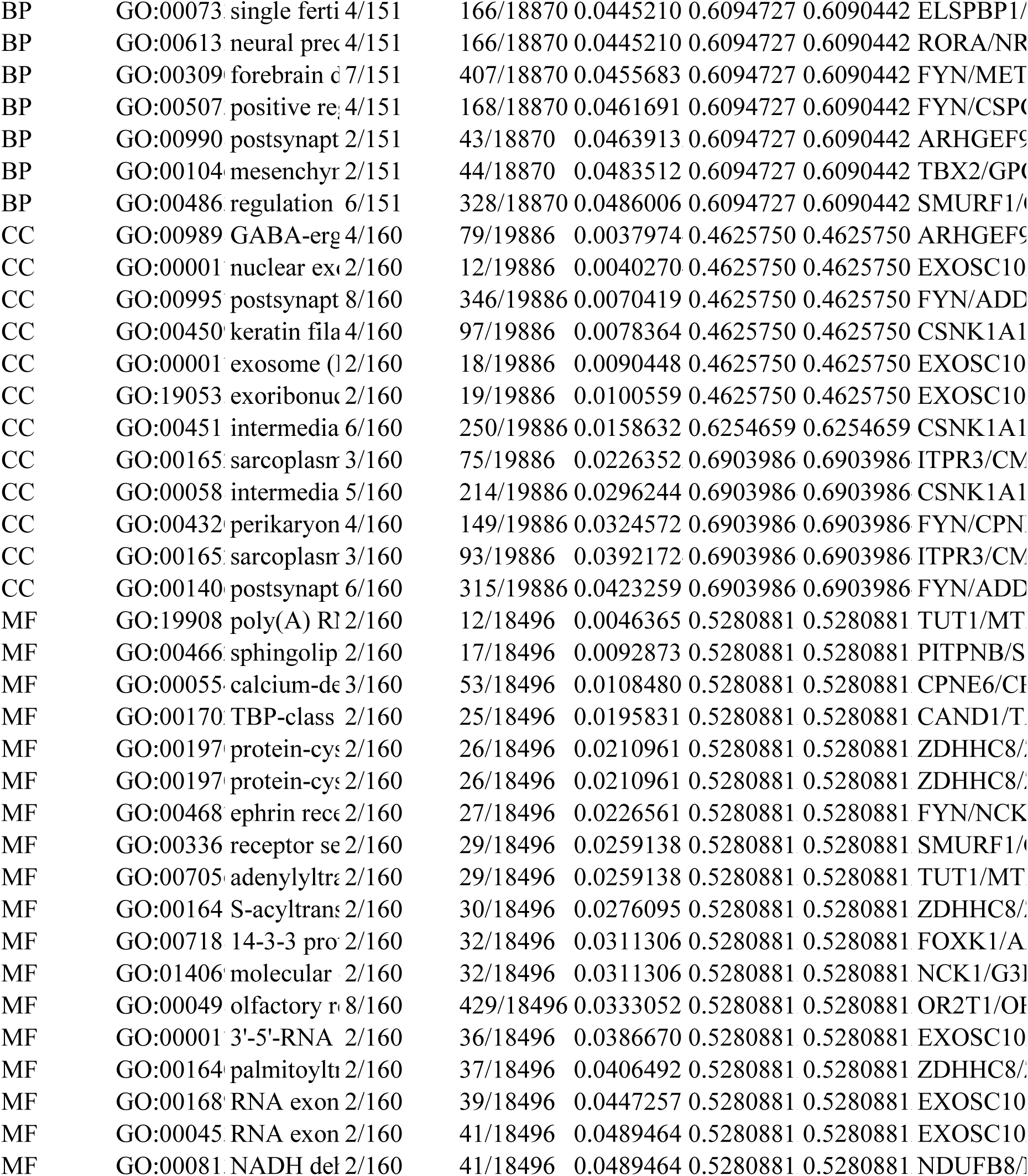

|  |  |  |  |  |  |  |  |  |
|---|---|---|---|---|---|---|---|---|
| BP | GO:00073 | single ferti | 4/151 | 166/18870 | 0.0445210 | 0.6094727 | 0.6090442 | ELSPBP1/ |
| BP | GO:00613 | neural prec | 4/151 | 166/18870 | 0.0445210 | 0.6094727 | 0.6090442 | RORA/NR |
| BP | GO:00309 | forebrain d | 7/151 | 407/18870 | 0.0455683 | 0.6094727 | 0.6090442 | FYN/MET |
| BP | GO:00507 | positive re | 4/151 | 168/18870 | 0.0461691 | 0.6094727 | 0.6090442 | FYN/CSP |
| BP | GO:00990 | postsynapt | 2/151 | 43/18870 | 0.0463913 | 0.6094727 | 0.6090442 | ARHGEF9 |
| BP | GO:00104 | mesenchy | 2/151 | 44/18870 | 0.0483512 | 0.6094727 | 0.6090442 | TBX2/GPC |
| BP | GO:00486 | regulation | 6/151 | 328/18870 | 0.0486006 | 0.6094727 | 0.6090442 | SMURF1/ |
| CC | GO:00989 | GABA-erg | 4/160 | 79/19886 | 0.0037974 | 0.4625750 | 0.4625750 | ARHGEF9 |
| CC | GO:00001 | nuclear ex | 2/160 | 12/19886 | 0.0040270 | 0.4625750 | 0.4625750 | EXOSC10 |
| CC | GO:00995 | postsynapt | 8/160 | 346/19886 | 0.0070419 | 0.4625750 | 0.4625750 | FYN/ADD |
| CC | GO:00450 | keratin fila | 4/160 | 97/19886 | 0.0078364 | 0.4625750 | 0.4625750 | CSNK1A1 |
| CC | GO:00001 | exosome ( | 2/160 | 18/19886 | 0.0090448 | 0.4625750 | 0.4625750 | EXOSC10 |
| CC | GO:19053 | exoribonuc | 2/160 | 19/19886 | 0.0100559 | 0.4625750 | 0.4625750 | EXOSC10 |
| CC | GO:00451 | intermedia | 6/160 | 250/19886 | 0.0158632 | 0.6254659 | 0.6254659 | CSNK1A1 |
| CC | GO:00165 | sarcoplasm | 3/160 | 75/19886 | 0.0226352 | 0.6903986 | 0.6903986 | ITPR3/CM |
| CC | GO:00058 | intermedia | 5/160 | 214/19886 | 0.0296244 | 0.6903986 | 0.6903986 | CSNK1A1 |
| CC | GO:00432 | perikaryon | 4/160 | 149/19886 | 0.0324572 | 0.6903986 | 0.6903986 | FYN/CPN |
| CC | GO:00165 | sarcoplasm | 3/160 | 93/19886 | 0.0392172 | 0.6903986 | 0.6903986 | ITPR3/CM |
| CC | GO:00140 | postsynapt | 6/160 | 315/19886 | 0.0423259 | 0.6903986 | 0.6903986 | FYN/ADD |
| MF | GO:19908 | poly(A) R | 2/160 | 12/18496 | 0.0046365 | 0.5280881 | 0.5280881 | TUT1/MT |
| MF | GO:00466 | sphingolip | 2/160 | 17/18496 | 0.0092873 | 0.5280881 | 0.5280881 | PITPNB/S |
| MF | GO:00055 | calcium-de | 3/160 | 53/18496 | 0.0108480 | 0.5280881 | 0.5280881 | CPNE6/CP |
| MF | GO:00170 | TBP-class | 2/160 | 25/18496 | 0.0195831 | 0.5280881 | 0.5280881 | CAND1/T |
| MF | GO:00197 | protein-cy | 2/160 | 26/18496 | 0.0210961 | 0.5280881 | 0.5280881 | ZDHHC8/ |
| MF | GO:00197 | protein-cy | 2/160 | 26/18496 | 0.0210961 | 0.5280881 | 0.5280881 | ZDHHC8/ |
| MF | GO:00468 | ephrin rec | 2/160 | 27/18496 | 0.0226561 | 0.5280881 | 0.5280881 | FYN/NCK |
| MF | GO:00336 | receptor se | 2/160 | 29/18496 | 0.0259138 | 0.5280881 | 0.5280881 | SMURF1/ |
| MF | GO:00705 | adenylyltr | 2/160 | 29/18496 | 0.0259138 | 0.5280881 | 0.5280881 | TUT1/MT |
| MF | GO:00164 | S-acyltran | 2/160 | 30/18496 | 0.0276095 | 0.5280881 | 0.5280881 | ZDHHC8/ |
| MF | GO:00718 | 14-3-3 pro | 2/160 | 32/18496 | 0.0311306 | 0.5280881 | 0.5280881 | FOXK1/A |
| MF | GO:01406 | molecular | 2/160 | 32/18496 | 0.0311306 | 0.5280881 | 0.5280881 | NCK1/G3 |
| MF | GO:00049 | olfactory r | 8/160 | 429/18496 | 0.0333052 | 0.5280881 | 0.5280881 | OR2T1/OR |
| MF | GO:00001 | 3'-5'-RNA | 2/160 | 36/18496 | 0.0386670 | 0.5280881 | 0.5280881 | EXOSC10 |
| MF | GO:00164 | palmitoylt | 2/160 | 37/18496 | 0.0406492 | 0.5280881 | 0.5280881 | ZDHHC8/ |
| MF | GO:00168 | RNA exon | 2/160 | 39/18496 | 0.0447257 | 0.5280881 | 0.5280881 | EXOSC10 |
| MF | GO:00045 | RNA exon | 2/160 | 41/18496 | 0.0489464 | 0.5280881 | 0.5280881 | EXOSC10 |
| MF | GO:00081 | NADH del | 2/160 | 41/18496 | 0.0489464 | 0.5280881 | 0.5280881 | NDUFB8/ |

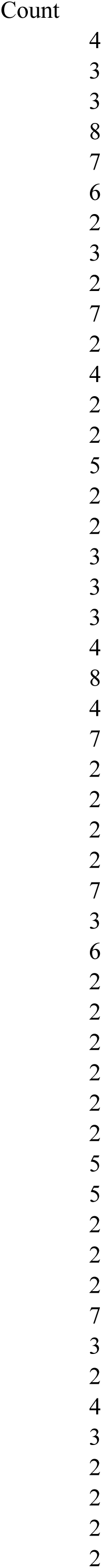

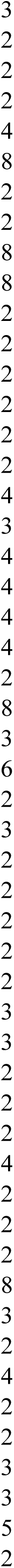

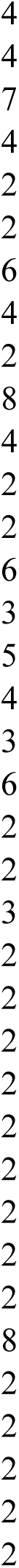

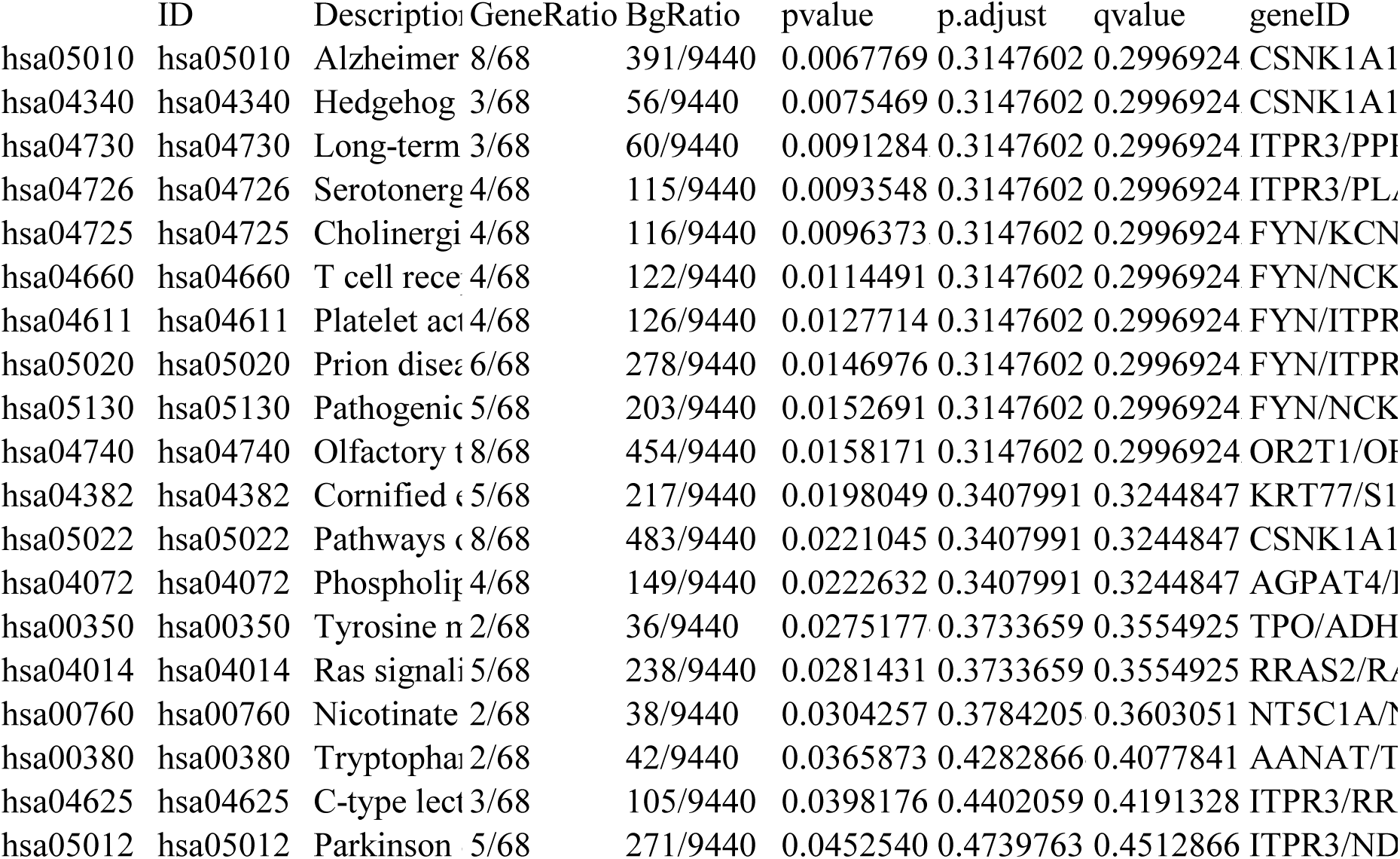

|  | ID | Descriptio | GeneRatio | BgRatio | pvalue | p.adjust | qvalue | geneID |
|---|---|---|---|---|---|---|---|---|
| hsa05010 | hsa05010 | Alzheimer | 8/68 | 391/9440 | 0.0067769 | 0.3147602 | 0.2996924 | CSNK1A1 |
| hsa04340 | hsa04340 | Hedgehog | 3/68 | 56/9440 | 0.0075469 | 0.3147602 | 0.2996924 | CSNK1A1 |
| hsa04730 | hsa04730 | Long-term | 3/68 | 60/9440 | 0.0091284 | 0.3147602 | 0.2996924 | ITPR3/PP |
| hsa04726 | hsa04726 | Serotonerg | 4/68 | 115/9440 | 0.0093548 | 0.3147602 | 0.2996924 | ITPR3/PL |
| hsa04725 | hsa04725 | Cholinergi | 4/68 | 116/9440 | 0.0096373 | 0.3147602 | 0.2996924 | FYN/KCN |
| hsa04660 | hsa04660 | T cell rece | 4/68 | 122/9440 | 0.0114491 | 0.3147602 | 0.2996924 | FYN/NCK |
| hsa04611 | hsa04611 | Platelet ac | 4/68 | 126/9440 | 0.0127714 | 0.3147602 | 0.2996924 | FYN/ITPR |
| hsa05020 | hsa05020 | Prion dise | 6/68 | 278/9440 | 0.0146976 | 0.3147602 | 0.2996924 | FYN/ITPR |
| hsa05130 | hsa05130 | Pathogeni | 5/68 | 203/9440 | 0.0152691 | 0.3147602 | 0.2996924 | FYN/NCK |
| hsa04740 | hsa04740 | Olfactory | 8/68 | 454/9440 | 0.0158171 | 0.3147602 | 0.2996924 | OR2T1/O |
| hsa04382 | hsa04382 | Cornified | 5/68 | 217/9440 | 0.0198049 | 0.3407991 | 0.3244847 | KRT77/S1 |
| hsa05022 | hsa05022 | Pathways | 8/68 | 483/9440 | 0.0221045 | 0.3407991 | 0.3244847 | CSNK1A1 |
| hsa04072 | hsa04072 | Phospholi | 4/68 | 149/9440 | 0.0222632 | 0.3407991 | 0.3244847 | AGPAT4/ |
| hsa00350 | hsa00350 | Tyrosine | 2/68 | 36/9440 | 0.0275177 | 0.3733659 | 0.3554925 | TPO/ADH |
| hsa04014 | hsa04014 | Ras signal | 5/68 | 238/9440 | 0.0281431 | 0.3733659 | 0.3554925 | RRAS2/R |
| hsa00760 | hsa00760 | Nicotinate | 2/68 | 38/9440 | 0.0304257 | 0.3784205 | 0.3603051 | NT5C1A/ |
| hsa00380 | hsa00380 | Tryptopha | 2/68 | 42/9440 | 0.0365873 | 0.4282866 | 0.4077841 | AANAT/T |
| hsa04625 | hsa04625 | C-type lec | 3/68 | 105/9440 | 0.0398176 | 0.4402059 | 0.4191328 | ITPR3/RR |
| hsa05012 | hsa05012 | Parkinson | 5/68 | 271/9440 | 0.0452540 | 0.4739763 | 0.4512866 | ITPR3/ND |

| Count |
|---|
| 8 |
| 3 |
| 3 |
| 4 |
| 4 |
| 4 |
| 4 |
| 6 |
| 5 |
| 8 |
| 5 |
| 8 |
| 4 |
| 2 |
| 5 |
| 2 |
| 2 |
| 3 |
| 5 |

| ID | Descriptio | setSize | enrichmen | NES | pvalue | p.adjust | qvalue | rank |
|---|---|---|---|---|---|---|---|---|
| KEGG_M | KEGG_M | 18 | 0.6898124 | 2.4997770 | 5.8791232 | 0.0021341 | 0.0018256 | 3107 |
| KEGG_M | KEGG_M | 15 | 0.6961287 | 2.4033257 | 2.1515941 | 0.0025013 | 0.0021397 | 3107 |
| KEGG_M | KEGG_M | 29 | 0.5543296 | 2.2978562 | 2.7563495 | 0.0025013 | 0.0021397 | 3772 |
| KEGG_M | KEGG_M | 17 | 0.6306572 | 2.2642624 | 2.0016271 | 0.0066053 | 0.0056505 | 4871 |
| KEGG_M | KEGG_M | 21 | 0.5817228 | 2.2114060 | 1.8485571 | 0.0066053 | 0.0056505 | 5309 |
| KEGG_M | KEGG_M | 22 | 0.5686142 | 2.2002874 | 3.9080134 | 0.0118217 | 0.0101128 | 4545 |
| KEGG_M | KEGG_M | 37 | 0.4998669 | 2.1875528 | 2.5748372 | 0.0025013 | 0.0021397 | 4468 |
| KEGG_M | KEGG_M | 37 | 0.4946167 | 2.1645764 | 3.5521327 | 0.0025788 | 0.0022060 | 4468 |
| KEGG_M | KEGG_M | 36 | 0.4876289 | 2.1325992 | 1.3017810 | 0.0066053 | 0.0056505 | 4468 |
| KEGG_M | KEGG_M | 19 | 0.5633952 | 2.0809260 | 9.8642082 | 0.0275439 | 0.0235622 | 3694 |
| KEGG_M | KEGG_M | 11 | 0.6641928 | 2.0722142 | 0.0016537 | 0.0414941 | 0.0354959 | 4203 |
| KEGG_M | KEGG_M | 37 | 0.4685547 | 2.0505224 | 1.7243183 | 0.0066053 | 0.0056505 | 4468 |
| KEGG_M | KEGG_M | 43 | 0.4512144 | 2.0322269 | 1.4747793 | 0.0066053 | 0.0056505 | 4468 |
| KEGG_M | KEGG_M | 46 | 0.4346672 | 2.0137989 | 1.6746638 | 0.0066053 | 0.0056505 | 6111 |
| KEGG_M | KEGG_M | 15 | 0.5684114 | 1.9623927 | 0.0021898 | 0.0496829 | 0.0425010 | 3902 |
| KEGG_M | KEGG_M | 24 | 0.4873666 | 1.9328929 | 0.0017146 | 0.0414941 | 0.0354959 | 4315 |

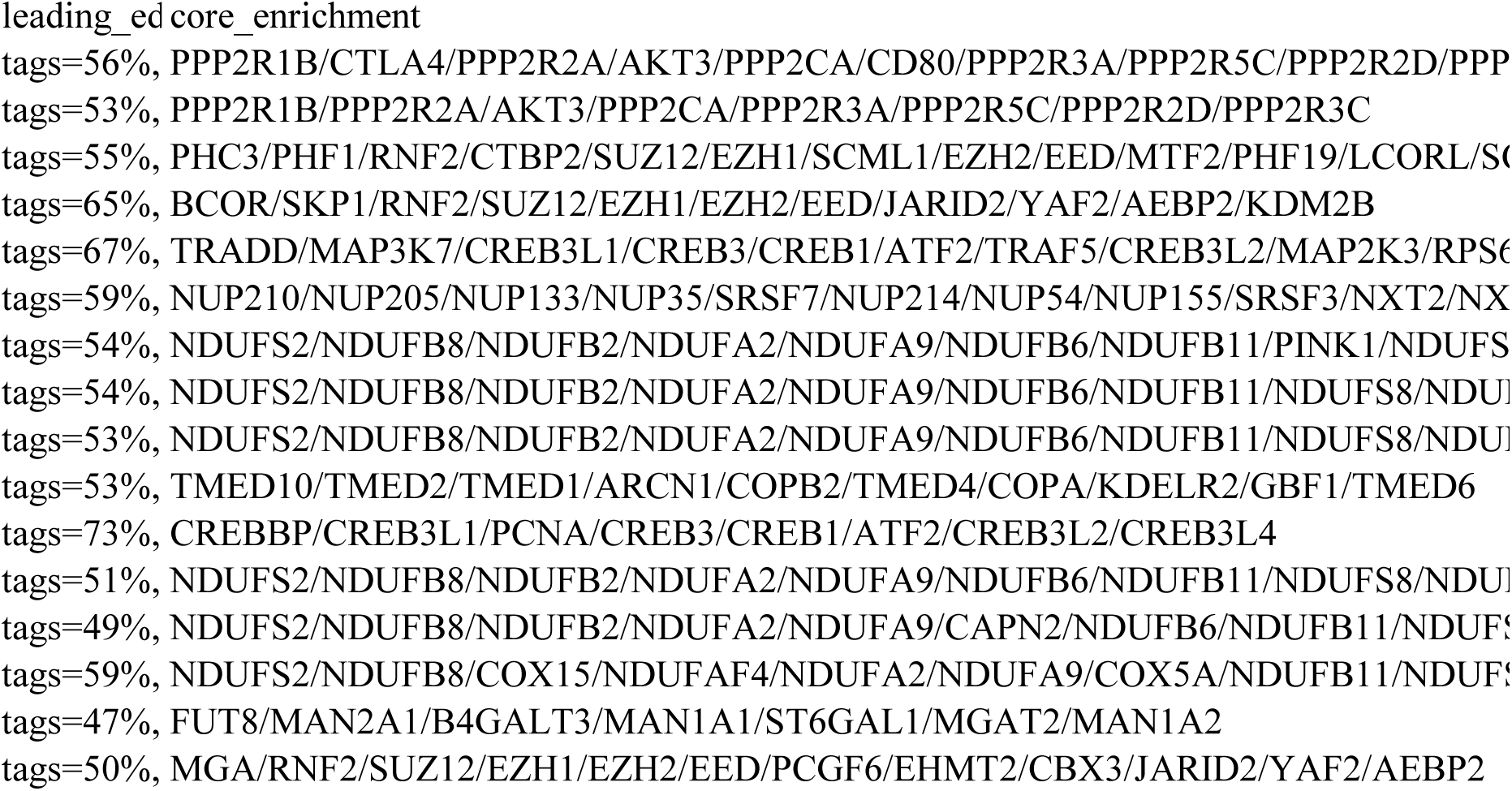

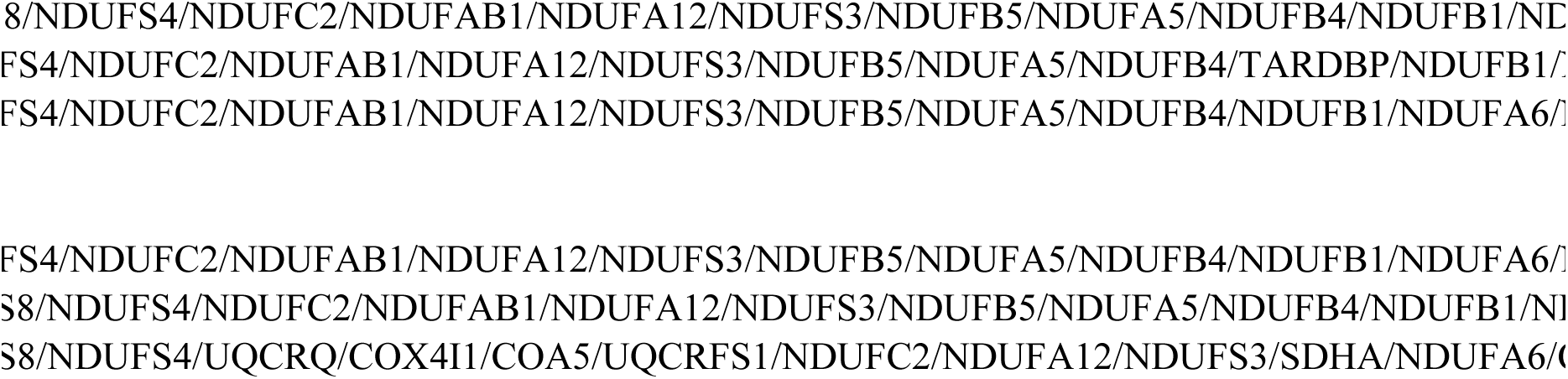

|  |
|---|
| COX10/NDUFAF2/NDUFV1/NDUFB7/NDUFA10/NDUFV2/NDUFS1/NDUFAF1 |

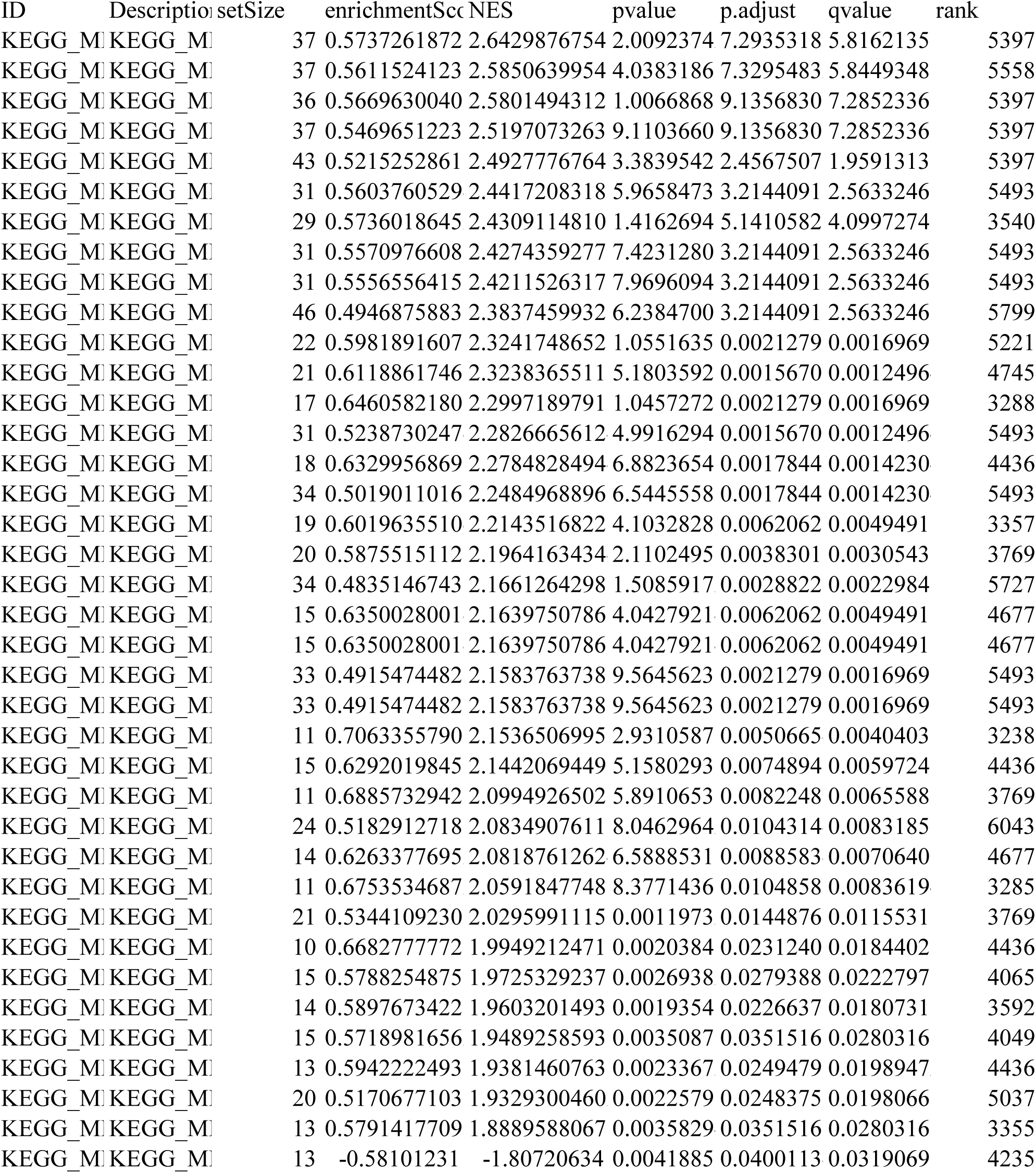

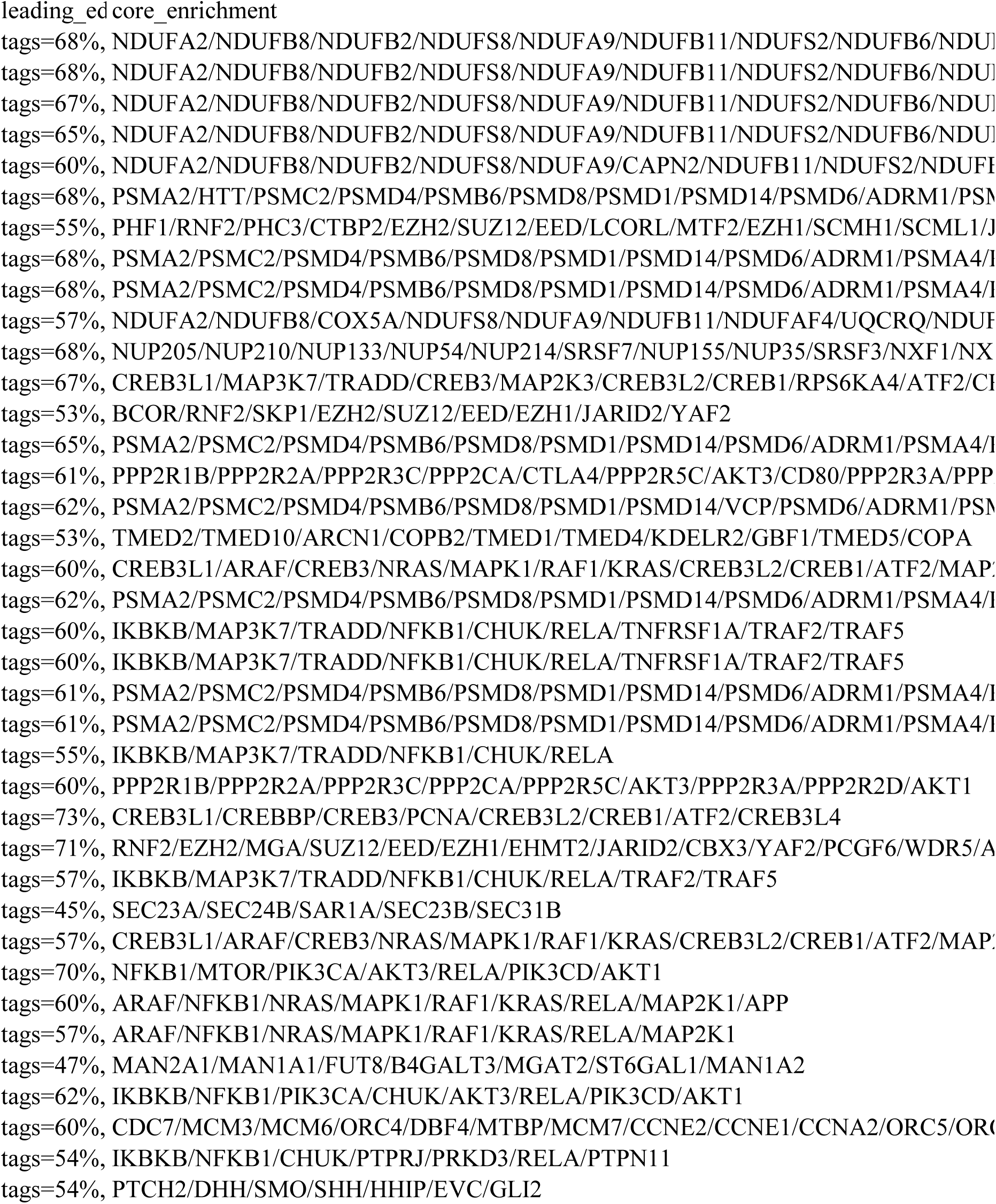

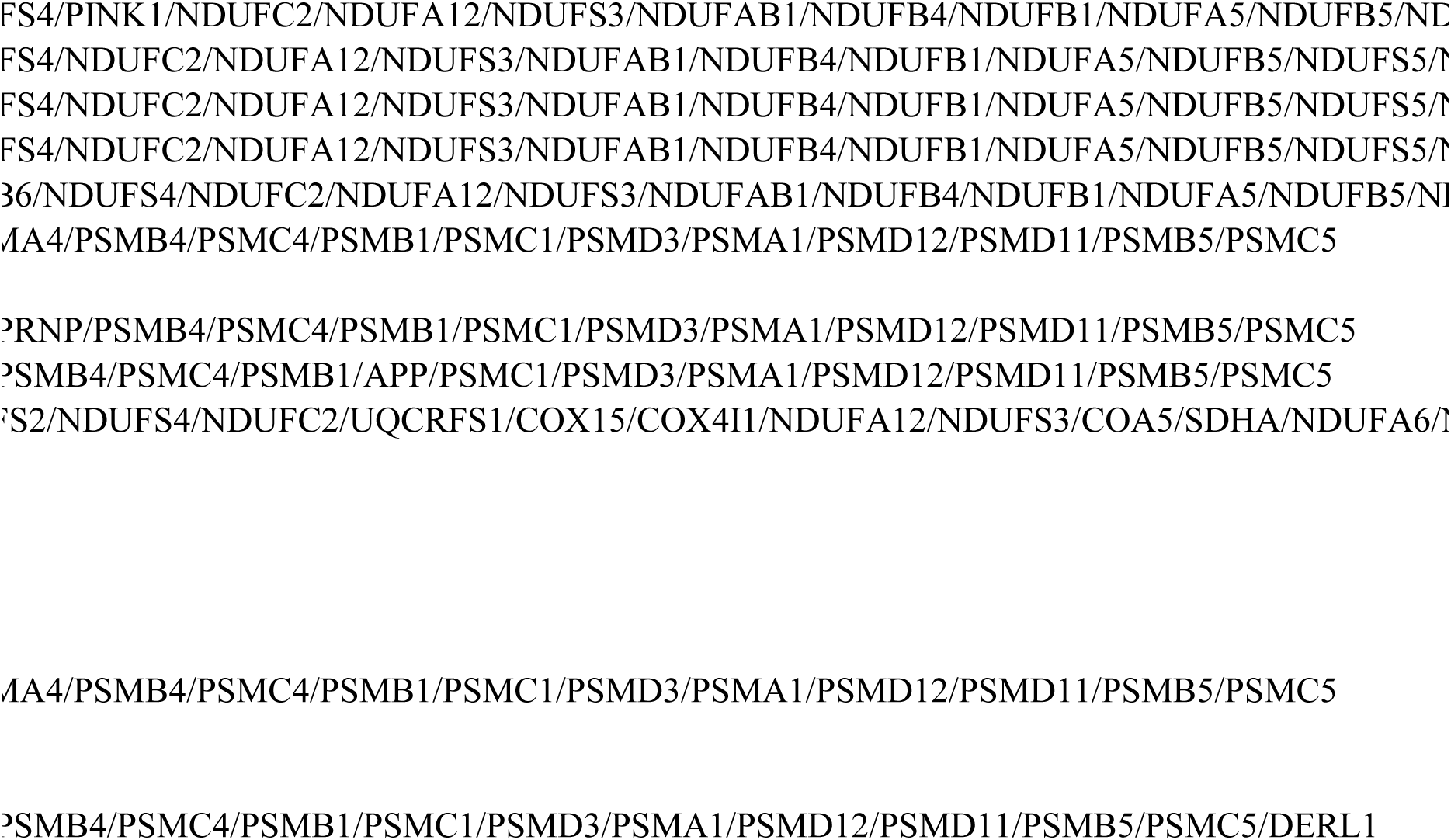

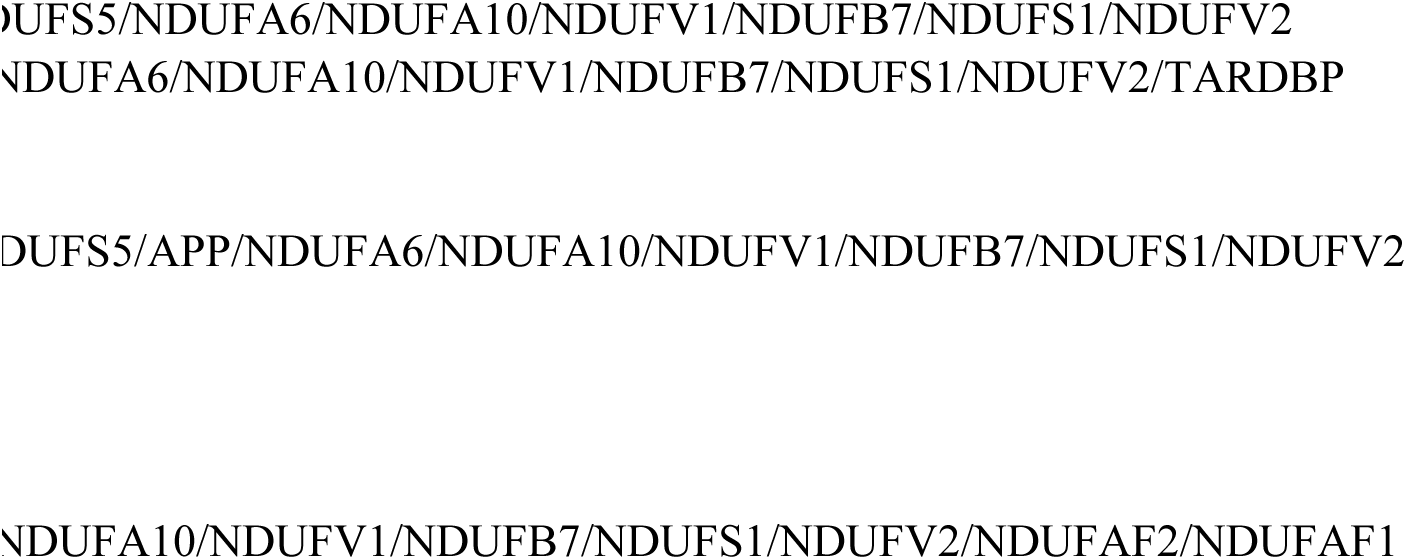

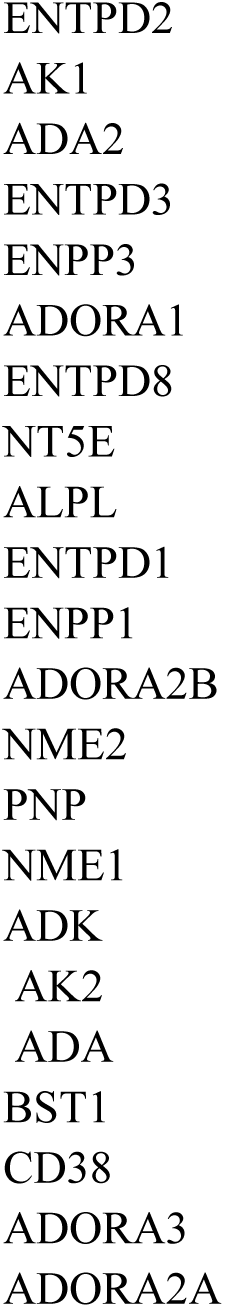

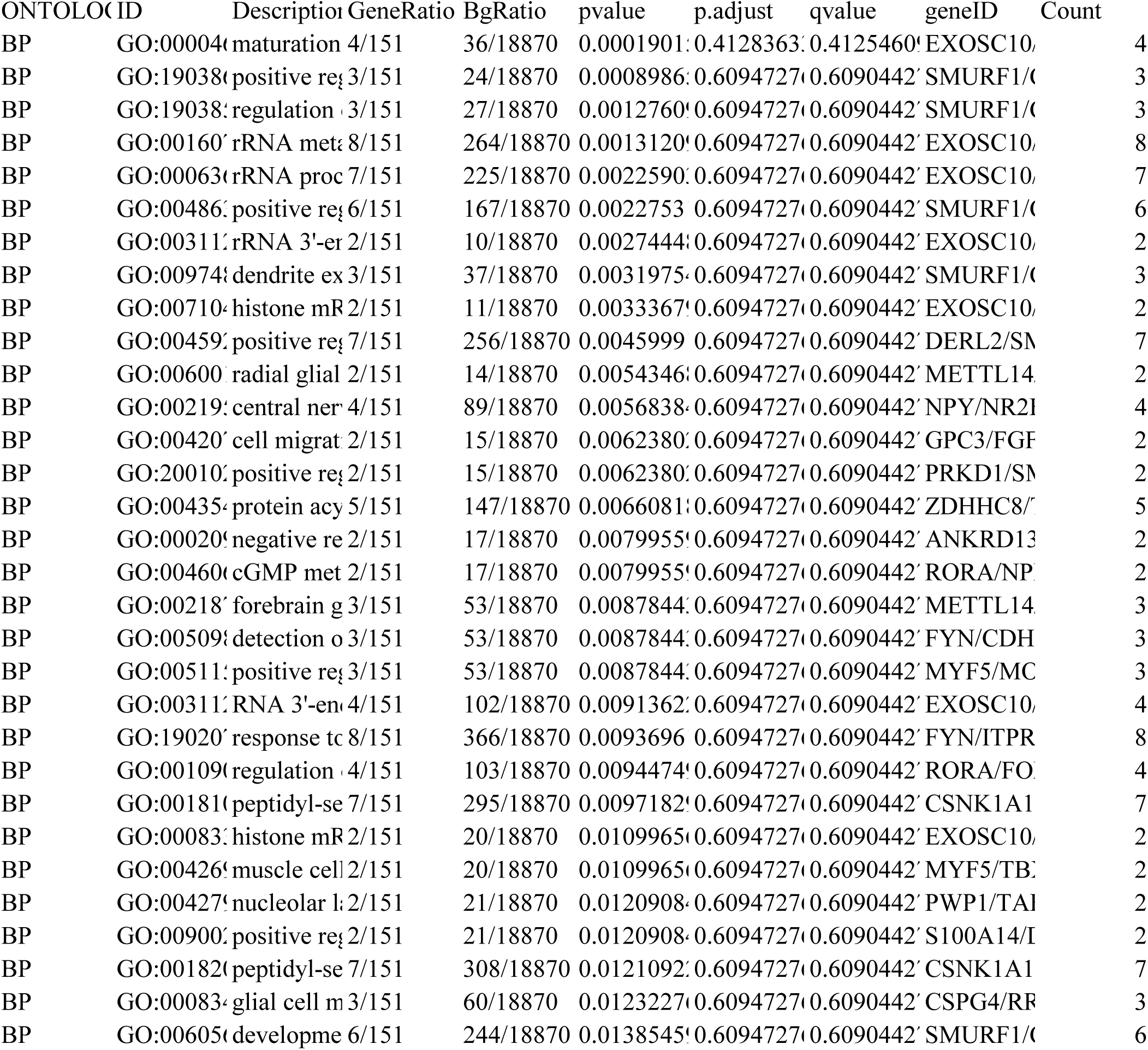

| ONTOLOG | ID | Descriptio | GeneRatio | BgRatio | pvalue | p.adjust | qvalue | geneID | Count |
|---|---|---|---|---|---|---|---|---|---|
| BP | GO:00004 | maturation | 4/151 | 36/18870 | 0.0001901 | 0.4128363 | 0.4125460 | EXOSC10/ | 4 |
| BP | GO:19038 | positive re | 3/151 | 24/18870 | 0.0008986 | 0.6094727 | 0.6090442 | SMURF1/ | 3 |
| BP | GO:19038 | regulation | 3/151 | 27/18870 | 0.0012760 | 0.6094727 | 0.6090442 | SMURF1/ | 3 |
| BP | GO:00160 | rRNA met | 8/151 | 264/18870 | 0.0013120 | 0.6094727 | 0.6090442 | EXOSC10/ | 8 |
| BP | GO:00063 | rRNA proc | 7/151 | 225/18870 | 0.0022590 | 0.6094727 | 0.6090442 | EXOSC10/ | 7 |
| BP | GO:00486 | positive re | 6/151 | 167/18870 | 0.0022753 | 0.6094727 | 0.6090442 | SMURF1/ | 6 |
| BP | GO:00311 | rRNA 3'-e | 2/151 | 10/18870 | 0.0027444 | 0.6094727 | 0.6090442 | EXOSC10/ | 2 |
| BP | GO:00974 | dendrite ex | 3/151 | 37/18870 | 0.0031975 | 0.6094727 | 0.6090442 | SMURF1/ | 3 |
| BP | GO:00710 | histone m | 2/151 | 11/18870 | 0.0033367 | 0.6094727 | 0.6090442 | EXOSC10/ | 2 |
| BP | GO:00459 | positive re | 7/151 | 256/18870 | 0.0045999 | 0.6094727 | 0.6090442 | DERL2/S | 7 |
| BP | GO:00600 | radial glial | 2/151 | 14/18870 | 0.0054346 | 0.6094727 | 0.6090442 | METTL14 | 2 |
| BP | GO:00219 | central ner | 4/151 | 89/18870 | 0.0056838 | 0.6094727 | 0.6090442 | NPY/NR2 | 4 |
| BP | GO:00420 | cell migrat | 2/151 | 15/18870 | 0.0062380 | 0.6094727 | 0.6090442 | GPC3/FG | 2 |
| BP | GO:20010 | positive re | 2/151 | 15/18870 | 0.0062380 | 0.6094727 | 0.6090442 | PRKD1/S | 2 |
| BP | GO:00435 | protein acy | 5/151 | 147/18870 | 0.0066081 | 0.6094727 | 0.6090442 | ZDHHC8/ | 5 |
| BP | GO:00020 | negative re | 2/151 | 17/18870 | 0.0079955 | 0.6094727 | 0.6090442 | ANKRD1 | 2 |
| BP | GO:00460 | cGMP met | 2/151 | 17/18870 | 0.0079955 | 0.6094727 | 0.6090442 | RORA/NP | 2 |
| BP | GO:00218 | forebrain g | 3/151 | 53/18870 | 0.0087844 | 0.6094727 | 0.6090442 | METTL14 | 3 |
| BP | GO:00509 | detection o | 3/151 | 53/18870 | 0.0087844 | 0.6094727 | 0.6090442 | FYN/CDH | 3 |
| BP | GO:00511 | positive re | 3/151 | 53/18870 | 0.0087844 | 0.6094727 | 0.6090442 | MYF5/MC | 3 |
| BP | GO:00311 | RNA 3'-en | 4/151 | 102/18870 | 0.0091362 | 0.6094727 | 0.6090442 | EXOSC10/ | 4 |
| BP | GO:19020 | response t | 8/151 | 366/18870 | 0.0093696 | 0.6094727 | 0.6090442 | FYN/ITPR | 8 |
| BP | GO:00109 | regulation | 4/151 | 103/18870 | 0.0094474 | 0.6094727 | 0.6090442 | RORA/FO | 4 |
| BP | GO:00181 | peptidyl-se | 7/151 | 295/18870 | 0.0097182 | 0.6094727 | 0.6090442 | CSNK1A1 | 7 |
| BP | GO:00083 | histone m | 2/151 | 20/18870 | 0.0109965 | 0.6094727 | 0.6090442 | EXOSC10/ | 2 |
| BP | GO:00426 | muscle cel | 2/151 | 20/18870 | 0.0109965 | 0.6094727 | 0.6090442 | MYF5/TB | 2 |
| BP | GO:00427 | nucleolar l | 2/151 | 21/18870 | 0.0120908 | 0.6094727 | 0.6090442 | PWP1/TA | 2 |
| BP | GO:00900 | positive re | 2/151 | 21/18870 | 0.0120908 | 0.6094727 | 0.6090442 | S100A14/ | 2 |
| BP | GO:00182 | peptidyl-se | 7/151 | 308/18870 | 0.0121092 | 0.6094727 | 0.6090442 | CSNK1A1 | 7 |
| BP | GO:00083 | glial cell m | 3/151 | 60/18870 | 0.0123227 | 0.6094727 | 0.6090442 | CSPG4/R | 3 |
| BP | GO:00605 | developme | 6/151 | 244/18870 | 0.0138545 | 0.6094727 | 0.6090442 | SMURF1/ | 6 |

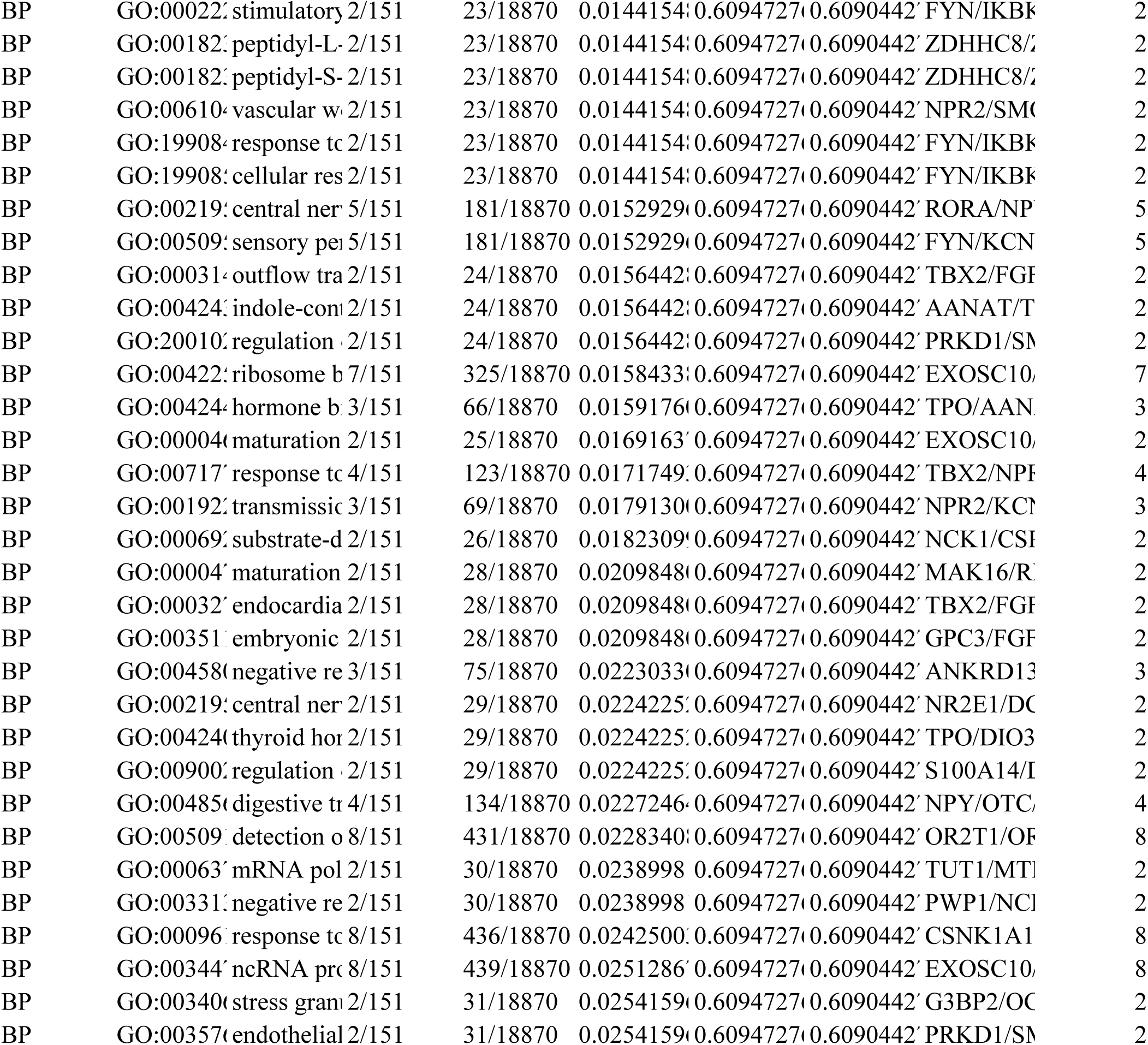

|  |  |  |  |  |  |  |  |  |  |
|---|---|---|---|---|---|---|---|---|---|
| BP | GO:00022 | stimulatory | 2/151 | 23/18870 | 0.0144154 | 0.6094727 | 0.6090442 | FYN/IKBK | 2 |
| BP | GO:00182 | peptidyl-L- | 2/151 | 23/18870 | 0.0144154 | 0.6094727 | 0.6090442 | ZDHHC8/Z | 2 |
| BP | GO:00182 | peptidyl-S- | 2/151 | 23/18870 | 0.0144154 | 0.6094727 | 0.6090442 | ZDHHC8/Z | 2 |
| BP | GO:00610 | vascular w | 2/151 | 23/18870 | 0.0144154 | 0.6094727 | 0.6090442 | NPR2/SM | 2 |
| BP | GO:19908 | response t | 2/151 | 23/18870 | 0.0144154 | 0.6094727 | 0.6090442 | FYN/IKBK | 2 |
| BP | GO:19908 | cellular res | 2/151 | 23/18870 | 0.0144154 | 0.6094727 | 0.6090442 | FYN/IKBK | 2 |
| BP | GO:00219 | central ner | 5/151 | 181/18870 | 0.0152929 | 0.6094727 | 0.6090442 | RORA/NP | 5 |
| BP | GO:00509 | sensory pe | 5/151 | 181/18870 | 0.0152929 | 0.6094727 | 0.6090442 | FYN/KCN | 5 |
| BP | GO:00031 | outflow tra | 2/151 | 24/18870 | 0.0156442 | 0.6094727 | 0.6090442 | TBX2/FG | 2 |
| BP | GO:00424 | indole-con | 2/151 | 24/18870 | 0.0156442 | 0.6094727 | 0.6090442 | AANAT/T | 2 |
| BP | GO:20010 | regulation | 2/151 | 24/18870 | 0.0156442 | 0.6094727 | 0.6090442 | PRKD1/SM | 2 |
| BP | GO:00422 | ribosome b | 7/151 | 325/18870 | 0.0158433 | 0.6094727 | 0.6090442 | EXOSC10/ | 7 |
| BP | GO:00424 | hormone b | 3/151 | 66/18870 | 0.0159176 | 0.6094727 | 0.6090442 | TPO/AAN | 3 |
| BP | GO:00004 | maturation | 2/151 | 25/18870 | 0.0169163 | 0.6094727 | 0.6090442 | EXOSC10/ | 2 |
| BP | GO:00717 | response t | 4/151 | 123/18870 | 0.0171749 | 0.6094727 | 0.6090442 | TBX2/NP | 4 |
| BP | GO:00192 | transmissi | 3/151 | 69/18870 | 0.0179130 | 0.6094727 | 0.6090442 | NPR2/KC | 3 |
| BP | GO:00069 | substrate-d | 2/151 | 26/18870 | 0.0182309 | 0.6094727 | 0.6090442 | NCK1/CS | 2 |
| BP | GO:00004 | maturation | 2/151 | 28/18870 | 0.0209848 | 0.6094727 | 0.6090442 | MAK16/R | 2 |
| BP | GO:00032 | endocardia | 2/151 | 28/18870 | 0.0209848 | 0.6094727 | 0.6090442 | TBX2/FG | 2 |
| BP | GO:00351 | embryonic | 2/151 | 28/18870 | 0.0209848 | 0.6094727 | 0.6090442 | GPC3/FG | 2 |
| BP | GO:00458 | negative re | 3/151 | 75/18870 | 0.0223033 | 0.6094727 | 0.6090442 | ANKRD13 | 3 |
| BP | GO:00219 | central ner | 2/151 | 29/18870 | 0.0224225 | 0.6094727 | 0.6090442 | NR2E1/D | 2 |
| BP | GO:00424 | thyroid hor | 2/151 | 29/18870 | 0.0224225 | 0.6094727 | 0.6090442 | TPO/DIO3 | 2 |
| BP | GO:00900 | regulation | 2/151 | 29/18870 | 0.0224225 | 0.6094727 | 0.6090442 | S100A14/ | 2 |
| BP | GO:00485 | digestive tr | 4/151 | 134/18870 | 0.0227246 | 0.6094727 | 0.6090442 | NPY/OTC/ | 4 |
| BP | GO:00509 | detection o | 8/151 | 431/18870 | 0.0228340 | 0.6094727 | 0.6090442 | OR2T1/O | 8 |
| BP | GO:00063 | mRNA pol | 2/151 | 30/18870 | 0.0238998 | 0.6094727 | 0.6090442 | TUT1/MT | 2 |
| BP | GO:00331 | negative re | 2/151 | 30/18870 | 0.0238998 | 0.6094727 | 0.6090442 | PWP1/NC | 2 |
| BP | GO:00096 | response t | 8/151 | 436/18870 | 0.0242500 | 0.6094727 | 0.6090442 | CSNK1A1 | 8 |
| BP | GO:00344 | ncRNA pr | 8/151 | 439/18870 | 0.0251286 | 0.6094727 | 0.6090442 | EXOSC10/ | 8 |
| BP | GO:00340 | stress gran | 2/151 | 31/18870 | 0.0254159 | 0.6094727 | 0.6090442 | G3BP2/O | 2 |
| BP | GO:00357 | endothelial | 2/151 | 31/18870 | 0.0254159 | 0.6094727 | 0.6090442 | PRKD1/SM | 2 |

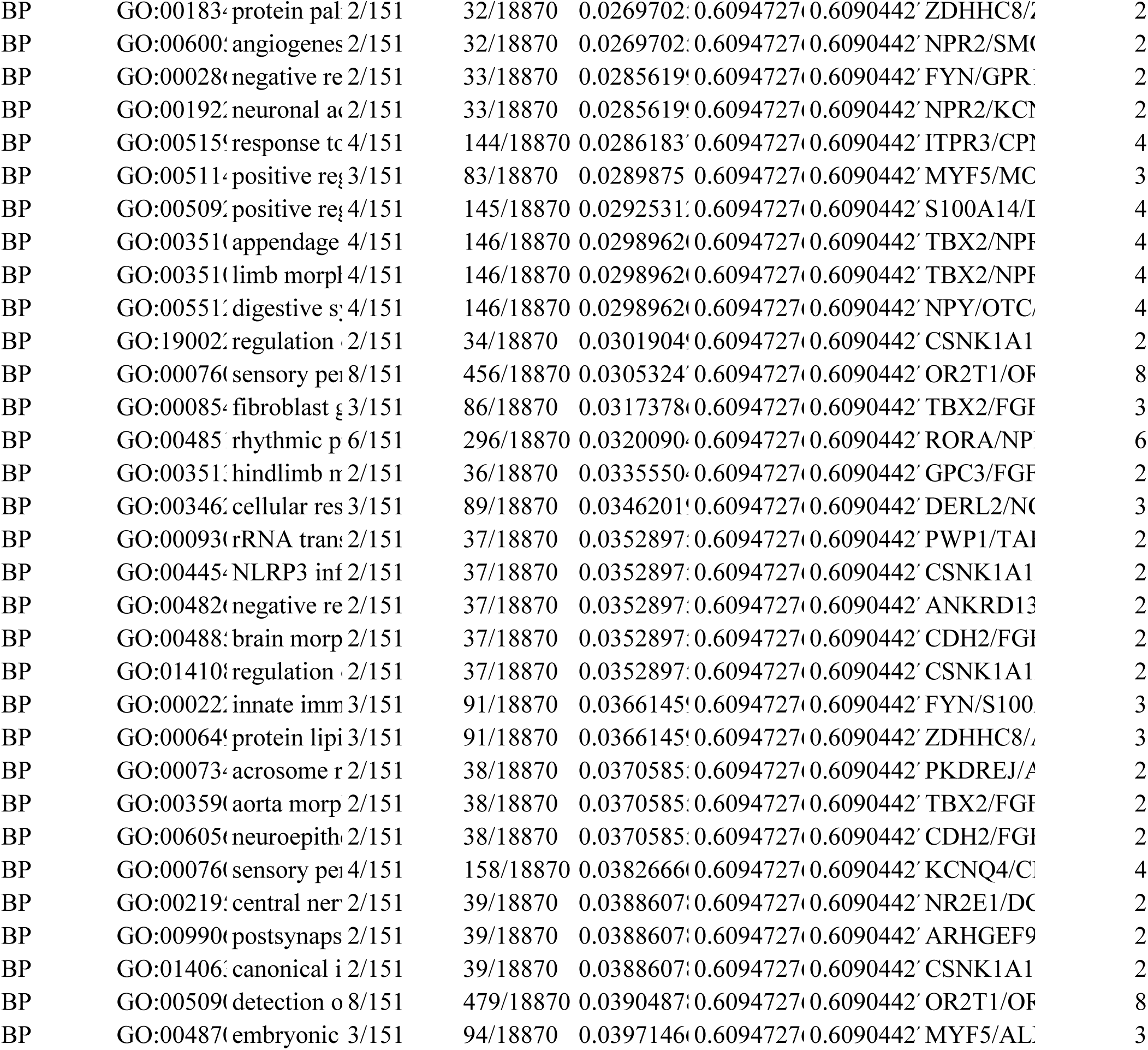

|  |  |  |  |  |  |  |  |  |  |
|---|---|---|---|---|---|---|---|---|---|
| BP | GO:00183 | protein pal | 2/151 | 32/18870 | 0.0269702 | 0.6094727 | 0.6090442 | ZDHHC8/Z | 2 |
| BP | GO:00600 | angiogenes | 2/151 | 32/18870 | 0.0269702 | 0.6094727 | 0.6090442 | NPR2/SM | 2 |
| BP | GO:00028 | negative re | 2/151 | 33/18870 | 0.0285619 | 0.6094727 | 0.6090442 | FYN/GPR | 2 |
| BP | GO:00192 | neuronal a | 2/151 | 33/18870 | 0.0285619 | 0.6094727 | 0.6090442 | NPR2/KC | 2 |
| BP | GO:00515 | response to | 4/151 | 144/18870 | 0.0286183 | 0.6094727 | 0.6090442 | ITPR3/CP | 4 |
| BP | GO:00511 | positive re | 3/151 | 83/18870 | 0.0289875 | 0.6094727 | 0.6090442 | MYF5/MC | 3 |
| BP | GO:00509 | positive re | 4/151 | 145/18870 | 0.0292531 | 0.6094727 | 0.6090442 | S100A14/I | 4 |
| BP | GO:00351 | appendage | 4/151 | 146/18870 | 0.0298962 | 0.6094727 | 0.6090442 | TBX2/NP | 4 |
| BP | GO:00351 | limb morph | 4/151 | 146/18870 | 0.0298962 | 0.6094727 | 0.6090442 | TBX2/NP | 4 |
| BP | GO:00551 | digestive sy | 4/151 | 146/18870 | 0.0298962 | 0.6094727 | 0.6090442 | NPY/OTC | 4 |
| BP | GO:19002 | regulation | 2/151 | 34/18870 | 0.0301904 | 0.6094727 | 0.6090442 | CSNK1A1 | 2 |
| BP | GO:00076 | sensory pe | 8/151 | 456/18870 | 0.0305324 | 0.6094727 | 0.6090442 | OR2T1/O | 8 |
| BP | GO:00085 | fibroblast | 3/151 | 86/18870 | 0.0317378 | 0.6094727 | 0.6090442 | TBX2/FG | 3 |
| BP | GO:00485 | rhythmic p | 6/151 | 296/18870 | 0.0320090 | 0.6094727 | 0.6090442 | RORA/NP | 6 |
| BP | GO:00351 | hindlimb m | 2/151 | 36/18870 | 0.0335550 | 0.6094727 | 0.6090442 | GPC3/FG | 2 |
| BP | GO:00346 | cellular res | 3/151 | 89/18870 | 0.0346201 | 0.6094727 | 0.6090442 | DERL2/N | 3 |
| BP | GO:00093 | rRNA trans | 2/151 | 37/18870 | 0.0352897 | 0.6094727 | 0.6090442 | PWP1/TA | 2 |
| BP | GO:00445 | NLRP3 inf | 2/151 | 37/18870 | 0.0352897 | 0.6094727 | 0.6090442 | CSNK1A1 | 2 |
| BP | GO:00482 | negative re | 2/151 | 37/18870 | 0.0352897 | 0.6094727 | 0.6090442 | ANKRD13 | 2 |
| BP | GO:00488 | brain morp | 2/151 | 37/18870 | 0.0352897 | 0.6094727 | 0.6090442 | CDH2/FG | 2 |
| BP | GO:01410 | regulation | 2/151 | 37/18870 | 0.0352897 | 0.6094727 | 0.6090442 | CSNK1A1 | 2 |
| BP | GO:00022 | innate imm | 3/151 | 91/18870 | 0.0366145 | 0.6094727 | 0.6090442 | FYN/S100 | 3 |
| BP | GO:00064 | protein lipi | 3/151 | 91/18870 | 0.0366145 | 0.6094727 | 0.6090442 | ZDHHC8/ | 3 |
| BP | GO:00073 | acrosome r | 2/151 | 38/18870 | 0.0370585 | 0.6094727 | 0.6090442 | PKDREJ/ | 2 |
| BP | GO:00359 | aorta morp | 2/151 | 38/18870 | 0.0370585 | 0.6094727 | 0.6090442 | TBX2/FG | 2 |
| BP | GO:00605 | neuroepith | 2/151 | 38/18870 | 0.0370585 | 0.6094727 | 0.6090442 | CDH2/FG | 2 |
| BP | GO:00076 | sensory pe | 4/151 | 158/18870 | 0.0382666 | 0.6094727 | 0.6090442 | KCNQ4/C | 4 |
| BP | GO:00219 | central ner | 2/151 | 39/18870 | 0.0388607 | 0.6094727 | 0.6090442 | NR2E1/D | 2 |
| BP | GO:00990 | postsynaps | 2/151 | 39/18870 | 0.0388607 | 0.6094727 | 0.6090442 | ARHGEF9 | 2 |
| BP | GO:01406 | canonical i | 2/151 | 39/18870 | 0.0388607 | 0.6094727 | 0.6090442 | CSNK1A1 | 2 |
| BP | GO:00509 | detection o | 8/151 | 479/18870 | 0.0390487 | 0.6094727 | 0.6090442 | OR2T1/O | 8 |
| BP | GO:00487 | embryonic | 3/151 | 94/18870 | 0.0397146 | 0.6094727 | 0.6090442 | MYF5/AL | 3 |

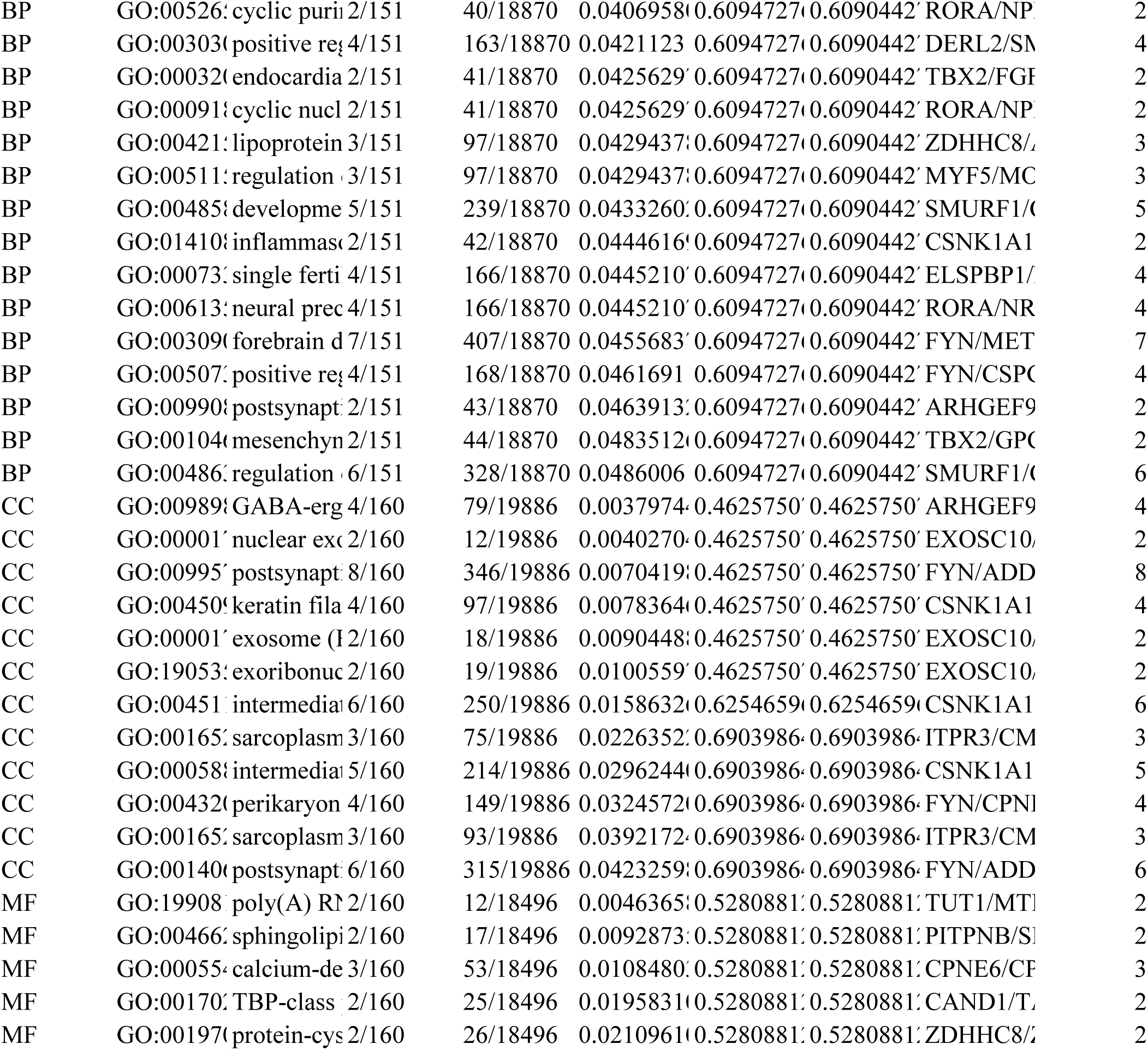

|  |  |  |  |  |  |  |  |  |  |
|---|---|---|---|---|---|---|---|---|---|
| BP | GO:00526 | cyclic puri | 2/151 | 40/18870 | 0.0406958 | 0.6094727 | 0.6090442 | RORA/NP | 2 |
| BP | GO:00303 | positive re | 4/151 | 163/18870 | 0.0421123 | 0.6094727 | 0.6090442 | DERL2/S | 4 |
| BP | GO:00032 | endocardia | 2/151 | 41/18870 | 0.0425629 | 0.6094727 | 0.6090442 | TBX2/FG | 2 |
| BP | GO:00091 | cyclic nucl | 2/151 | 41/18870 | 0.0425629 | 0.6094727 | 0.6090442 | RORA/NP | 2 |
| BP | GO:00421 | lipoprotein | 3/151 | 97/18870 | 0.0429437 | 0.6094727 | 0.6090442 | ZDHHC8/ | 3 |
| BP | GO:00511 | regulation | 3/151 | 97/18870 | 0.0429437 | 0.6094727 | 0.6090442 | MYF5/MC | 3 |
| BP | GO:00485 | developme | 5/151 | 239/18870 | 0.0433260 | 0.6094727 | 0.6090442 | SMURF1/ | 5 |
| BP | GO:01410 | inflammas | 2/151 | 42/18870 | 0.0444616 | 0.6094727 | 0.6090442 | CSNK1A1 | 2 |
| BP | GO:00073 | single ferti | 4/151 | 166/18870 | 0.0445210 | 0.6094727 | 0.6090442 | ELSPBP1/ | 4 |
| BP | GO:00613 | neural prec | 4/151 | 166/18870 | 0.0445210 | 0.6094727 | 0.6090442 | RORA/NR | 4 |
| BP | GO:00309 | forebrain d | 7/151 | 407/18870 | 0.0455683 | 0.6094727 | 0.6090442 | FYN/MET | 7 |
| BP | GO:00507 | positive re | 4/151 | 168/18870 | 0.0461691 | 0.6094727 | 0.6090442 | FYN/CSP | 4 |
| BP | GO:00990 | postsynapt | 2/151 | 43/18870 | 0.0463913 | 0.6094727 | 0.6090442 | ARHGEF9 | 2 |
| BP | GO:00104 | mesenchyn | 2/151 | 44/18870 | 0.0483512 | 0.6094727 | 0.6090442 | TBX2/GP | 2 |
| BP | GO:00486 | regulation | 6/151 | 328/18870 | 0.0486006 | 0.6094727 | 0.6090442 | SMURF1/ | 6 |
| CC | GO:00989 | GABA-erg | 4/160 | 79/19886 | 0.0037974 | 0.4625750 | 0.4625750 | ARHGEF9 | 4 |
| CC | GO:00001 | nuclear ex | 2/160 | 12/19886 | 0.0040270 | 0.4625750 | 0.4625750 | EXOSC10/ | 2 |
| CC | GO:00995 | postsynapt | 8/160 | 346/19886 | 0.0070419 | 0.4625750 | 0.4625750 | FYN/ADD | 8 |
| CC | GO:00450 | keratin fila | 4/160 | 97/19886 | 0.0078364 | 0.4625750 | 0.4625750 | CSNK1A1 | 4 |
| CC | GO:00001 | exosome ( | 2/160 | 18/19886 | 0.0090448 | 0.4625750 | 0.4625750 | EXOSC10/ | 2 |
| CC | GO:19053 | exoribonuc | 2/160 | 19/19886 | 0.0100559 | 0.4625750 | 0.4625750 | EXOSC10/ | 2 |
| CC | GO:00451 | intermedia | 6/160 | 250/19886 | 0.0158632 | 0.6254659 | 0.6254659 | CSNK1A1 | 6 |
| CC | GO:00165 | sarcoplasm | 3/160 | 75/19886 | 0.0226352 | 0.6903986 | 0.6903986 | ITPR3/CM | 3 |
| CC | GO:00058 | intermedia | 5/160 | 214/19886 | 0.0296244 | 0.6903986 | 0.6903986 | CSNK1A1 | 5 |
| CC | GO:00432 | perikaryon | 4/160 | 149/19886 | 0.0324572 | 0.6903986 | 0.6903986 | FYN/CPN | 4 |
| CC | GO:00165 | sarcoplasm | 3/160 | 93/19886 | 0.0392172 | 0.6903986 | 0.6903986 | ITPR3/CM | 3 |
| CC | GO:00140 | postsynapt | 6/160 | 315/19886 | 0.0423259 | 0.6903986 | 0.6903986 | FYN/ADD | 6 |
| MF | GO:19908 | poly(A) R | 2/160 | 12/18496 | 0.0046365 | 0.5280881 | 0.5280881 | TUT1/MT | 2 |
| MF | GO:00466 | sphingolipi | 2/160 | 17/18496 | 0.0092873 | 0.5280881 | 0.5280881 | PITPNB/S | 2 |
| MF | GO:00055 | calcium-de | 3/160 | 53/18496 | 0.0108480 | 0.5280881 | 0.5280881 | CPNE6/C | 3 |
| MF | GO:00170 | TBP-class | 2/160 | 25/18496 | 0.0195831 | 0.5280881 | 0.5280881 | CAND1/T | 2 |
| MF | GO:00197 | protein-cys | 2/160 | 26/18496 | 0.0210961 | 0.5280881 | 0.5280881 | ZDHHC8/ | 2 |

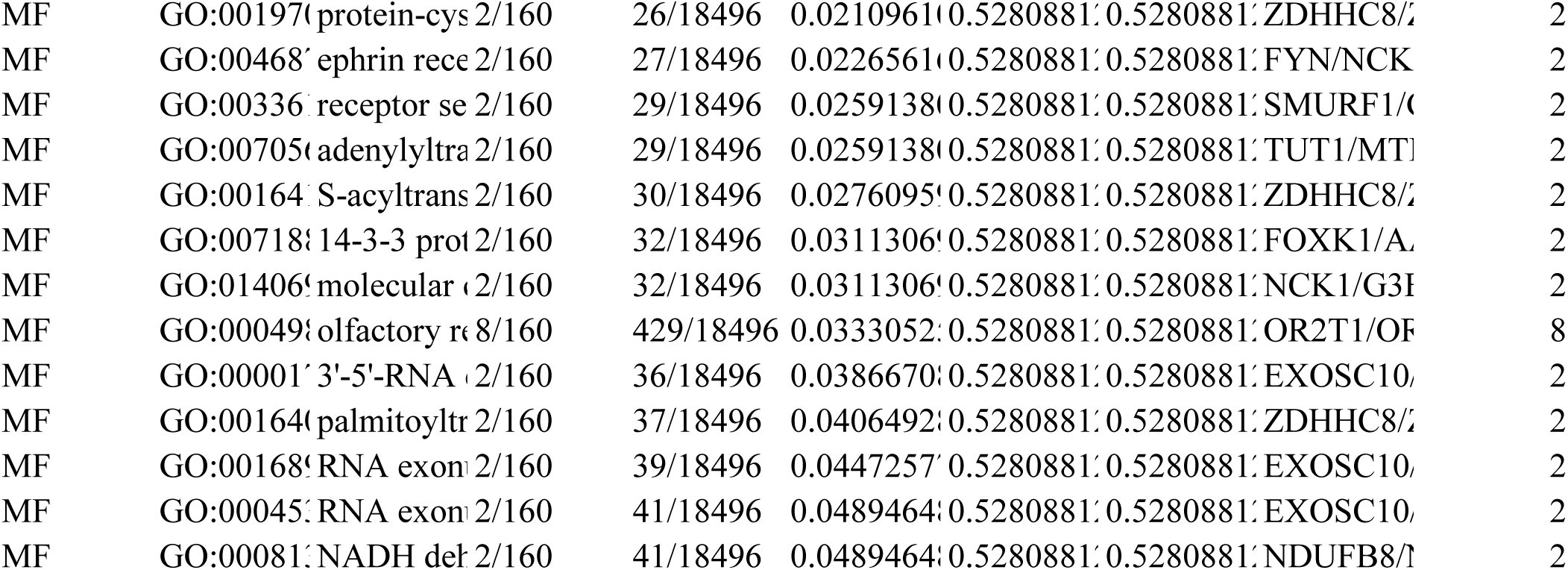

|  |  |  |  |  |  |  |  |  |  |
|---|---|---|---|---|---|---|---|---|---|
| MF | GO:00197 | protein-cys | 2/160 | 26/18496 | 0.0210961 | 0.5280881 | 0.5280881 | ZDHHC8/ | 2 |
| MF | GO:00468 | ephrin rece | 2/160 | 27/18496 | 0.0226561 | 0.5280881 | 0.5280881 | FYN/NCK | 2 |
| MF | GO:00336 | receptor se | 2/160 | 29/18496 | 0.0259138 | 0.5280881 | 0.5280881 | SMURF1/ | 2 |
| MF | GO:00705 | adenylyltra | 2/160 | 29/18496 | 0.0259138 | 0.5280881 | 0.5280881 | TUT1/MT | 2 |
| MF | GO:00164 | S-acyltrans | 2/160 | 30/18496 | 0.0276095 | 0.5280881 | 0.5280881 | ZDHHC8/ | 2 |
| MF | GO:00718 | 14-3-3 prot | 2/160 | 32/18496 | 0.0311306 | 0.5280881 | 0.5280881 | FOXK1/A | 2 |
| MF | GO:01406 | molecular | 2/160 | 32/18496 | 0.0311306 | 0.5280881 | 0.5280881 | NCK1/G3 | 2 |
| MF | GO:00049 | olfactory r | 8/160 | 429/18496 | 0.0333052 | 0.5280881 | 0.5280881 | OR2T1/O | 8 |
| MF | GO:00001 | 3'-5'-RNA | 2/160 | 36/18496 | 0.0386670 | 0.5280881 | 0.5280881 | EXOSC10/ | 2 |
| MF | GO:00164 | palmitoyltr | 2/160 | 37/18496 | 0.0406492 | 0.5280881 | 0.5280881 | ZDHHC8/ | 2 |
| MF | GO:00168 | RNA exon | 2/160 | 39/18496 | 0.0447257 | 0.5280881 | 0.5280881 | EXOSC10/ | 2 |
| MF | GO:00045 | RNA exon | 2/160 | 41/18496 | 0.0489464 | 0.5280881 | 0.5280881 | EXOSC10/ | 2 |
| MF | GO:00081 | NADH del | 2/160 | 41/18496 | 0.0489464 | 0.5280881 | 0.5280881 | NDUFB8/ | 2 |

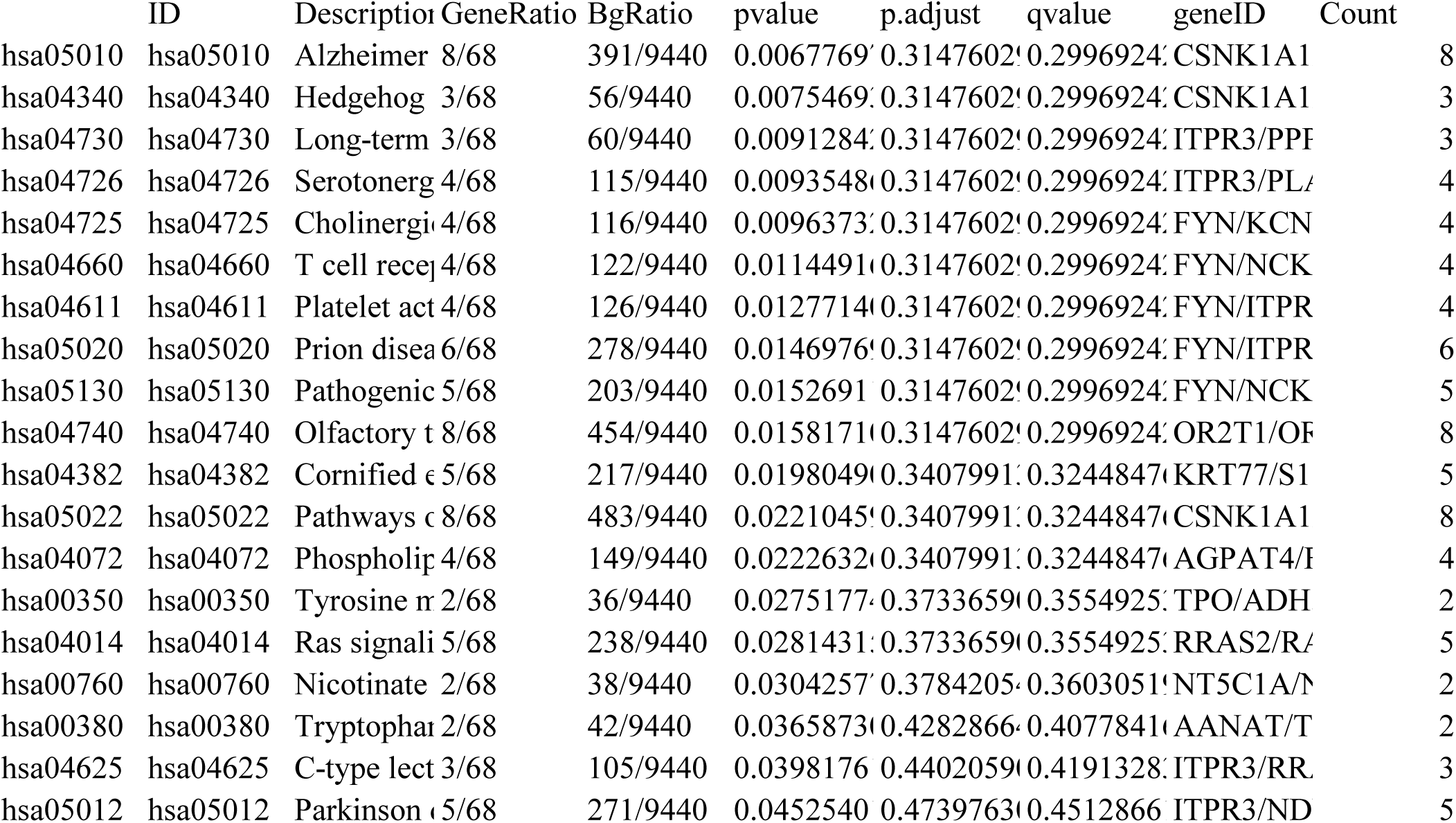

|  | ID | Descriptio | GeneRatio | BgRatio | pvalue | p.adjust | qvalue | geneID | Count |
|---|---|---|---|---|---|---|---|---|---|
| hsa05010 | hsa05010 | Alzheimer | 8/68 | 391/9440 | 0.0067769 | 0.3147602 | 0.2996924 | CSNK1A1 | 8 |
| hsa04340 | hsa04340 | Hedgehog | 3/68 | 56/9440 | 0.0075469 | 0.3147602 | 0.2996924 | CSNK1A1 | 3 |
| hsa04730 | hsa04730 | Long-term | 3/68 | 60/9440 | 0.0091284 | 0.3147602 | 0.2996924 | ITPR3/PP | 3 |
| hsa04726 | hsa04726 | Serotonerg | 4/68 | 115/9440 | 0.0093548 | 0.3147602 | 0.2996924 | ITPR3/PL | 4 |
| hsa04725 | hsa04725 | Cholinergi | 4/68 | 116/9440 | 0.0096373 | 0.3147602 | 0.2996924 | FYN/KCN | 4 |
| hsa04660 | hsa04660 | T cell rece | 4/68 | 122/9440 | 0.0114491 | 0.3147602 | 0.2996924 | FYN/NCK | 4 |
| hsa04611 | hsa04611 | Platelet act | 4/68 | 126/9440 | 0.0127714 | 0.3147602 | 0.2996924 | FYN/ITPR | 4 |
| hsa05020 | hsa05020 | Prion disea | 6/68 | 278/9440 | 0.0146976 | 0.3147602 | 0.2996924 | FYN/ITPR | 6 |
| hsa05130 | hsa05130 | Pathogenic | 5/68 | 203/9440 | 0.0152691 | 0.3147602 | 0.2996924 | FYN/NCK | 5 |
| hsa04740 | hsa04740 | Olfactory t | 8/68 | 454/9440 | 0.0158171 | 0.3147602 | 0.2996924 | OR2T1/O | 8 |
| hsa04382 | hsa04382 | Cornified | 5/68 | 217/9440 | 0.0198049 | 0.3407991 | 0.3244847 | KRT77/S1 | 5 |
| hsa05022 | hsa05022 | Pathways | 8/68 | 483/9440 | 0.0221045 | 0.3407991 | 0.3244847 | CSNK1A1 | 8 |
| hsa04072 | hsa04072 | Phospholi | 4/68 | 149/9440 | 0.0222632 | 0.3407991 | 0.3244847 | AGPAT4/ | 4 |
| hsa00350 | hsa00350 | Tyrosine | 2/68 | 36/9440 | 0.0275177 | 0.3733659 | 0.3554925 | TPO/ADH | 2 |
| hsa04014 | hsa04014 | Ras signali | 5/68 | 238/9440 | 0.0281431 | 0.3733659 | 0.3554925 | RRAS2/R | 5 |
| hsa00760 | hsa00760 | Nicotinate | 2/68 | 38/9440 | 0.0304257 | 0.3784205 | 0.3603051 | NT5C1A/ | 2 |
| hsa00380 | hsa00380 | Tryptopha | 2/68 | 42/9440 | 0.0365873 | 0.4282866 | 0.4077841 | AANAT/T | 2 |
| hsa04625 | hsa04625 | C-type lect | 3/68 | 105/9440 | 0.0398176 | 0.4402059 | 0.4191328 | ITPR3/RR | 3 |
| hsa05012 | hsa05012 | Parkinson | 5/68 | 271/9440 | 0.0452540 | 0.4739763 | 0.4512866 | ITPR3/ND | 5 |

| ID | Descriptio | setSize | enrichment | NES | pvalue | p.adjust | qvalue | rank | leading_ed | core_enrichment |
|---|---|---|---|---|---|---|---|---|---|---|
| KEGG_MI | KEGG_MI | 18 | 0.6898124 | 2.4997770 | 5.8791232 | 0.0021341 | 0.0018256 | 3107 | tags=56%, | PPP2R1B/CTLA4/PPP2R2A/AK |
| KEGG_MI | KEGG_MI | 15 | 0.6961287 | 2.4033257 | 2.1515941 | 0.0025013 | 0.0021397 | 3107 | tags=53%, | PPP2R1B/PPP2R2A/AKT3/PPP2 |
| KEGG_MI | KEGG_MI | 29 | 0.5543296 | 2.2978562 | 2.7563495 | 0.0025013 | 0.0021397 | 3772 | tags=55%, | PHC3/PHF1/RNF2/CTBP2/SUZ1 |
| KEGG_MI | KEGG_MI | 17 | 0.6306572 | 2.2642624 | 2.0016271 | 0.0066053 | 0.0056505 | 4871 | tags=65%, | BCOR/SKP1/RNF2/SUZ12/EZH1 |
| KEGG_MI | KEGG_MI | 21 | 0.5817228 | 2.2114060 | 1.8485571 | 0.0066053 | 0.0056505 | 5309 | tags=67%, | TRADD/MAP3K7/CREB3L1/CR |
| KEGG_MI | KEGG_MI | 22 | 0.5686142 | 2.2002874 | 3.9080134 | 0.0118217 | 0.0101128 | 4545 | tags=59%, | NUP210/NUP205/NUP133/NUP3 |
| KEGG_MI | KEGG_MI | 37 | 0.4998669 | 2.1875528 | 2.5748372 | 0.0025013 | 0.0021397 | 4468 | tags=54%, | NDUFS2/NDUFB8/NDUFB2/ND |
| KEGG_MI | KEGG_MI | 37 | 0.4946167 | 2.1645764 | 3.5521327 | 0.0025788 | 0.0022060 | 4468 | tags=54%, | NDUFS2/NDUFB8/NDUFB2/ND |
| KEGG_MI | KEGG_MI | 36 | 0.4876289 | 2.1325992 | 1.3017810 | 0.0066053 | 0.0056505 | 4468 | tags=53%, | NDUFS2/NDUFB8/NDUFB2/ND |
| KEGG_MI | KEGG_MI | 19 | 0.5633952 | 2.0809260 | 9.8642082 | 0.0275439 | 0.0235622 | 3694 | tags=53%, | TMED10/TMED2/TMED1/ARC |
| KEGG_MI | KEGG_MI | 11 | 0.6641928 | 2.0722142 | 0.0016537 | 0.0414941 | 0.0354959 | 4203 | tags=73%, | CREBBP/CREB3L1/PCNA/CRE |
| KEGG_MI | KEGG_MI | 37 | 0.4685547 | 2.0505224 | 1.7243183 | 0.0066053 | 0.0056505 | 4468 | tags=51%, | NDUFS2/NDUFB8/NDUFB2/ND |
| KEGG_MI | KEGG_MI | 43 | 0.4512144 | 2.0322269 | 1.4747793 | 0.0066053 | 0.0056505 | 4468 | tags=49%, | NDUFS2/NDUFB8/NDUFB2/ND |
| KEGG_MI | KEGG_MI | 46 | 0.4346672 | 2.0137989 | 1.6746638 | 0.0066053 | 0.0056505 | 6111 | tags=59%, | NDUFS2/NDUFB8/COX15/NDU |
| KEGG_MI | KEGG_MI | 15 | 0.5684114 | 1.9623927 | 0.0021898 | 0.0496829 | 0.0425010 | 3902 | tags=47%, | FUT8/MAN2A1/B4GALT3/MAN |
| KEGG_MI | KEGG_MI | 24 | 0.4873666 | 1.9328929 | 0.0017146 | 0.0414941 | 0.0354959 | 4315 | tags=50%, | MGA/RNF2/SUZ12/EZH1/EZH2 |

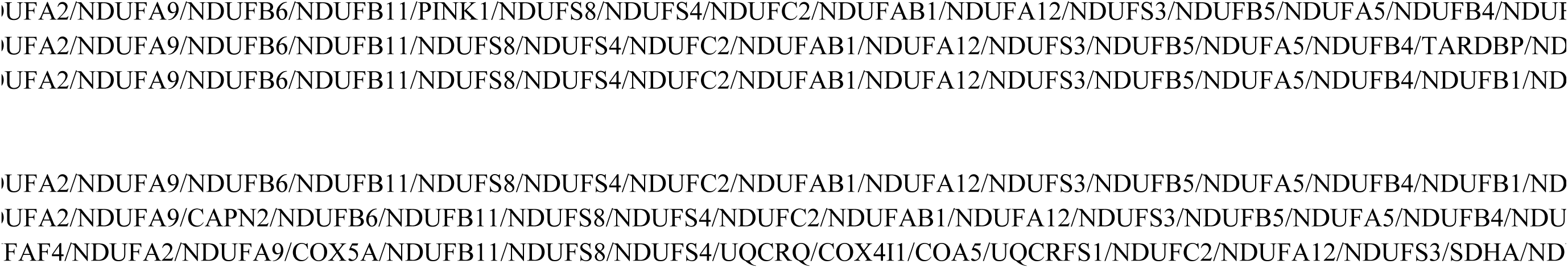

|  |
|---|
| UFA6/COX10/NDUFAF2/NDUFV1/NDUFB7/NDUFA10/NDUFV2/NDUFS1/NDUFAF1 |

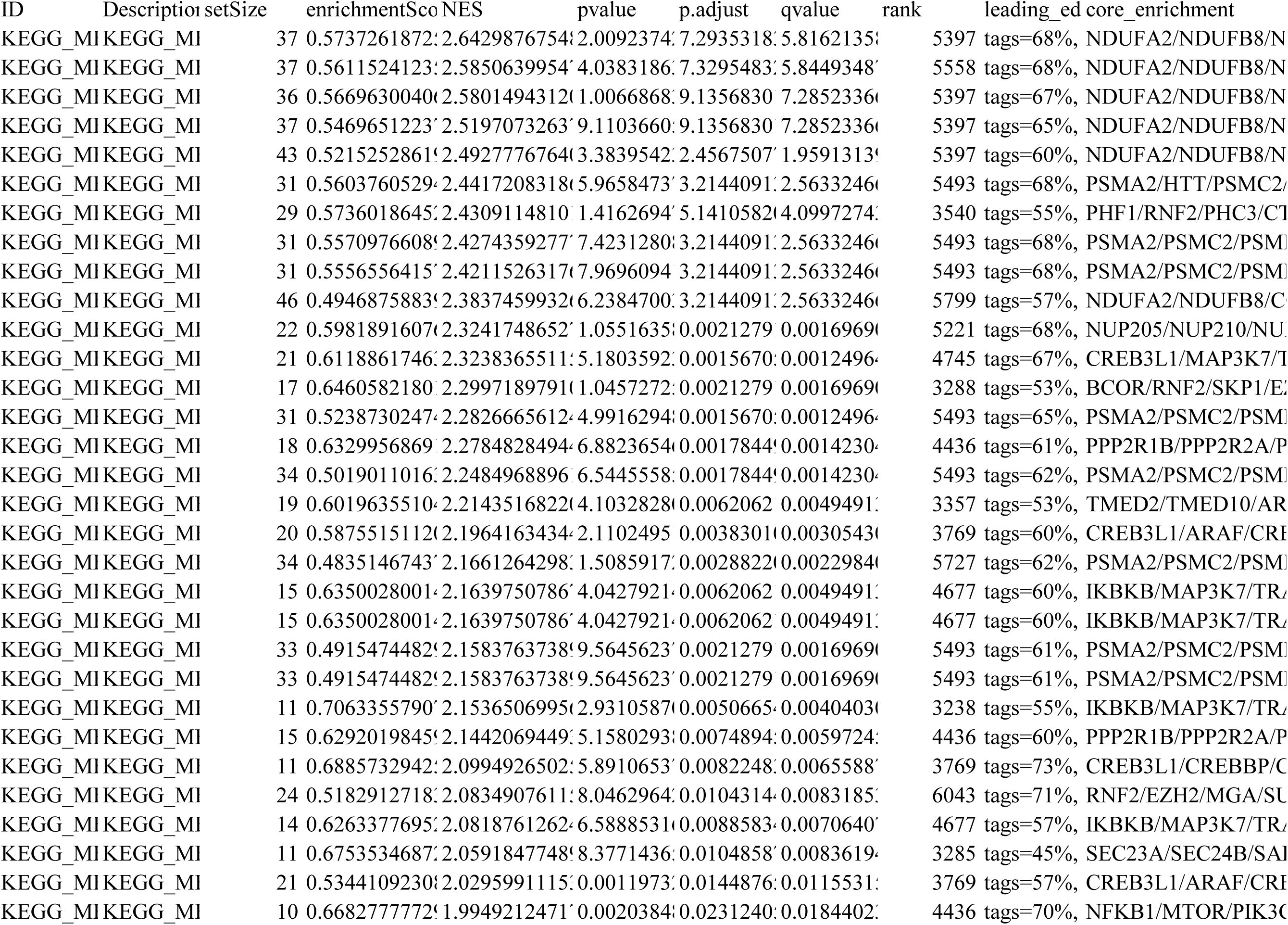

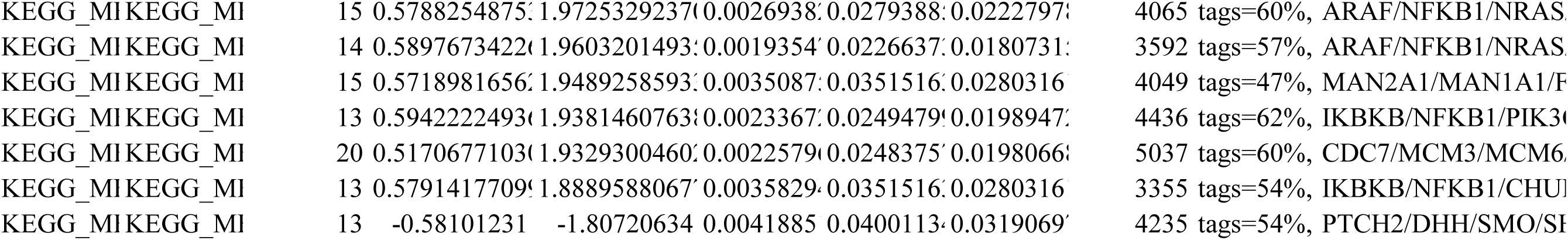

|  |  |  |  |  |  |  |  |  |  |  |
|---|---|---|---|---|---|---|---|---|---|---|
| KEGG_MI | KEGG_MI | 15 | 0.5788254875: | 1.9725329237( | 0.0026938: | 0.0279388: | 0.0222797: | 4065 | tags=60%, | ARAF/NFKB1/NRAS |
| KEGG_MI | KEGG_MI | 14 | 0.5897673422( | 1.9603201493: | 0.0019354' | 0.0226637: | 0.0180731: | 3592 | tags=57%, | ARAF/NFKB1/NRAS |
| KEGG_MI | KEGG_MI | 15 | 0.5718981656: | 1.9489258593: | 0.0035087: | 0.0351516: | 0.0280316 | 4049 | tags=47%, | MAN2A1/MAN1A1/F |
| KEGG_MI | KEGG_MI | 13 | 0.5942222493( | 1.9381460763: | 0.0023367: | 0.0249479! | 0.0198947: | 4436 | tags=62%, | IKBKB/NFKB1/PIK3( |
| KEGG_MI | KEGG_MI | 20 | 0.5170677103( | 1.9329300460: | 0.0022579( | 0.0248375' | 0.0198066: | 5037 | tags=60%, | CDC7/MCM3/MCM6 |
| KEGG_MI | KEGG_MI | 13 | 0.5791417709! | 1.8889588067' | 0.0035829: | 0.0351516: | 0.0280316 | 3355 | tags=54%, | IKBKB/NFKB1/CHUI |
| KEGG_MI | KEGG_MI | 13 | -0.58101231 | -1.80720634 | 0.0041885 | 0.0400113: | 0.0319069' | 4235 | tags=54%, | PTCH2/DHH/SMO/SI |

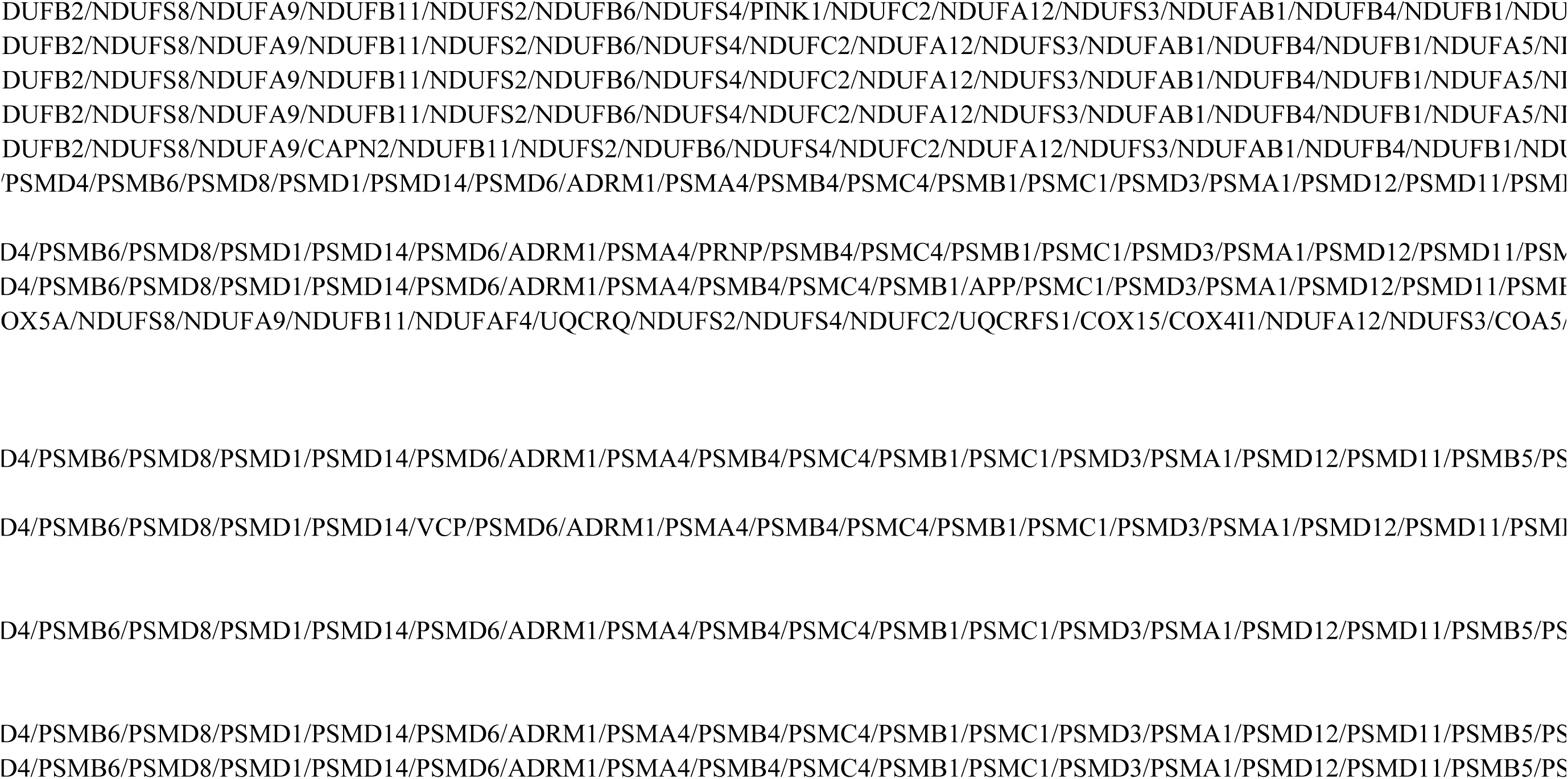

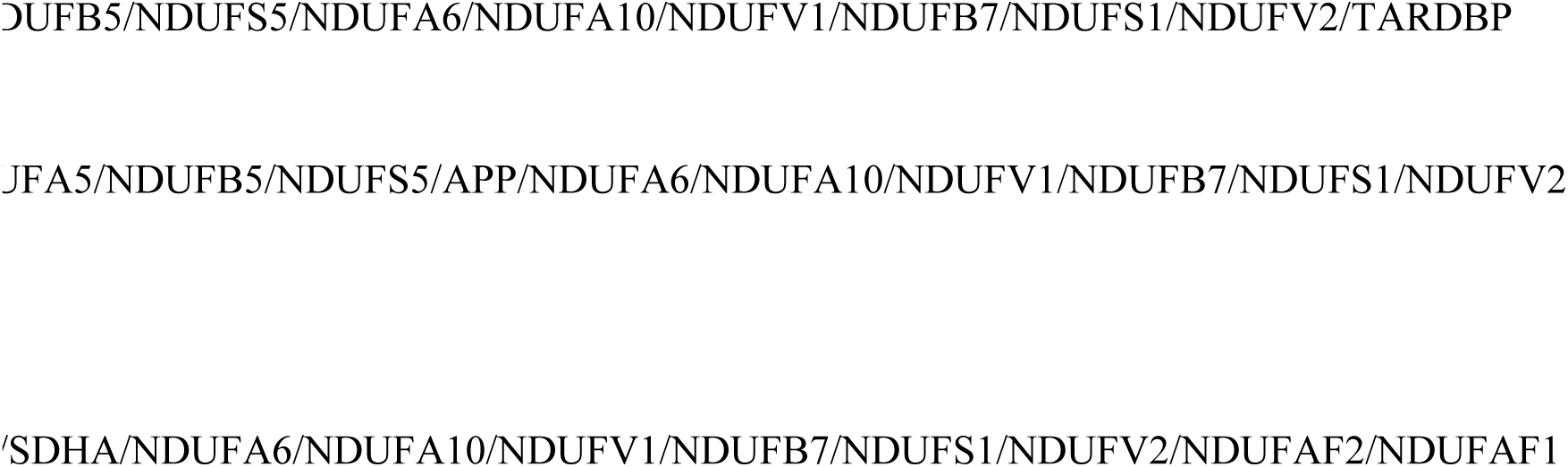

### Supplementary Tables

Tables S1. Curated ARGs.

Tables S2. GO enrichment results for candidate genes.

Tables S3. KEGG pathway enrichment for candidate genes.

Tables S4. GSEA results for DYRK2.

Tables S5. GSEA results for SIGIRR.

## Notes

### Competing Interest Statement

The authors have declared no competing interest.

## References

1. Wang HH, Garruti G, Liu M, Portincasa P, Wang DQ. Cholesterol and Lipoprotein Metabolism and Atherosclerosis: Recent Advances In reverse Cholesterol Transport. Annals of hepatology. 2017;16:s27–s42. doi: 10.5604/01.3001.0010.5495

2. Zhang Y, Wernly B, Cao X, Mustafa SJ, Tang Y, Zhou Z. Adenosine and adenosine receptor-mediated action in coronary microcirculation. Basic Res Cardiol. 2021;116:22. doi: 10.1007/s00395-021-00859-7

3. Marchi E, Muraca I, Berteotti M, Gori AM, Valenti R, Marcucci R. Adenosine in Interventional Cardiology: Physiopathologic and Pharmacologic Effects in Coronary Artery Disease. International Journal of Molecular Sciences. 2024;25. doi: 10.3390/ijms25115852

4. Franco R, Lillo A, Navarro G, Reyes-Resina I. The adenosine A2A receptor is a therapeutic target in neurological, heart and oncogenic diseases. Expert Opinion on Therapeutic Targets. 2022;26:791–800. doi: 10.1080/14728222.2022.2136570

5. Liang W, Zhou C, Wang J, Zhao J, Liu F, Wang G, Xu C, Zhang Y, Wang W, Cai S, et al. A prognostic signature based on adenosine metabolism related genes for ovarian cancer. Frontiers in oncology. 2022;12:1003512. doi: 10.3389/fonc.2022.1003512

6. Ritchie ME, Phipson B, Wu D, Hu Y, Law CW, Shi W, Smyth GK. limma powers differential expression analyses for RNA-sequencing and microarray studies. Nucleic acids research. 2015;43:e47. doi: 10.1093/nar/gkv007

7. Ito K, Murphy D. Application of ggplot2 to Pharmacometric Graphics. CPT: pharmacometrics & systems pharmacology. 2013;2:e79. doi: 10.1038/psp.2013.56

8. Gu Z. Complex heatmap visualization. iMeta. 2022;1:e43. doi: 10.1002/imt2.43

9. Hänzelmann S, Castelo R, Guinney J. GSVA: gene set variation analysis for microarray and RNA-seq data. BMC bioinformatics. 2013;14:7. doi: 10.1186/1471-2105-14-7

10. Langfelder P, Horvath S. WGCNA: an R package for weighted correlation network analysis. BMC bioinformatics. 2008;9:559. doi: 10.1186/1471-2105-9-559

11. Zhang X, Chao P, Zhang L, Xu L, Cui X, Wang S, Wusiman M, Jiang H, Lu C. Single-cell RNA and transcriptome sequencing profiles identify immune-associated key genes in the development of diabetic kidney disease. Frontiers in immunology. 2023;14:1030198. doi: 10.3389/fimmu.2023.1030198

12. Li Y, Chen C, Diao M, Wei Y, Zhu Y, Hu W. Gene model-related m6A expression levels predict the risk of preeclampsia. BMC medical genomics. 2022;15:103. doi: 10.1186/s12920-022-01254-4

13. Zhou W, Li H, Zhang J, Liu C, Liu D, Chen X, Ouyang J, Zeng T, Peng S, Ouyang F, et al. Identification and mechanism analysis of biomarkers related to butyrate metabolism in COVID-19 patients. Ann Med. 2025;57:2477301. doi: 10.1080/07853890.2025.2477301

14. Wu T, Hu E, Xu S, Chen M, Guo P, Dai Z, Feng T, Zhou L, Tang W, Zhan L, et al. clusterProfiler 4.0: A universal enrichment tool for interpreting omics data. Innovation (Cambridge (Mass)). 2021;2:100141. doi: 10.1016/j.xinn.2021.100141

15. Gu Z, Gu L, Eils R, Schlesner M, Brors B. circlize Implements and enhances circular visualization in R. *Bioinformatics (Oxford*, England). 2014;30:2811–2812. doi: 10.1093/bioinformatics/btu393

16. Hou N, Li M, He L, Xie B, Wang L, Zhang R, Yu Y, Sun X, Pan Z, Wang K. Predicting 30-days mortality for MIMIC-III patients with sepsis-3: a machine learning approach using XGboost. J Transl Med. 2020;18:462. doi: 10.1186/s12967-020-02620-5

17. Zhang Z, Zhao Y, Canes A, Steinberg D, Lyashevska O. Predictive analytics with gradient boosting in clinical medicine. Ann Transl Med. 2019;7:152. doi: 10.21037/atm.2019.03.29

18. Zhou H, Xin Y, Li S. A diabetes prediction model based on Boruta feature selection and ensemble learning. BMC bioinformatics. 2023;24:224. doi: 10.1186/s12859-023-05300-5

19. Robin X, Turck N, Hainard A, Tiberti N, Lisacek F, Sanchez JC, Müller M. pROC: an open-source package for R and S+ to analyze and compare ROC curves. BMC bioinformatics. 2011;12:77. doi: 10.1186/1471-2105-12-77

20. Wang X, Zhang Y. Multi-omics joint screening of biomarkers related to M2 macrophages in gastric cancer. Discover oncology. 2024;15:738. doi: 10.1007/s12672-024-01623-8

21. Chen Y, He J, Pan X. Prediction of risk factors for preoperative deep vein thrombosis in patients with pelvic fracture. Frontiers in surgery. 2025;12:1585460. doi: 10.3389/fsurg.2025.1585460

22. Orifjon S, Jammatov J, Sousa C, Barros R, Vasconcelos O, Rodrigues P. Translation and Adaptation of the Adult Developmental Coordination Disorder/Dyspraxia Checklist (ADC) into Asian Uzbekistan. Sports (Basel). 2023;11. doi: 10.3390/sports11070135

23. Lu H, Xu Z, Shao L, Li P, Xia Y. High infiltration of immune cells with lower immune activity mediated the heterogeneity of gastric adenocarcinoma and promoted metastasis. Heliyon. 2024;10:e37092. doi: 10.1016/j.heliyon.2024.e37092

24. Zhang Y, Miao Y, Tan J, Chen F, Lei P, Zhang Q. Identification of mitochondrial related signature associated with immune microenvironment in Alzheimer’s disease. Journal of translational medicine. 2023;21:458. doi: 10.1186/s12967-023-04254-9

25. Zhang H, Meltzer P, Davis S. RCircos: an R package for Circos 2D track plots. BMC bioinformatics. 2013;14:244. doi: 10.1186/1471-2105-14-244

26. Ladias P, Markopoulos GS, Kostoulas C, Bouba I, Markoula S, Georgiou I. Cancer Associated PRDM9: Implications for Linking Genomic Instability and Meiotic Recombination. Int J Mol Sci. 2023;24. doi: 10.3390/ijms242216522

27. Sato N, Uematsu M, Fujimoto K, Uematsu S, Imoto S. ggkegg: analysis and visualization of KEGG data utilizing the grammar of graphics. *Bioinformatics (Oxford*, England). 2023;39. doi: 10.1093/bioinformatics/btad622

28. Satija R, Farrell JA, Gennert D, Schier AF, Regev A. Spatial reconstruction of single-cell gene expression data. Nature biotechnology. 2015;33:495–502. doi: 10.1038/nbt.3192

29. Zhu L, Wang W, Ren C, Wang Y, Zhang G, Liu J, Wang W. Cellular Phenotypic Transformation in Heart Failure Caused by Coronary Heart Disease and Dilated Cardiomyopathy: Delineating at Single-Cell Level. Biomedicines. 2022;10. doi: 10.3390/biomedicines10020402

30. Qiu X, Hill A, Packer J, Lin D, Ma YA, Trapnell C. Single-cell mRNA quantification and differential analysis with Census. Nature methods. 2017;14:309–315. doi: 10.1038/nmeth.4150

31. Grentner A, Ragueneau E, Gong C, Prinz A, Gansberger S, Oyarzun I, Hermjakob H, Griss J. ReactomeGSA: new features to simplify public data reuse. *Bioinformatics (Oxford*, England). 2024;40. doi: 10.1093/bioinformatics/btae338

32. Cao L, Lu X, Wang X, Wu H, Miao X. From single-cell to spatial transcriptomics: decoding the glioma stem cell niche and its clinical implications. Frontiers in immunology. 2024;15:1475235. doi: 10.3389/fimmu.2024.1475235

33. Chang J, Wu H, Wu J, Liu M, Zhang W, Hu Y, Zhang X, Xu J, Li L, Yu P, et al. Constructing a novel mitochondrial-related gene signature for evaluating the tumor immune microenvironment and predicting survival in stomach adenocarcinoma. Journal of translational medicine. 2023;21:191. doi: 10.1186/s12967-023-04033-6

34. Hori M, Kitakaze M. Adenosine, the heart, and coronary circulation. Hypertension (Dallas, Tex : 1979). 1991;18:565–574. doi: 10.1161/01.hyp.18.5.565

35. Ogilby JD, Heo J, Iskandrian AS. Effect of adenosine on coronary blood flow and its use as a diagnostic test for coronary artery disease. Cardiovascular research. 1993;27:48–53. doi: 10.1093/cvr/27.1.48

36. Correa-Sáez A, Jiménez-Izquierdo R, Garrido-Rodríguez M, Morrugares R, Muñoz E, Calzado MA. Updating dual-specificity tyrosine-phosphorylation-regulated kinase 2 (DYRK2): molecular basis, functions and role in diseases. Cellular and molecular life sciences : CMLS. 2020;77:4747–4763. doi: 10.1007/s00018-020-03556-1

37. Tandon V, de la Vega L, Banerjee S. Emerging roles of DYRK2 in cancer. The Journal of biological chemistry. 2021;296:100233. doi: 10.1074/jbc.REV120.015217

38. Kawamura A, Yoshida S, Yoshida K. The diverse functions of DYRK2 in response to cellular stress. Histology and histopathology. 2024;39:1427–1434. doi: 10.14670/hh-18-744

39. Yoshida S, Kawamura A, Aoki K, Wiriyasermkul P, Sugimoto S, Tomiyoshi J, Tajima A, Ishida Y, Katoh Y, Tsukada T, et al. Positive regulation of Hedgehog signaling via phosphorylation of GLI2/GLI3 by DYRK2 kinase. Proceedings of the National Academy of Sciences of the United States of America. 2024;121:e2320070121. doi: 10.1073/pnas.2320070121

40. Santos-Durán GN, Barreiro-Iglesias A. Roles of dual specificity tyrosine-phosphorylation-regulated kinase 2 in nervous system development and disease. Frontiers in neuroscience. 2022;16:994256. doi: 10.3389/fnins.2022.994256

41. Cheng S, Cui L, Chen J, Xiu Q, Si R, Yang X, Shi Y. SIGIRR: An Orphan Receptor Mediating Anti-inflammatory Actions. Expert reviews in molecular medicine. 2025;27:e24. doi: 10.1017/erm.2025.10009

42. Tian F, Lei J, Ni Y, Zhong D, Xie N, Ma J, Wang H, Si S, Wu Y, Jiang T. Regulation of CD18 stability by SIGIRR-modulated ubiquitination: new insights into the relationship between innate immune response and acute lung injury. The FEBS journal. 2023;290:2721–2743. doi: 10.1111/febs.16708

43. Su Z, Tao X. Current Understanding of IL-37 in Human Health and Disease. Frontiers in immunology. 2021;12:696605. doi: 10.3389/fimmu.2021.696605

44. Mohanta SK, Heron C, Klaus-Bergmann A, Horstmann H, Brakenhielm E, Giannarelli C, Habenicht AJR, Gerhardt H, Weber C. Metabolic and Immune Crosstalk in Cardiovascular Disease. Circulation research. 2025;136:1433–1453. doi: 10.1161/circresaha.125.325496

45. Kubota A, Frangogiannis NG. Macrophages in myocardial infarction. American journal of physiology Cell physiology. 2022;323:C1304–c1324. doi: 10.1152/ajpcell.00230.2022

46. Akhtar S, Sagar K, Roy A, Hote MP, Arava S, Sharma A. CCR5-mediated homing of regulatory T cells and monocytic-myeloid derived suppressor cells to dysfunctional endothelium contributes to early atherosclerosis. Immunology. 2024;173:712–729. doi: 10.1111/imm.13859

47. Miano JM, Fisher EA, Majesky MW. Fate and State of Vascular Smooth Muscle Cells in Atherosclerosis. Circulation. 2021;143:2110–2116. doi: 10.1161/circulationaha.120.049922

48. Connor SM, Rashid M, Ryan KJ, Patel K, Boyd JD, Smith J, Elyaman W, Bennett DA, Bradshaw EM. GW5074 Increases Microglial Phagocytic Activities: Potential Therapeutic Direction for Alzheimer’s Disease. Frontiers in cellular neuroscience. 2022;16:894601. doi: 10.3389/fncel.2022.894601

49. Sarkar R, Choudhury SM, Kanneganti TD. Classical apoptotic stimulus, staurosporine, induces lytic inflammatory cell death, PANoptosis. The Journal of biological chemistry. 2024;300:107676. doi: 10.1016/j.jbc.2024.107676

